# Microbial dynamics inference at ecosystem-scale

**DOI:** 10.1101/2021.12.14.469105

**Authors:** Travis E. Gibson, Younhun Kim, Sawal Acharya, David E. Kaplan, Nicholas DiBenedetto, Richard Lavin, Bonnie Berger, Jessica R. Allegretti, Lynn Bry, Georg K. Gerber

## Abstract

Dynamical systems models are a powerful tool for analyzing interactions, stability, resilience, and other key properties in biomedically important microbial ecosystems, such as the gut microbiome. Challenges to modeling and inference in this setting include the large number of species present, and data sparsity/noise characteristics. Here, we introduce a Bayesian statistical method, the Microbial Dynamical Systems Inference Engine 2 (MDSINE2), which infers compact and interpretable ecosystems-scale dynamical systems models from microbiome time-series data. We model microbial dynamics as stochastic processes driven by inferred interaction modules, or groups of microbes with similar interaction structure and responses to perturbations. Additionally, we model the noise characteristics of sequencing and qPCR measurements to provide uncertainty quantification for all outputs. To evaluate MDSINE2, and provide a benchmarking resource for the community, we generated the most densely sampled microbiome time-series to date, which consists of a cohort of mice that received fecal transplants from a human donor and were then subjected to dietary and antibiotic perturbations. Benchmarking on simulated and real data demonstrate that MDSINE2 significantly outperforms state-of-the-art methods, and moreover identifies interaction modules that shed new light on ecosystems-scale interactions in the gut microbiome. We provide MDSINE2 as an open-source Python package at: https://github.com/gerberlab/MDSINE2.

## Introduction

Microbiomes are inherently dynamic^2^, changing over time due to both microbial interactions, as well as responses to external perturbations. Dynamics of a microbiome reveals important information not only about the individual microbial constituents, but also about how the ecosystem as a whole behaves; for instance, unstable responses of the ecosystem to perturbations can be indicative of an inability to maintain homeostatic function^3^. Mathematical models of dynamical systems have a long history in ecology and biomedicine, and have led to many insights, including for microbial ecosystems^4^. Dynamical systems models are particularly powerful because, once inferred from data, they can be directly interrogated using mathematical tools or computational simulations to study aspects including: stability and other ecological properties^5–8^; topological properties of the interaction network such as motifs^9–11^; and *in silico* forecasts of the system such as “knock-outs” of taxa or responses to perturbations not yet experimentally studied.

However, the scale and complexity of microbiomes, as well as limitations of measurement modalities, present challenges to modeling and inferring dynamical systems models from data. The gut microbiome contains hundreds of ecologically diverse yet interacting microbes. Indeed, there is increasing recognition that this interaction structure is critically important, driving behaviors such as whether so-called pathobiont bacteria will cause disease or remain harmless in the host^12^. A well-established modeling framework, which we and others have employed, uses the generalized Lotka-Volterra (gLV) equations^1, 13–15^ to model pair-wise interactions among microbial taxa. Although gLV models have been shown empirically to predict microbial dynamics with good accuracy for small ecosystems^1^, these models present significant challenges for scalability and interpretability, because the number of modeled interactions increases quadratically with the number of taxa in the system. This large number of interaction parameters must be inferred from microbiome data, which itself has fundamental limitations. The frequency and regularity of sampling is dependent on gut transit time and practical logistics, particularly for human studies. Further, microbiome data relies on sequencing and other high-throughput methods, which have complicated noise characteristics. Measurement noise presents particular challenges when we seek to characterize low abundance components of the microbiome, which can serve critical ecological roles^16, 17^, but are orders of magnitude lower than the predominant taxa in the gut. Sufficiently rich experimental data is also critical for model inference^18^. Perturbations are essential to analyze how components of the system interact and to assess stability and other key properties; data at equilibrium in a single or small number of conditions cannot be used to infer these properties of the system.

Here, we provide two new resources to the community to address the challenge of inferring microbial dynamics at ecosystem-scale: (1) a fully Bayesian computational method, the Microbial Dynamical Systems Inference Engine 2 (MDSINE2), implemented as an open-source software package, and (2) a densely sampled gut microbiome time-series dataset for benchmarking and other analyses. MDSINE2 has several advantages over state-of-the-art methods, including our own prior contributions^1, 13–15^. Our method is the first to both explicitly model measurement noise and fully propagate this information during inference, which has distinct advantages for quantifying uncertainty of predictions and more accurately capturing behavior of low abundance species in the ecosystem. Further, MDSINE2 automatically learns interaction modules, or groups of bacteria with similar interaction structure and responses to perturbations, which allows our model to scale to large ecosystems while maintaining interpretability. We make MDSINE2 available to the community as a fully open-source Python package, and also provide extensive Colab tutorials. To provide a community resource for benchmarking MDSINE2 and other methods, we generated, to our knowledge, the most temporally densely sampled gut microbiome dataset to date that includes perturbations essential for dynamical systems inference. For this dataset, we developed a cohort of “humanized” gnotobiotic mice via fecal transplantation from a healthy human donor and then subjected mice to series of dietary and antibiotic perturbations, while collecting an average of 77 fecal samples per mouse over 65 days. Samples were then analyzed with both high-throughput 16S rRNA gene amplicon sequencing to estimate microbial relative abundances and 16S rRNA gene qPCR to quantitate total microbial concentrations, with physical replicates included to calibrate computational noise models.

## Results

### MDSINE2 is an open-source computational tool for inferring dynamical systems models of microbiomes at scale

To infer accurate and interpretable large-scale dynamical systems models from microbiome time-series data, we developed MDSINE2 (Figure 1), a fully Bayesian machine learning model. Inference is performed using a custom Markov Chain Monte Carlo (MCMC) algorithm to approximate the posterior probability, which is implemented in a software package that leverages vectorization and just-in-time compilation technologies for high efficiency. Inputs to the MDSINE2 software are: (1) time-series measurements of bacterial abundances in the form of counts (e.g., 16S rRNA gene amplicon or shotgun metagenomics data), (2) total bacterial concentrations (e.g., from 16S rRNA gene qPCR measurements), and (3) associated metadata for the samples (Figure 1A). The software also provides a variety of tools for interrogating trajectories of taxa (including under perturbations or with removal of taxa from the system), analyzing topological properties of the interaction network, quantitating the keystoneness of individual taxa or modules, and formally assessing the stability of the microbial ecosystem (Figure 1E,F,G). The core MDSINE2 software is maintained in the Gerber Lab repo https://github.com/gerberlab/MDSINE2 while the code to reproduce the figures for this paper along with Google Colab notebooks for exploring the model are maintained at https://github.com/gerberlab/MDSINE2_Paper.

**Figure 1.**
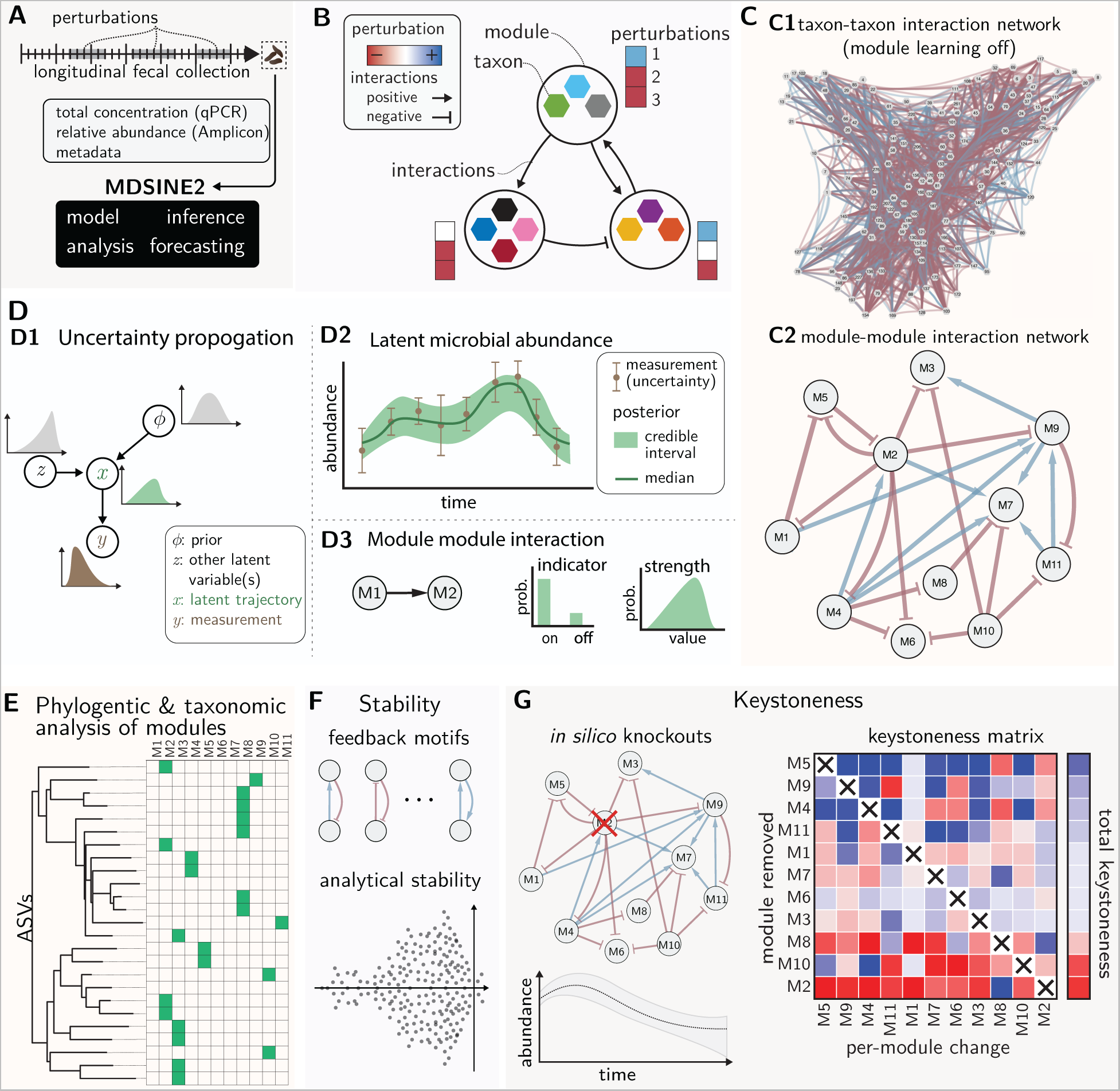
Schematic of the MDSINE2 method for inferring interpretable dynamical systems models of microbiomes at scale. **(A)** Input data are measurements of total bacterial concentration (e.g., 16S rRNA gene qPCR) and measurements of taxa abundances (e.g., 16S rRNA gene amplicon sequencing). Measurements are obtained from studies in which the microbiome undergoes perturbations, providing sufficiently rich information for inference. **(B)** MDSINE2 infers dynamical systems models from data with the option of automatically learning interaction modules, or groups of taxa that share the same interactions with other modules and perturbations. This is a more compact representation that is more readily interpretable than learning interactions among all microbes. **(C)** Example microbial interaction networks for the same number of taxa without module learning (C1) and with module learning (C2)**. (D)** MDSINE is fully Bayesian and propagates error throughout the model (D1), providing estimates of uncertainty for all variables, (e.g. (D2) latent trajectory along with measurements and their uncertainty, (D3) indicator and interaction strength for ecological interactions. The software provides a variety of tools for analysis and visualization of the inferred dynamical system, including: **(E)** analyses of taxonomic composition and phylogeny of modules, **(F)** formal analyses of ecosystem stability and interaction motifs, and **(G)** keystoneness (quantitative impact on the ecosystem when modules are removed).

Our computational model is based on generalized Lotka-Volterra dynamics, but with several key innovations compared to prior state of the art^1, 13–15^. First, MDSINE2 explicitly models the measurement uncertainty associated with microbiome sequencing and qPCR measurements. Unlike our own previous method^1^, which handled data denoising and dynamical systems inference in two separate steps, MDSINE2 fully propagates uncertainty through the model (Figure 1D) and provides quantitative measures (e.g., Bayes factors) of uncertainty for all model parameters. Second, MDSINE2 includes stochastic effects in dynamics, which capture non-deterministic fluctuations in microbial trajectories that occur due to unmeasured effects on the ecosystem. Third, MDSINE2 extends the gLV model to include automatically learn *interaction modules* (Figure 1B), which we define as groups of taxa that share common interaction structure (i.e., are promoted or inhibited by the same taxa outside the module) and have a common response to external perturbations (e.g., antibiotics). Interaction modules are motivated by both empirical observations that groups of microbial taxa covary^19, 20^ and theoretical ecology concepts such as guilds, or groups of taxa that utilize resources in a similar way^21^. Modular structure (Figures 1B and 1C2) reduces the complexity of the system to be analyzed, which has the potential to increase human interpretability^22, 23^. Further, scalability is enabled: the number of parameters in the model is reduced from order quadratic in the number of taxa (e.g., all potential pairwise interactions between taxa in the gLV equations) to order quadratic in the number of modules (which scales logarithmically with the number of taxa)^24^. The number of modules is treated probabilistically with full uncertainty quantification, using a nonparametric Bayesian framework, and learned from the data, alleviating the need for the user to pre-specify this information. See Methods §1 and Supplemental Text §1 for details on the model and Methods §4.1 and Supplemental Text §2 for details on inference.

### MDSINE2 outperforms state-of-the-art models on existing in silico benchmark datasets

To provide an initial point of comparison, we assessed MDSINE2 against our prior method, as well as two state-of-the-art gLV-based methods: ridge regression (gLV-L2) and elastic net regression (gLV-net)^13^, using an *in silico* benchmarking standard we previously published^1^ (Figure 2). Briefly, the dataset simulates gLV dynamics for a small ecosystem of 10 taxa with parameters resembling those learned on real experimental data (Figure 2A and Methods §4.3). We note that we did not compare against methods that use relative abundance as input, because these methods cannot estimate ground truth growth rates or interaction strength, requiring one to provide a rescaling factor (which is unidentifiable when trained on relative abundance data). MDSINE2 significantly outperformed the comparator methods, including our own prior method, on all metrics: inference of growth rates, interaction strengths, and the ground-truth network structure (Figure 2B,C,D and Supplemental Table 1). These results suggest that the new modeling capabilities in MDSINE2 provide significant performance boosts on critical dynamical systems inference tasks. However, this *in silico* dataset simulates only a small ecosystem, and moreover is not expected to fully recapitulate characteristics of real data. Thus, we next sought to generate a new experimental dataset for fully benchmarking the methods and to provide insights into the dynamics of real, large-scale microbial ecosystems.

**Figure 2:**
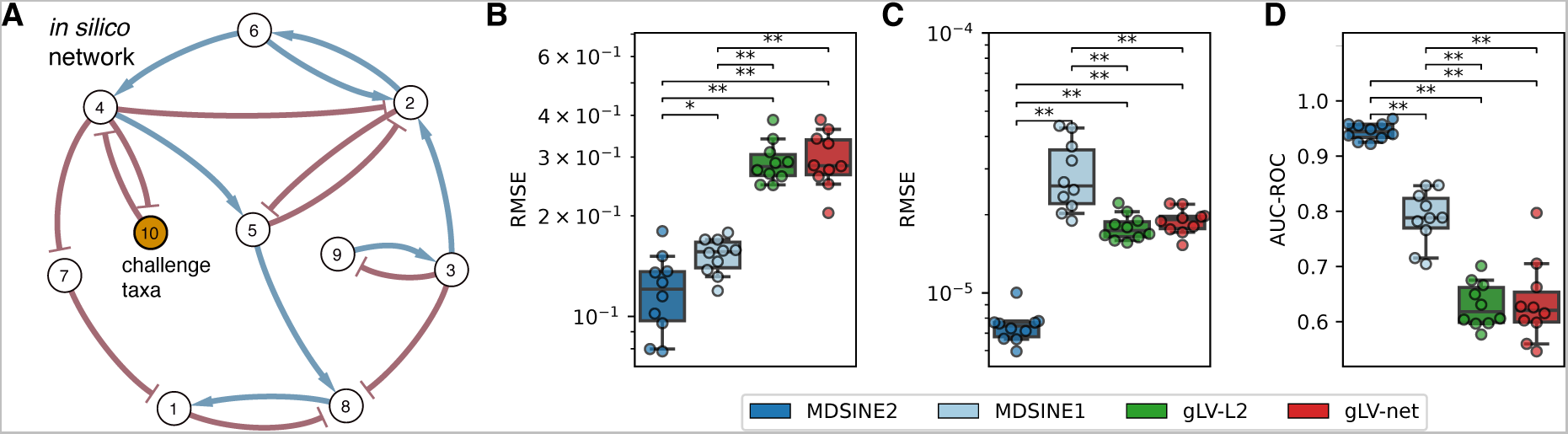
MDSINE2 outperforms state-of-the-art methods on the a synthetic data benchmarking standard. **(A)** Underlying dynamical systems network topology^1^, which was used to simulate data under a gLV model. Root mean-squared error (RMSE) for **(B)** growth rates (lower is better) **(C)** interaction strengths (lower is better), and **(D)** Area under the receiver operating curve (AUC-ROC) for network topology (higher is better, 1 is maximum). * p<0.05, ** p<0.01.

### High-temporal resolution gut microbiome study with serial perturbations is a resource for microbial dynamical systems inference benchmarking and analysis

Given the importance of sufficient temporal resolution and perturbation information for inferring dynamical systems models from data^18^, and the dearth of appropriate datasets available to researchers, we created a new dataset to serve as a benchmarking and analysis resource for the community. To generate the dataset, we created a cohort of “humanized” mice (Figure 3A) by performing a fecal microbiota transplantation from a healthy human donor into germ-free mice (n=4). Gnotobiotic mice, in which bacteria-free mice are inoculated with micro-organisms, are a compelling experimental system that has been shown to sustain the vast majority of bacterial species found in native human microbiomes^25^ and recapitulate important aspects of human physiology and pathology^1, 26, 27^. From the standpoint of dynamical systems inference, gnotobiotic models allow for targeted strong perturbations and high-temporal resolution fecal sampling.

**Figure 3:**
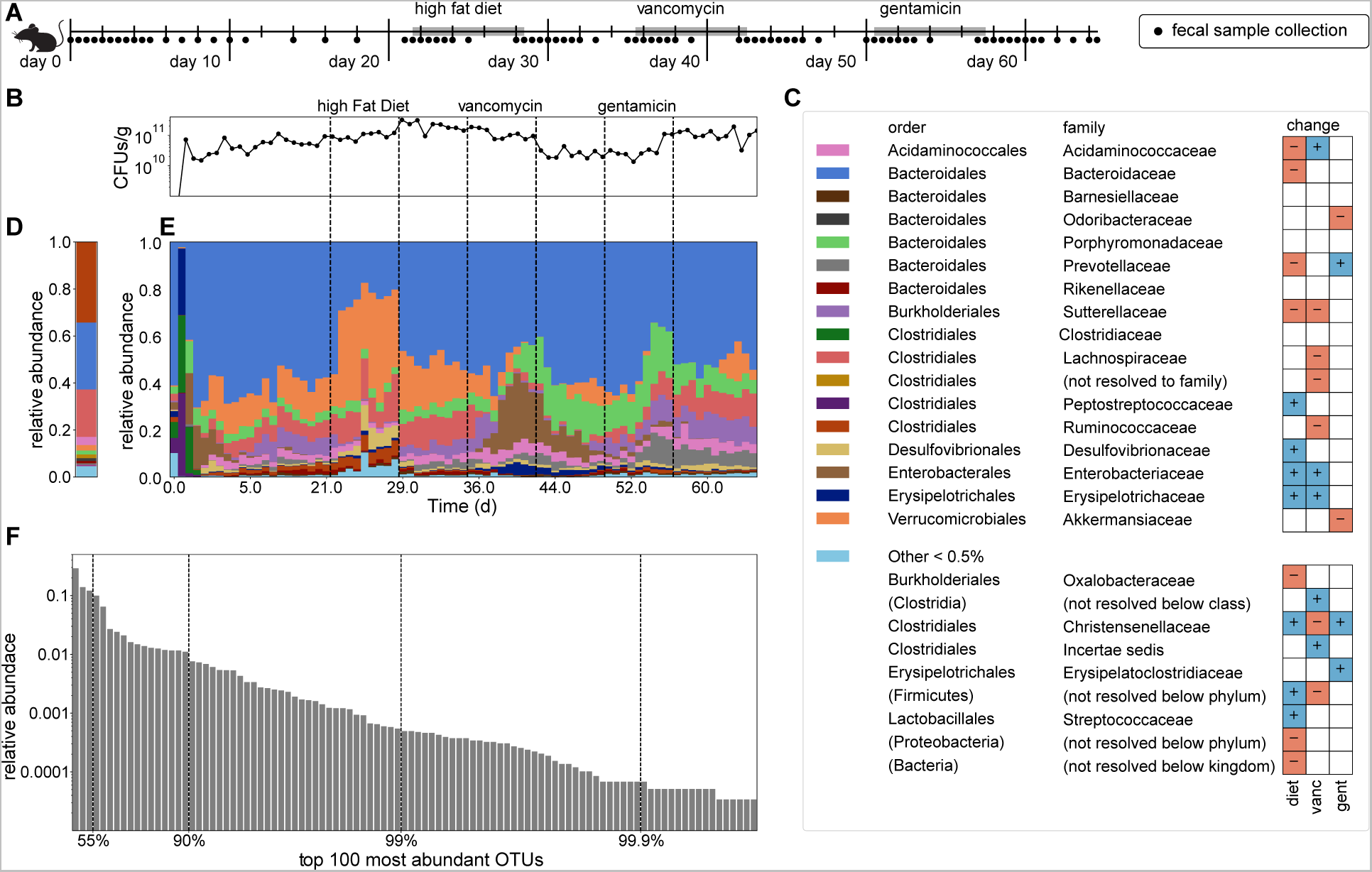
High-temporal resolution gnotobiotic mice colonization and perturbation studies using a human donor microbiome shows reproducible differential responses to perturbations. **(A)** Experimental design for the study (n = 4 mice) with an average of 77 serial fecal samples/mouse. **(B)** Average total bacterial concentrations in serial fecal samples. **(C)** Legend and taxonomy for panels D and E along with significant differential abundances for each taxonomic group across the three perturbations (see Supplemental Table 2 for *p*-values). **(D)** Relative abundance of microbes in human donor sample. **(E)** Relative abundances of microbes in serial fecal samples, averaged over the biological replicates. **(F)** Relative abundances of the top 100 most abundant ASVs across all mouse samples.

After an equilibration period of three weeks, mice were subjected to a sequence of three perturbations (high fat diet (HFD), vancomycin, and gentamicin) designed to provide rich data for dynamical systems inference, by differentially perturbating components of the microbiome (e.g., high fat/simple carbohydrate versus complex carbohydrate utilizers and bacteria susceptible or resistant to different antibiotics). Mice were separately housed and fecal samples were collected over a 65-day duration, with an average of 77 samples per mouse (Figure 3A). Samples were interrogated for relative abundance via 16S rRNA amplicon (Figure 3D, E) sequencing and total bacterial concentration via qPCR using a universal 16S rDNA primer (Figure 3B).

The resulting 24,027,980 million sequencing reads were bioinformatically processed using DADA2^28^, resulting in 1088 Amplicon Sequence Variants (ASVs). To yield a dataset with high-quality time-series information for dynamical systems inference tasks^18^, taxa that were not present at ≥0.01% relative abundance for seven consecutive time-points in at least two mice were filtered out, resulting in 141 ASVs for inference benchmarking; of note, this is more than 10X the scale of the original benchmarking dataset we provided to the community^1^. Figure 3F displays the relative abundances of the top 100 most prevalent taxa, illustrating the commonly seen phenomenon in gut microbiome datasets of orders-of-magnitude ranges in abundance among taxa (i.e., the top three most abundant taxa account for 55% of reads, and 90% of the reads account for only 17 taxa).

As expected from the experimental design, phylogenetically diverse groups of taxa underwent reproducible differential shifts from the applied perturbations (Figure 3C,E Supplemental Figure 1, Supplemental Tables 2,3). Standard univariate fold-change analyses^29^ demonstrated significant shifts across mice at the Phylum level, including decreases of Bacteroidetes and increases of Firmicutes on the high fat diet; decreases of Firmicutes and Actinobacteria, and increases of Proteobacteria on vancomycin; and increases in Firmicutes and Actinobacteria on gentamicin. At the Family level, significant shifts across mice were also seen in many taxa, including decreases in Prevotellaceae and Sutterellaceae, and increases in Peptostreptococcaceae, Desulfovibrionaceae, and Erysipelotrichaceae on the high fat diet; decreases in Sutterellaceae, Lachnospiraceae, and Ruminococcaceae and increases in Enterobacteriaceae and Erysipelotrichaceae on vancomycin; and decreases in Odoribacteraceae and Akkermansia and increases in Prevotellaceae on gentamicin. These results, showing consistent colonization with 141 ASVs in mice and reproducible differential responses to perturbations, coupled with the temporal resolution of the data, suggest its utility for inferring dynamical systems models, which we demonstrate in the subsequent sections.

### MDSINE2 outperformed state-of-the-art methods in forecasting microbial dynamics at scale

Using the experimental dataset we generated, we evaluated MDSINE2 and the gLV ridge and elastic net comparator methods for their ability to forecast held-out microbial concentrations. We additionally evaluated MDSINE2 without interaction modules (MDSINE2^–M^), to assess the impact of this feature on model performance and also to more directly compare our method to the state-of-the-art methods, which do not infer modules. For forecasting comparisons, we employed one-subject-hold-out training and testing methodology, e.g., holding out all data from one mouse, training on the remaining data, and forecasting all taxa trajectories for the entire 65-day time-series for the held-out mouse given only an initial data point. We evaluated performance using root-mean-squared error (RMSE) over the time-series, e.g., a measure of the difference between the predicted and ground-truth measurement.

Our method significantly outperformed both comparator methods on this benchmark for training with total bacterial concentration (Figure 4A, Supplemental Table 4). MDSINE2^–M^, which does not learn modules and instead encodes parameters for each pairwise interaction among taxa (19,740 parameters from 141 ASVs), showed slight but statistically significant better forecasting accuracy over MDSINE2 (272 parameters from a median of 17 interaction modules). This result is consistent with our prior finding that model constraints can impact forecasting performance to some extent^1^. However, the actual gap in performance between MDSINE2^–M^ and MDSINE was quite minor, suggesting that the much more compact dynamical system representation learned by MDSINE2 still captures the system behavior quite accurately. Given the large range in taxa abundances, we were interested in assessing how predictive performance varied with abundance, and in particular the impact on lower abundance taxa, which have an inherently larger relative variance^30^ due to being measured with fewer sequencing reads. MDSINE2^–M^ significantly outperformed the two comparator methods at all abundances (Figure 4B,C and Supplemental Table 5), including down to 10^5^ CFU/g, which is near the limit of detection for the dataset. MDSINE2 similarly significantly outperformed the comparator methods in all but the most abundant bin (concentrations greater than 2.5x10^9^ CFU/g). Interestingly, while MDSINE2^–M^ showed minor but statistically significant outperformance of MDSINE2 in the top 7 deciles (concentrations greater than 2x10^7^ CFU/g) there was no significant performance difference in the second and third lowest deciles ranging from 4.3x10^6^ CFU/g to 2x10^7^ CFU/g, and MDSINE2 significantly outperformed MDSINE2^–M^ for the lowest decile 1.1x10^5^ CFU/g to 4.3x10^6^ CFU/g. These results suggest that interaction modules capture dynamics well across the range of taxa abundances and may even improve accuracy for the least abundant taxa. We note that none of the methods, including our own, can predict exactly zero abundances. Analysis of cases in which the held-out sample ASVs have zero reads show poor predictive performance for all methods, which all generally predict values near the limit of detection of 10^5^-10^6^ CFU/g, as expected (Supplemental Figure 2A). Interestingly the method that had the best performance for the zero read cases, gLV-L2, had the worst performance in all other cases.

**Figure 4:**
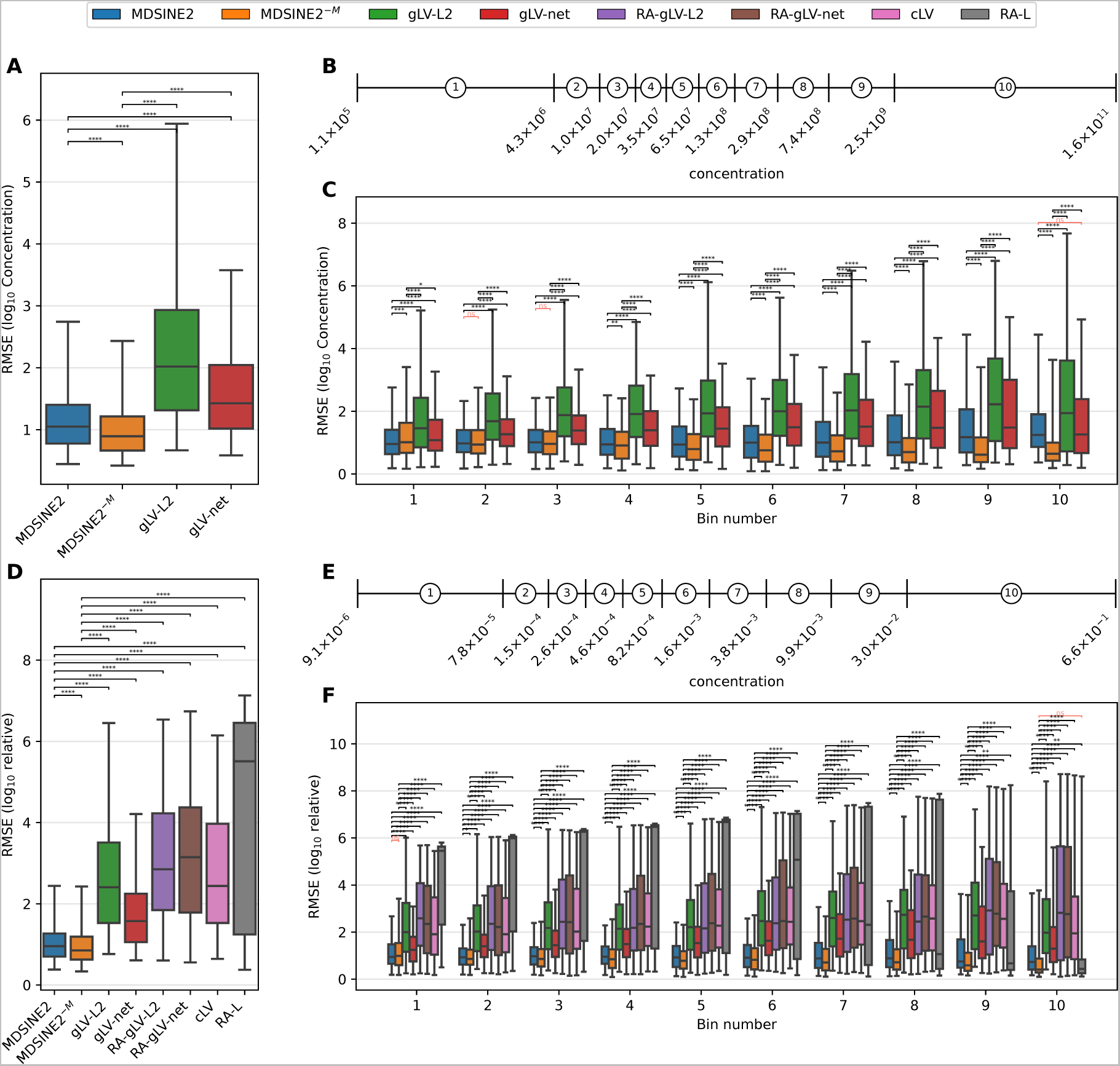
MDSINE2 outperforms state-of-the-art methods in forecasting microbial concentrations and relative abundances. Results are for cross-fold validation of forecasts. Models were trained on three of four mice, then given the initial condition of the hold out mouse time-series, and then the inferred model was simulated forwards and the output was compared to the entire held out time-series. All tests are two-sided Wilcoxon signed-rank tests. Non-significant tests are denoted with red bracket. Errors are calculated from forecasted abundances with respect to hold out data across all folds and for all taxa with non-zero reads within each sample. **(A-C)** Results for models trained on microbial concentrations. **(A)** Forecasting error for all non-zero taxa samples. **(B)** Concentration ranges for deciles. **(C)** Forecasting error grouped by decile of holdout sample taxa concentrations. **(D-F)** Results comparing models trained on concentrations and relative abundances. **(D)** Forecasting error for all non-zero taxa samples. **(E)** Relative abundance ranges for deciles. **(F)** Forecasting error grouped by decile of holdout sample taxa relative abundance. * p<0.05, ** p<0.01, *** p<0.001, **** p<0.0001, ns: not significant (p>0.05).

To assess the role that bacterial concentration information gives for model forecasting, we compared the performance of the four models just described against other state-of-the-art methods that learn from relative abundance information alone. These methods include: compositional Lotka Volterra (cLV)^15^, linear dynamics trained on relative abundances (RA-L), gLV ridge trained on relative abundances (RA-gLV-L2), and gLV elastic net trained on relative abundances (RA-gLV-net). Overall, MDSINE2 and MDSINE2^–M^ outperformed the comparator methods, and MDSINE2^–M^ ultimately had the best performance (Figure 4D,E,F and Supplemental Tables 6, 7). However, we found that methods trained on data that included the qPCR measurements significantly outperformed methods trained on only relative abundance information, with the only exception being gLV-L2 compared to cLV (Figure 4F, Supplemental Table 6). These results highlight a well-studied issue with relative abundance data, that increases in one taxa’s abundance are indistinguishable from other taxa’s abundances simultaneously decreasing^31^, which has implications for identifiability (the ability to learn model parameters from data) from a dynamical systems perspective^18^.

### MDSINE2 learned a compact set of interaction modules that revealed keystone sets of species and stability characteristics of a complex gut microbial ecosystem

On our benchmarking dataset, MDSINE2 discovered 17 interaction modules (Figure 5, Supplemental Table 8), ranging in size from 1 to 35 taxa, and connected through 56 interactions predicted with *decisive evidence* (Bayes Factor [BF] ≥ 100, Figure 5C)^32^. This represents a 97% reduction in interaction parameters over MDSINE2^-M^ (2179 edges predicted with *decisive evidence* [Supplemental Figure 5]), with nearly comparable forecasting performance as described in the previous section. As one measure of the biological relevance of modules, we evaluated the relatedness of taxa within modules compared to random assortments of taxa. Using phylogenetic distance as a metric, we found that the average within module distance was significantly lower (*p*=0.0135) than chance (Supplemental Figure 3). From the taxonomic perspective, five modules were significantly enriched at the Family level with two at the Order, two at the Class, and four at the Phylum levels, including expected segregation of modules along Gram positive and negative Phyla, e.g., Bacteroidetes and Firmicutes (Supplemental Figure 4)^32^. These results suggest that the modules, which are wholly learned from data measuring the dynamic behavior of the microbe, also significantly reflect phylogenetic signal.

**Figure 5:**
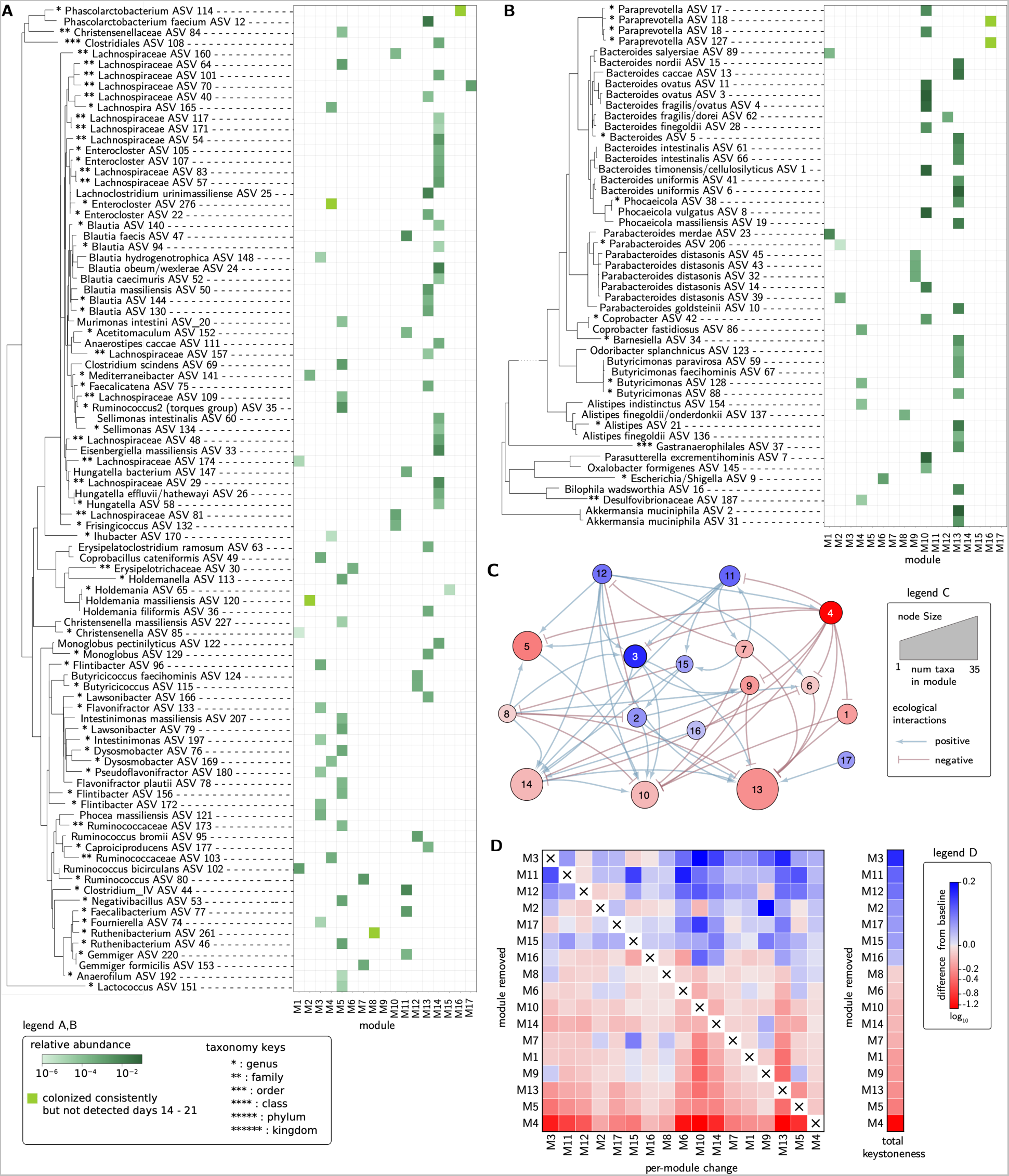
MDSINE2 infers modular representations of complex microbiome dynamics. Our method automatically learns modules of ASVs based on similarity of their dynamic interactions and responses to perturbations. Results are split into: **(A)** Gram-positive, and **(B)** Gram-negative ASVs, for display purposes. **(A2,B2)** Phylogenetic tree of ASVs. **(A3,B3)** Module memberships. Intensity of color in the grid indicates abundance post-colonization and prior to perturbations (average over Days 14 to Day 21). **(C)** Inferred module interaction network displaying only edges with BF > 100 (*decisive* evidence). Size of nodes are proportional to the number of ASVs in the module and color of nodes corresponds to module keystoneness. **(D)** Keystoneness analysis measures the relative importance of a module in maintaining the steady state abundance of the community. A positive keystoneness for a module means its removal from the community reduces the steady state abundance of its community members, and a negative keystoneness means that the module’s removal results in an increase in the abundance of the other community members. A module’s impact on the other community members was computed both at the module level and across all modules (total keystoneness).

To evaluate quantitatively the relative importance of interaction modules in the ecosystem, we performed a module-level keystoneness analysis^1^ (Figure 5D, and Methods §4.6). Keystone taxa were originally defined as those taxa that are fundamental to the integrity of the ecological community, without whom the community would collapse^34^, and have been suggested as drivers of microbial community structure and function^35, 36^. Here, we extend the concept to groups of taxa (modules), to aid in interpretability, and also generalize to a quantitative measure of *keystoneness* with both positive and negative values. Positive keystone modules (“promoters”) are those that when removed result in a reduction in the microbial abundances of the other members of the ecosystem; negative keystone modules (“suppressors”) are those that when removed result in increases of abundances of the other members of the ecosystem. The magnitude of the keystoneness measure thus represents the degree of community wide disruption (in terms of microbial abundance change, with the removal of the module).

For our cohort, the top positive and negative keystoneness modules were M3 and M4 respectively. Investigating their role through the ecological network, we see that all the outgoing edges of M3 are promoting, while all the outgoing edges of M4 are repressive, suggesting the different ecological roles that these modules play in the network. M3 is enriched for the family Ruminococaceae. Promoting M3 are two other positive keystoneness modules M11 and M12, each containing taxa capable of degrading resistant starches (*Ruminococcus bromii* ASV95, *Gemmiger* ASV220) and others with butyrate production capabilities (*Faecalibacterium* ASV77, *Butyricicoccus faecihominis* ASV124, *Butyricicoccus* ASV115). Downstream modules being promoted by M3 of note are M10 (enriched for Bacteroidaceae), M13 (enriched for Bacteroidetes and the largest module in the network), and M14 (enriched for Lachnospiraceae). One explanation for this structure, consistent with known biology, is that the positive keystoneness modules are connected in a cross-feeding chain beginning with specialized starch degrading taxa that ultimately support the more abundant generalist taxa (e.g., Bacteroidaceae). In contrast to the this specialist-to-generalist structure, the module with the highest negative keystoneness, M4, contains a diverse group of taxa and also suppresses multiple modules in the network, including M3 and M11, the top two positive keystoneness modules, as well as the primarily gram-negative modules M10 and M13. An annotated module network can be found in Supplemental Figure 6.

**Figure 6:**
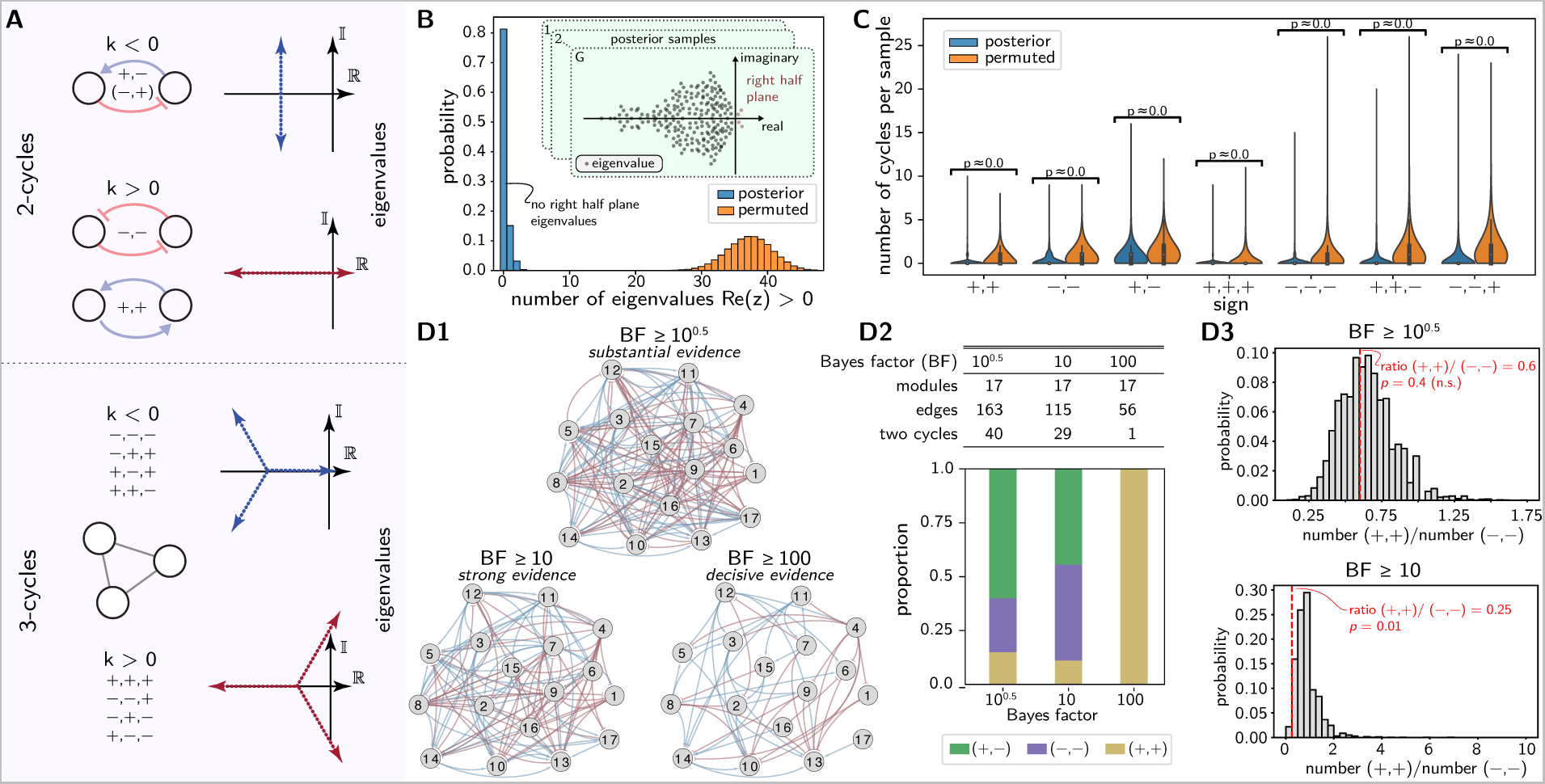
Analytical stability analysis of microbiomes demonstrates significantly more stable dynamics than expected by chance and reveals cycle feedback motifs. **(A)** two and three-cycle motifs and their corresponding eigenvalue asymptotes for increased interaction strength. Only the two-cycle with negative feedback can remain stable for all feedback gains (top motif). **(B)** As a measure of stability, we computed the number of right half plane eigen-values for each posterior sample of our model trained on the microbiome data, and for the null model (“permuted”). Our model had an 80% probability of having no unstable eigenvalues. **(C)** Distribution of cycle motifs over posterior samples. Analyses showing **(D1)** networks derived with three different levels of confidence for including edges, **(D2)** descriptive statistics for the networks, and **(D3)** statical test for significance of mutualism to competition ratio (MCR [(+,+)/(−,−)]), which was significant for the network constructed from edges with *strong* evidence *p*=0.01.

To assess the overall robustness of our inferred ecosystem to external perturbations we performed a formal stability analysis. Informally, stable systems are those systems that when undergoing a perturbation do not deviate far from their equilibrium state, and when that perturbation is removed, return to their original equilibrium state (asymptotic stability). Healthy gut microbiomes have many empirical features of a stable dynamical system, such as maintaining an equilibrium taxa abundance profile over long periods of time (years), and a rapid return to equilibrium (within days) after perturbations are removed (e.g. antibiotics, diet change, etc.)^33, 37–39^. MDSINE2 learns an ecosystem-scale model of dynamics, and we can thus perform formal analyses of the underlying model parameters and equations to gauge stability. To compute stability, we used the fact that for gLV dynamics, a necessary condition for asymptotic stability is that the eigenvalues of the pairwise interactions matrix have all negative real parts (Supplemental Text §3)^40^. We note that prior studies have performed similar analyses, but for much smaller models trained on only tens of taxa^1,7,13^.

Overall, we found that the dynamics of the model inferred were 80% likely to be stable (Figure 6B). This is consistent with prior work on smaller microbial networks suggesting that dynamics inferred from a healthy microbiome are indeed likely to be stable^7^. To assess how likely this scenario is given a network of the same size, we created a randomized network (random rewiring) that maintained a core topological feature of the network (the in-degree for each node was maintained) but that had the edges permuted, meaning that the number of interactions incoming to each node is maintained (Methods §4.8). With this random rewiring the system had a 0% probability of being stable (Figure 6B, Methods §4.7, Supplemental Text §3). These results indicate that the model MDSINE2 inferred from a healthy, complex gut microbiome is far more stable than would be expected by chance and highlights our method’s ability to perform dynamical systems analyses at ecosystems-scale.

We next sought to identify features of the ecological network inferred by MDSINE2 that could explain the remarkable stability of healthy gut microbiomes. Stability and control theory have established that the feedback cycle is the core topological feature driving stability^41^. Pairwise interactions, the simplest form of feedback cycles, have particular interpretations in ecology, and their contributions to stability are well-characterized for linear and gLV dynamical systems^6^: mutualism (+,+) and competition (−,−) are destabilizing, and parasitism (+,−) is stabilizing. For length three cycles and higher, more complex ecological interactions arise, and any sign combination is potentially destabilizing^42^ (Figure 6A). For all cycle lengths analyzed, we found that MDSINE2’s inferred model of dynamics had a significantly lower number of cycles than expected by chance (Figure 6C). Of note, the most prevalent network motif in the inferred model was the always stable negative feedback 2-cycle (+,−); however, this motif was also present significantly less frequently than expected by chance.

We next sought to understand the influence of uncertainty in the inferred network structure itself on stability estimates, given that ecological networks of the microbiome are inferred from relatively limited and noisy data. To gain insight into this phenomenon, we evaluated networks at different levels of evidence for edges, *substantial* [BF ≥ 10^0.5^], *strong* [BF ≥ 10], and *decisive* [BF ≥ 100] evidence (Figure 6D1, Supplemental Text §G). As the evidence threshold for an edge being included in the model was decreased, the number of edges in the network increased from 56 to 163 (Figure 6D2). As the number of edges increased, there was also an increase in the number of (two) cycles, as expected. Interestingly, for networks with more edges, the number of parasitism (+,−) cycles increased disproportionally among two cycles present, consistent with the property that stability becomes less likely the denser a network becomes (the more edges there are for a fixed number of nodes)^5,43, 44^, unless the cycles in the network are only parasitism (Figure 6A). Previous work has also examined the role of mutualism and competition cycles, hypothesizing that for healthy ecosystems, the mutualism to competition ratio (MCR [(+,+)/(−,−)]) would be less than one^45^, and demonstrating this phenomenon on networks inferred on small microbial ecosystems^7^. Our results provide support for this hypothesis on the complex, healthy gut microbiome, showing the that the mutualism to competition ratio was significantly lower than chance on networks with strong evidence for the existence of edges (Figure 6D3).

## Discussion

MDSINE2 is a fully Bayesian method that propagates uncertainty throughout the model^46^, including explicit models of measurement noise in both relative abundance and total bacterial concentration, which we calibrate using physical and technical replicates. This framework allows us to achieve unprecedented performance improvements over prior methods, including our own, which did not fully propagate measurement uncertainty. Another important component of our model is the ability to analyze microbial dynamical systems in terms of modules. Modularity has been extensively exploited for understanding other complex biological systems, such as mammalian genetic regulatory networks^47–49^, but relatively underexploited in the microbiome field^19, 20^.

Dynamical systems inference requires data with sufficient richness of information. Measurements taken when trajectories are significantly deviating from steady state, e.g., transients, are critical for efficient inference^18^. To generate sufficiently rich data, we performed multiple strong perturbations in a cohort of gnotobiotic mice serially over time. This experimental design allowed us to perform temporally dense sampling with biological replicates (effectively multiple initial conditions), coupled with strong perturbations. Thus, our time-series data, specifically generated for efficient dynamical systems inference, is expected to be a valuable resource to any researcher studying ecological dynamics.

Dynamical systems analyses can offer insights into how ecosystems behave under physiologically relevant perturbations, and an understanding of these dynamics can then in turn be used to develop strategies to control the ecosystem^50–52^. Our analyses on longitudinal data from a cohort of mice colonized with a microbiome from a healthy human donor identified specific (e.g., putative cross-feeding chains beginning with resistant starch degrading organisms) and general (e.g., topological motifs) properties in the ecological network that promote stability. These findings, and the overall analysis methodology we have introduced, provide a foundation for designing interventions to increase the stability of dysbiotic microbiomes.

There are several directions for future work on the computational model. First, MDSINE2’s model of measurement noise, a well-established model based on the negative binomial distribution, could be extended to include zero inflation or nonparametric features (which could be particularly advantageous for metagenomics or other data types that may not fit the negative binomial model well). A second direction for extension is the gLV model, which only allows for pairwise, “mass action” style interactions. Although it would be conceptually straightforward to include higher-order interactions, it remains unclear how prevalent such interactions are in mammalian microbiomes^58^; another approach would be to include interactions with saturation behavior, or even learn nonparametric models for interactions to avoid needing to prespecify their forms. A third, and particularly exciting area for extension, would be incorporating additional data modalities to more fully capture the host-microbial ecosystem, such as metagenomic, metatranscriptomic, and metabolomic data sources.

## Conclusion

We have introduced MDSINE2, a computational method for accurately inferring interpretable dynamical systems models of the microbiome at scale and demonstrated on synthetic data and a new resource of densely sampled microbiome time-series data, from “humanized” gnotobiotic mice, that our approach outperforms other methods when forecasting microbiome dynamics or predicting ground truth dynamics. Our approach provides new tools for characterizing the dynamical systems behaviors of complex host-microbial ecosystems and holds promise for guiding rational design of interventions to stably alter human microbiomes for prophylactic or therapeutic purposes.

## Methods

## 1. MDSINE2 Model

### 1.1 Overview

Our statistical model of microbial dynamics is a fully Bayesian model based on continuous-time stochastic generalized Lotka-Volterra (gLV) dynamics:

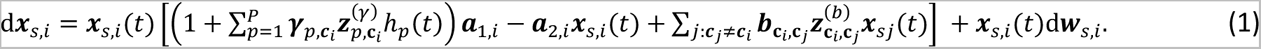

This formulation of stochastic behavior models multiplicative random effects on microbial abundances, which could arise from a variety of phenomena, such as temporal host, environmental or dietary fluctuations that result in short time-scale increases or decreases in abundance of each taxa.

The abundance of taxa *i* in time-series *S* (e.g., biological replicate) is denoted as ***x**_si_*. MDSINE2 probabilistically assigns each taxa to an interaction module, where ***c***_*i*_ denotes the module assignment for taxa *i*. The growth rate and self-interaction random variable for taxa *i* are denoted ***a***_1,*i*,_ and ***a***_2,*i*_, respectively. The *P* external perturbations are accounted for by the random variables ***γ***_p,***c***_*i*__ that denote the effect of perturbation *p* on taxa *i*’s growth rate; 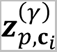 is a corresponding random indicator variable that probabilistically selects whether the perturbation affects the interaction module. The function ℎ*_p_*_’_ has a value of 1 during the time-period when the *p*-th perturbation is active and a value of 0 otherwise. The strength of the microbial interaction from taxa *j* to taxa *i* is denoted ***b***_*ci*,*cj*_, with 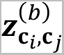 the corresponding random indicator variable for that microbial interaction. The stochastic variation of the microbial abundances over time is captured by the variable ***w**_s,i_*, specifying geometric Brownian motion for the stochastic component (e.g., a multiplicative stochastic process on the microbial abundance).

To support efficient inference, we use a first-order discretization (see Supplemental Text) to yield the discrete-time latent trajectories:

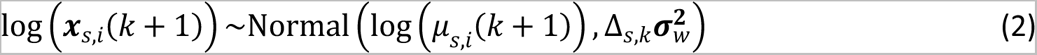

Where

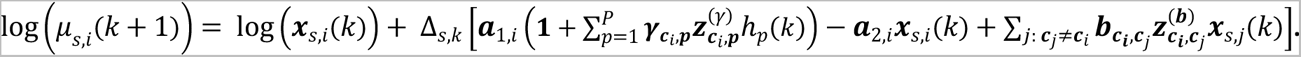

Here, Δ_*S*,*k*_= *t*_*S*,*k*+1_ − *t*_*S*,*k*_, the difference between adjacent time-points for the time-series *S*. Below we give additional details on the model, including prior probability distributions on variables; for complete mathematical and algorithmic details, see Supplemental Text.

### 1.2 Interaction Modules

We employ a Dirichlet Process (DP) prior^59^ to model interaction modules. The expected number of modules under this prior probability distribution is 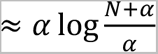, where *N* is the number of taxa and *α* is the concentration parameter^60^. This property is desirable for scaling to large ecosystems, as the expected number of microbial interactions in our model scales as *O*(log(*N*)^2^) (as opposed to *O*(*N*^2^) in the standard gLV model). We place a diffuse Gamma prior on the concentration parameter as described in^61^. Our formulation allows us to marginalize out the interaction and perturbation parameters during inference, which greatly increases efficiency^59^. See Supplemental Text for complete details.

### 1.3 Interaction Parameters and Perturbation Effects

To facilitate modularity and interpretability of inferred interaction networks, we assume no intra-module interactions and model only inter-module interactions, ***b***_***c**i*,***c**j*_ . We assume perturbations (e.g., antibiotics or dietary changes) have module-specific effects, ***γ***_***c**i*_. Further, we model the presence/absence of module-module interactions and module-perturbation effects by using the binary indicator variables ***z***^(***b***)^ and ***z***^(***γ***)^, respectively. These binary indicators allow the model to infer the structural edges that specify the underlying network topology between modules. Additionally, this formulation allows for direct calculation of the statistical evidence for presence of each interaction or perturbation effect using Bayes factors. See Supplemental Text for full details.

### 1.4 Measurement Model

The observed data are sequencing counts ***y***_*s*,*i*_(*k*) of taxa and qPCR measurements ***Q***_*s*,r_(*k*) of bacterial concentrations, where *j* indexes the qPCR measurement replicates. Sequencing counts are modeled using a negative binomial distribution^30^

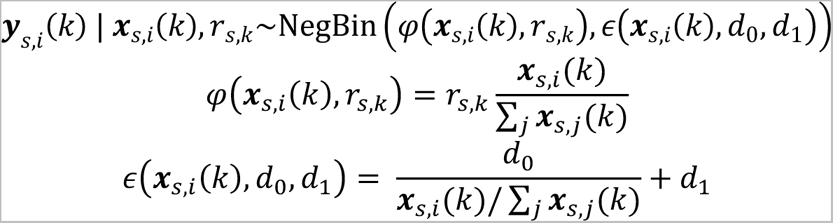

Here *r*_*S*,*k*_ is the total number of reads for the sample in time-series *s* at time *t*_*S*,*k*_, and *d*_0_ and *d*_1_ parameterize the function *∈*(⋅), which specifies the Negative Binomial dispersion parameter. We fit the parameters *d*_0_and *d*_1_using data from replicates (see below).

We model the qPCR measurements with a log-normal distribution:

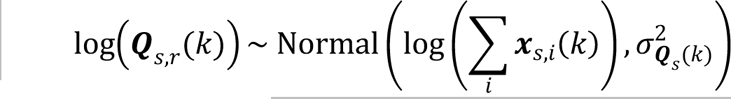

Here, 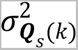 is the empirical variance of the set of qPCR measurement replicates for time-series s at time *t*_*S*,*k*_. See Supplemental Text for complete mathematical details of the measurement model and inference procedure.

### 1.5 Software

MDSINE2 was implemented in Python 3.7 using the Numpy^62^, Scipy^63^, Numba^64^, Matplotlib^65^, and Seaborn^66^ packages. The software is publicly available under the Gnu General Public License v3.0 (https://github.com/gerberlab/MDSINE2). The input to MDSINE2 consists of five tab-delimited files: (1) list of the sequence and taxonomic label for each taxa, (2) table of counts for each taxa in each sample, (3) table specifying the time points at which each sample was collected for each subject, (4) table of qPCR values for each sample, and (5) table of perturbation names, start times, end times, and associated subjects that received the perturbation. The software outputs inference results in two files: (a) a Python pickle file that contains the MDSINE2 inference objects, and (b) a HDF5 file containing all the MCMC posterior samples. Once inference is complete, the software includes functionality to visualize and interpret the posterior samples, including visualizing trajectories, module networks (with a Cytoscape^67^ export option) and keystoneness, as well as generating text files with summaries of posterior distributions. See online software documentation for complete details. We also give demos of the functionalities in the tutorials. The tutorials (Google Colab) can be accessed from https://github.com/gerberlab/MDSINE2_Paper

## 2. Gnotobiotic Experiments and Microbiome Data Generation

### 2.1 Mouse Experiments

A cohort of four male C57Bl/6 germfree mice was used in the experiments (BWH IACUC: 2016N000141). Mice were singly housed in Optmice cages within the Massachusetts Host-Microbiome Center (MHMC) at Brigham and Women’s Hospital.^25^ The mice were given a Fecal Microbiota Transplant (FMT) from a healthy human stool donor from an ongoing study at Brigham and Women’s Hospital (IRB# 2017P002420). Per the study protocol, samples were flash frozen without cryoprotectants and stored at -80°C. Material for FMTs was prepared by thawing the stool samples and homogenizing in 5 mL of pre-reduced 1x Phosphate Buffered Saline (PBS) with 0.05% cysteine inside an anaerobic chamber. Germfree mice were then orally gavaged with 200µl/mouse of FMT material. Post-gavage, mice were equilibrated for 3 weeks before beginning a series of three perturbations: high fat diet (HFD), vancomycin, and gentamicin (in that order). Each perturbation lasted for one week, followed by a one-week normalization period off perturbations. Aside from the HFD perturbation, mice were maintained on standard MHMC gnotobiotic mouse chow (Autoclavable Mouse Breeder Diet 5021; LabDiet). For the HFD perturbation, Research Diets D12492 (60 kcal% of fat) was used. For the vancomycin perturbation, drinking water was replaced with water containing vancomycin at a concentration of 100 ug/mL and 3% sucralose (filter sterilized). For the gentamicin perturbation, drinking water was replaced with water containing gentamicin at a concentration of 4 ug/mL and 3% sucralose (filter sterilized). In all situations, mice were allowed to eat and drink ad libitum. Mouse fecal pellets were collected in triplicate based on the sample collection timeline detailed in Figure 2. We also obtained additional samples to generate data for fitting the d0 and d1 parameters in our amplicon sequencing measurement noise model. For this purpose, a total of nine fecal pellets, three pellets on each of the three consecutive days (8, 9, 10) were collected from mouse 2. Each fecal pellet was divided into two parts. This resulted in 18 samples that were then processed through the entire sequencing pipeline, from DNA extraction through sequencing. To collect fecal pellets, each mouse was removed from the Optimice cage and placed inside an autoclaved Nalgene cup. After pellets were produced, mice were placed back in their cages and samples were collected from the cup with autoclaved forceps. Samples were placed in cryovial tubes and snap frozen in liquid nitrogen immediately, then stored at -80°C. At the end of experiments, mice were euthanized by overdose on inhaled vapors of isoflurane administered in an anesthesia chamber followed by cervical dislocation. These procedures are in accordance with the recommendations of the Panel on Euthanasia of the American Veterinary Medical Association.

### 2.2 DNA Extraction, 16S rRNA Amplicon Sequencing and qPCR

For DNA extraction, all samples were processed using the standard protocol^68^ at the Massachusetts Host-Microbiome Center (MHMC), which uses the Zymo Research ZymoBIOMICS DNA 96-well kit according to manufacturer instructions with the addition of bead beating for 20 minutes. Amplicon sequencing and qPCR were also performed using the standard MHMC protocol. Briefly, for amplicon sequencing, the v4 region of 16S rRNA gene was PCR amplified using 515F and 806R primers^69^: 5’-[Illumina adaptor]-[unique bar code]-[sequencing primer pad]-[linker]-[primer]

- (fwd primer): AATGATACGGCGACCACCGAGATCTACAC-NNNNNNNN-TATGGTAATT-GT-GTGCCAGCMGCCGCGGTAA
- (rev primer): CAAGCAGAAGACGGCATACGAGAT-NNNNNNNN-AGTCAGTCAG-CC-GGACTACHVGGGTWTCTAAT

Following PCR of the v4 region, 250 bp paired end reads were generated on an Illumina MiSeq with the following custom primers with index primer: ATTAGAWACCCBDGTAGTCC-GG-CTGACTGACT.

- 5’-[sequencing primer pad]-[linker]-[primer] Read 1: TATGGTAATT-GT-GTGCCAGCMGCCGCGGTAA
- 5’-[primer]-[linker]-[sequencing primer pad] Read 2: AGTCAGTCAG-CC-GGACTACHVGGGTWTCTAAT

qPCR for estimating total bacterial concentration was performed using universal 16S primers and a standard curve prepared from dilutions of *Bacteroides fragilis* (ATCC 51477). Samples were loaded into 384 well plates via the Eppendorf EP Motion liquid handler and then run on a QuantStudio 12K Flex Real-Time PCR System (ThermoFisher) using TaqMan Universal Master Mix II no UNG kit (ThermoFisher 4440040), TaqMan Gene Expression Assay (ThermoFisher 4331182), probe set Dye: FAM, Quencher: NFQ-MGB and Reference Dye: Rox for quantification (ThermoFisher assay ID Pa04230899_s1), all according to manufacturer’s instructions.

## 3. Bioinformatics

### 3.1 Generating ASV tables From Amplicon Reads

We generated an ASV read count table and assigned taxonomy using *DADA2 v1.16* according to the standard pipeline using pseudo-pooling^28^. Forward reads were trimmed to a length of 240 and reverse reads were trimmed to a length of 160. Our function calls for these core steps in the DADA2 pipeline were:

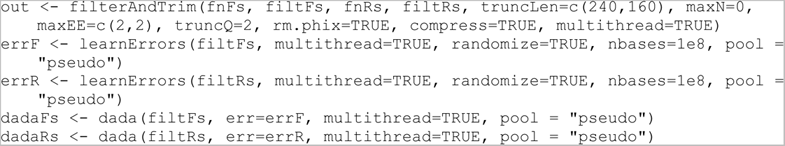

To assign taxonomic labels to ASVs, we used DADA2-formatted reference databases RDP trainset 16 and Silva version 138. When using `assignTaxonomy` in DADA2, we specified the maximum number of multiple species assignments to be 2. For species assignments, if one database returned a species assignment and the other did not, we labelled the ASV with the species from the database that returned the assignment. If both databases returned species assignments, but they were discordant, we set the assignment to the union of the returned assignments. If the total number of possible species assigned was greater than 4, then we did not set a species assignment. This process resulted in 1088 ASVs in total.

### 3.2 Phylogenetic Placement of Sequences

We performed phylogenetic placement of consensus ASVs onto a reference tree constructed from 16S rRNA sequences of type strains tagged as “good” quality, length between 1200bp and 1600bp in RDP 11.5 ^70^. We performed multiple alignment of the sequences using the RDP’s web-hosted alignment tool with default parameters ^71^. To facilitate a good multiple alignment, we filtered out sequences with insertions seen in ≤ 3 other sequences. A reference tree was constructed using FastTree^72^ version 2.1.7 SSE3 with the general-time-reversible maximum likelihood option. For phylogenetic placement, the aligned reference sequences were first trimmed to position 1045 to 1374 (corresponding to the region flanked by the 16S v4 primers) and a hidden Markov Model was learned using *hmmbuild* in HMMER v3.1^73^. ASV sequences were then aligned using *hmmalign* with the -*mapali* option. Finally, the aligned sequences were phylogenetically placed using *pplace*r *v1.1.alpha19* with default settings^74^.

### 3.3 Fold Change Analysis

Fold change analysis was performed using DESeq2^29^ v1.3.2.0. All default options were used with features only kept if there were at least 100 reads (summing across all the samples used in the analysis) using the following commands

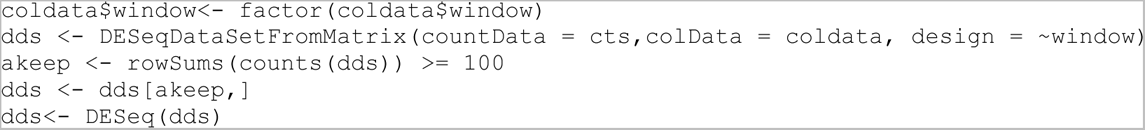

The scripts to perform this analysis are contained in https://github.com/gerberlab/MDSINE2_Paper/tree/master/analysis/deseq. The fold changes were calculated using the default Wald test in the software. The fold changes during the perturbations were calculated with respect to the “steady states” achieved just before the perturbation was applied. The HFD fold change was calculated by comparing days (23, 23.5, 24, 25) to days (16, 18, 21, 21.5), the vancomycin fold change was calculated by comparing days (37, 37.5, 38, 39) to (32, 33, 35, 35.5) and the gentamicin perturbation fold change was calculated by comparing days (52,52.5,53,54) *to* (46,47,50,50.5).

## 4. MDSINE2 analyses

### 4.1 Model Inference

Day 0 and 0.5 samples were excluded from inferences, due to their very low overall bacterial concentrations. First, using replicate data, we performed a fit for the negative-binomial model resulting in *d*_0_ = 4.2 × 10^-8^, *d*_$_ = 6.05 × 10^-2^. Using these hyperparameters, inference was performed using ten seeds configured to learn modules. For each seed, models were inferred using 10,000 MCMC iterations after 5,000 burn-in steps. Then, inference was performed on a single seed in fixed-cluster mode, where the consensus modules were derived using the concatenated outputs of the ten seeds. To assess convergence of each Markov chain, we used the 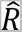 statistic^46^ and confirmed values of 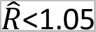 for the concentration parameter, the growth rates and the process variance, to indicate sufficient mixing. Inference took ∼26.5 hours (6.3 sec/iteration) on an Intel Xeon E5-2697V2 (2.70 GHz base frequency, 3.50 GHz max frequency) with an allocation of one core and 8 Gb of RAM separately for each task. Full details on inference are given in the Supplemental Text.

### 4.2 Benchmarking

We used implementations of the comparator methods provided in https://github.com/tyjo/clv. Following Joseph et al. (2020)^15^, we trained the models using elastic-net regression, and in addition, we trained gLV using ridge regression to provide comparisons to earlier work ^1,13^. Predictive performance of methods was assessed using a hold-one-subject-out cross validation procedure. Per fold, each method was provided data from all but one mouse in the cohort to infer model parameters. The inferred parameters were then used to forward simulate the trajectory of the held-out mouse, using the abundance at Day 1 as the initial condition. For comparator methods, the Runge-Kutta “rk45” procedure was used, as implemented in Joseph et al. (2020)^15^. For MDSINE2, each posterior sample was used to deterministically forward simulate (Equation 2 with no process variance), and the median of the distribution of simulations was used as the final forecast. The methods use different approaches to handle zeros in data. To make results as comparable as possible, we used the following settings. For gLV-elastic net and gLV-ridge, we set the minimum value for taxa to 10^C^ CFU/g, which is consistent with the limit of detection in our experiments. For gLV-ra and LRA, which are relative abundance methods, we set the minimum to 10^AD^ and for CLV, we set the additive offset *∈* = 10^AD^, consistent with the limit of detection for relative abundances in our experiments. These minimums were enforced both in data preprocessing and on the simulated trajectories, so that results remained comparable. For comparisons against methods that operate only on relative abundances (cLV, gLV-ra and LRA), we converted predictions of MDSINE2 or the gLV-based models to relative abundances. The following Root Mean Square Error metric was used:

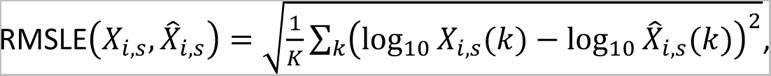

where *X_i,s_* denotes the measurements for taxa *i* in the held-out mouse *s* and 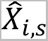 are the respective forecast estimates. In order to compare the errors between MDSINE2 and other methods, we performed one-tailed Wilcoxon signed-rank testing. The paired data points used for the test are the RMSLEs associated with MDSINE2 and the RMSLEs associated with the comparator method for all the ASVs in the hold-out subjects.

### 4.3 Synthetic Data

Benchmarking of models with synthetic data in Figure 2 followed the data generation procedure used to benchmark MDSINE in the original Bucci et al. (2016)^1^ manuscript. Briefly, gLV dynamics were simulated for a community of ten taxa with ten biological replicates for a total of 30 days each. Nine taxa were simulated in the system at day 0 and the challenge taxon was introduced on day ten. Measurements were assumed to be daily, with qPCR measurements simulated from a Lognormal Distribution and reads were simulated from a Dirichlet-Multinomial Distribution.

### 4.4 Consensus Module Construction and Fixed Module Inference

Consensus modules were constructed by performing agglomerative clustering on the co-clustering probability matrix where the number of clusters was the median number of modules over the posterior. See^19, 20^ for additional details.

### 4.5 Taxonomic Enrichment Analysis

Using the consensus modules, we performed enrichment analysis at four taxonomic levels: family, order, class, and phylum. The enrichment analysis was carried out using the hypergeometric test followed by the Benjamini-Hochberg procedure for multiple hypothesis tests. The hypergeometric probability is defined as 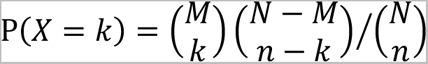. Here, *N* is the total number of ASVs used in the model, *M* is the total number of ASVs associated with a given taxonomic level, *n* is the size of the interaction module, and *k* is the number of ASVs in the interaction module that is associated with the given taxonomic level.

### 4.6 Keystoneness

The keystoneness measure was computed by removing all the taxa for each module *m*, forward simulating trajectories (as described in Benchmarking) for the remaining taxa over 100 days and comparing the final state of these trajectories to the final state with all taxa present in the ecosystem. As in our perturbation experiments for stability analysis, final states were computed as the mean of values over the last 12 hours in the final simulated day. To be precise, final state estimates *x*^(*g*)^(full system) and 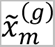 (system with module removed) for each MCMC step *g*, which are used to compute the keystoneness measure, are given by:

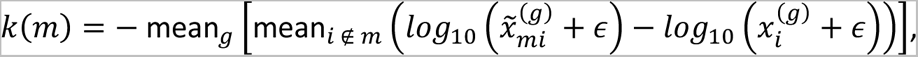

where the subscript *i* denotes the taxon index. Just as in the simulation-based stability analysis, ɛ = 10^5^. Per this formulation, a positive keystoneness value indicates an overall decrease in the system on average (meaning *m* has a positive effect on other ASVs when present), while a negative keystoneness value indicates an increase (meaning *m* has a suppressive effect when present).

### 4.7 Stability

As a measure of stability, we computed the number of right half plane eigenvalues for the interaction matrix in each posterior sample. For a null model we generated in-degree preserving permutations of the interaction matrix (permutations are performed at the module level for each sample, see Methods §4.8). The number of right half plane eigenvalues for the interaction matrix was then determined by counting the number of eigenvalues whose real part was greater than zero. For a theoretical discussion on the use of the eigenvalues of the interaction matrix for determining stability, see Supplemental Text §3.1.

### 4.8 Network Null Model

To provide test of statistical significance for network topological features, we generated indegree-preserving permutations of the interaction matrix, performed by permuting the off-diagonal elements in each row, to serve as null distributions of the network topological features. With this permutation, the total number of edges in the network remains the same and the numbers of edges coming into each node remains the same, but the sources for the edges are uniformly randomly assigned with each permutation.

## Contributions

TEG: Gnotobiotic study design. Statistical model. Inference Algorithm. Software. Data analysis. Writing. Reviewing.

YK: Software. Data analysis. Writing methods.

SA: Software. Data analysis. Writing methods.

DEK: Software. Writing methods.

ND: Gnotobiotic study design and experiments.

RL: Gnotobiotic study design.

BB: Discussions with YK and critical review of the manuscript

JRA: Human donor sample collection and data acquisition. Critical review of the manuscript.

LB: Gnotobiotic study design and experiments. Detailed review of the manuscript.

GKG: Project conception. Gnotobiotic study design. Statistical model. Inference Algorithm. Software. Data analysis. Writing. Reviewing. Project management.

## Competing Interests

No industry support was provided for this study.

TEG: None.

YK: None.

SA: None.

DK: None.

ND: None.

RL: None.

BB: None.

JRA consults for Finch Therapeutics, BMS, Pfizer, Janssen, Morphic, Iterative Scopes, Artugen, Servatus, Pandion, Merck, Baccain and has research support from Merck. No industry support was provided for this study.

LB is the inventor of patents for defined bacterial therapeutics for *C. difficile,* and is the SAB chair and a shareholder in ParetoBio, Inc. No industry support was provided for this study.

GKG is a shareholder in Kaleido Biosciences, Inc., and is on the SAB and is a shareholder in ParetoBio, Inc. His interests were reviewed and are managed by Brigham and Women’s Hospital and Mass General Brigham in accordance with their conflict-of-interest policies.

### Funding

Research reported in this publication was supported by DARPA BRICS HR0011-15-C-0094 (Gerber and Bry), NIH R01GM130777 (Gerber), NIH R35GM143056 (Gibson), NIH R35GM141861 (Berger), NIH P30DK056338 (Bry), NSF MTM2 2025512 (Gerber), the BWH President’s Scholar Award (Gerber)

## Software and Data Availability

First time users wanting to explore the model and the data should start by visiting the repo for this paper that reproduces figures and contains notebooks for exploring the model https://github.com/gerberlab/MDSINE2_Paper. The core MDSINE2 package is available at https://github.com/gerberlab/MDSINE2. All sequencing data for this study are available at https://www.ncbi.nlm.nih.gov/bioproject/PRJNA784519.

---

**Supplemental Figure 1:**
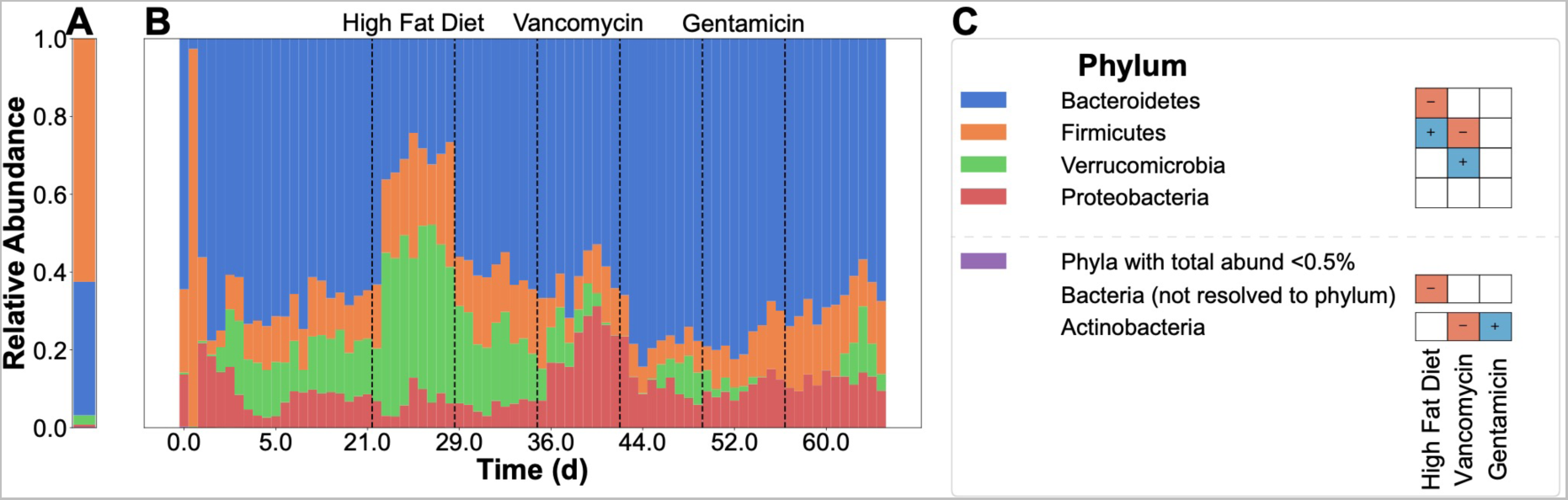
Phylum level grouping of reads for the high-temporal resolution gnotobiotic mouse study. **(A)** Relative abundance of microbes in human donor sample. **(B)** Relative abundances of microbes in serial fecal samples, averaged over the biological replicates. **(C)** Legend and taxonomy for panels A and B along with significant differential abundances for each taxonomic group across the three perturbations (see Supplemental Table 1 for *p*-values).

**Supplemental Figure 2:**
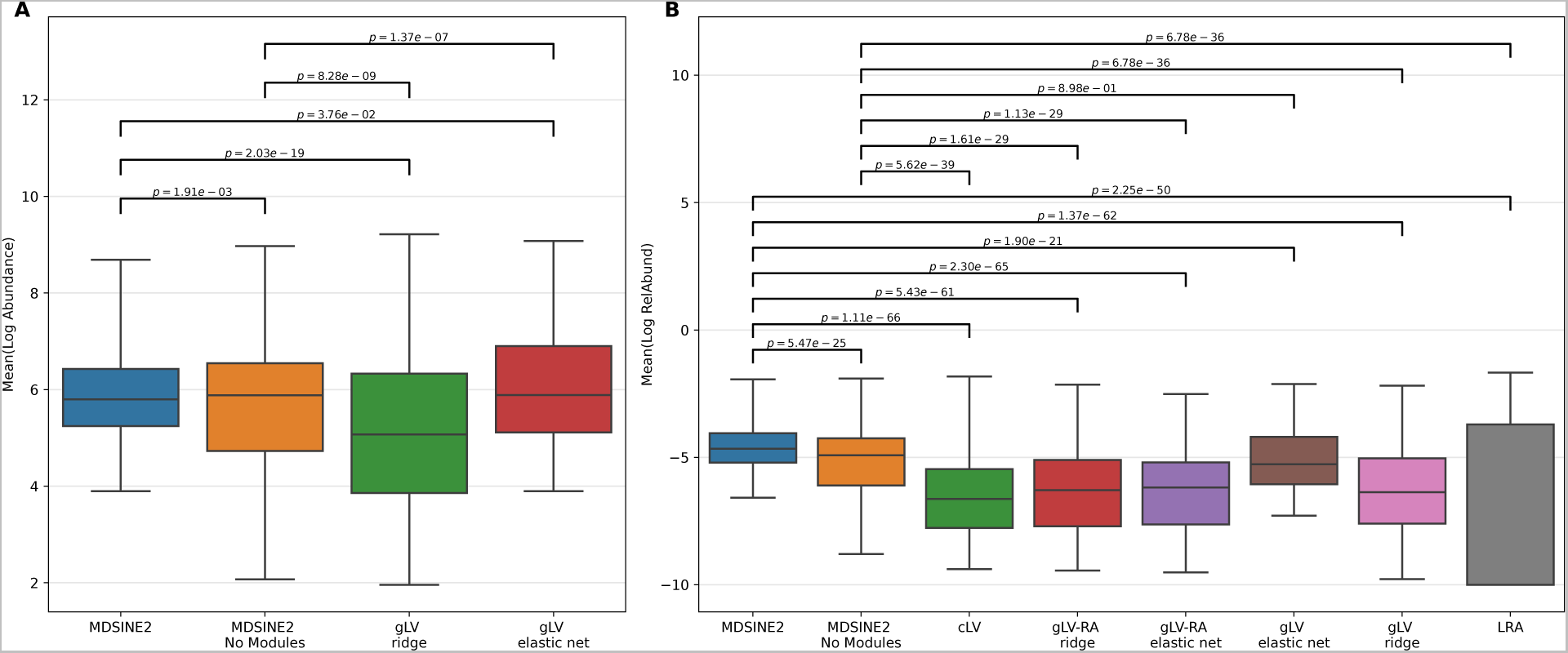
All models have relatively poor performance for the zero read taxa samples with errors on the order of the limit of detection. Results are for cross-fold validation of forecasts. Models were trained on three of four mice, then given the initial condition of the hold out mouse time-series, and then the inferred model was simulated forwards and the output was compared to the entire held out time-series. All tests are two-sided Wilcoxon signed-rank test (paired non-parametric). Non-significant tests are denoted with red bracket. Errors are calculated from forecasted abundance with respect to hold out data across all folds and for all taxa with zero reads within each sample. **(A)** Forecasting error for all zero taxa samples for models trained on concentrations. **(B)** Forecasting error for all zero taxa samples for models trained on relative abundances.

**Supplemental Figure 3:**
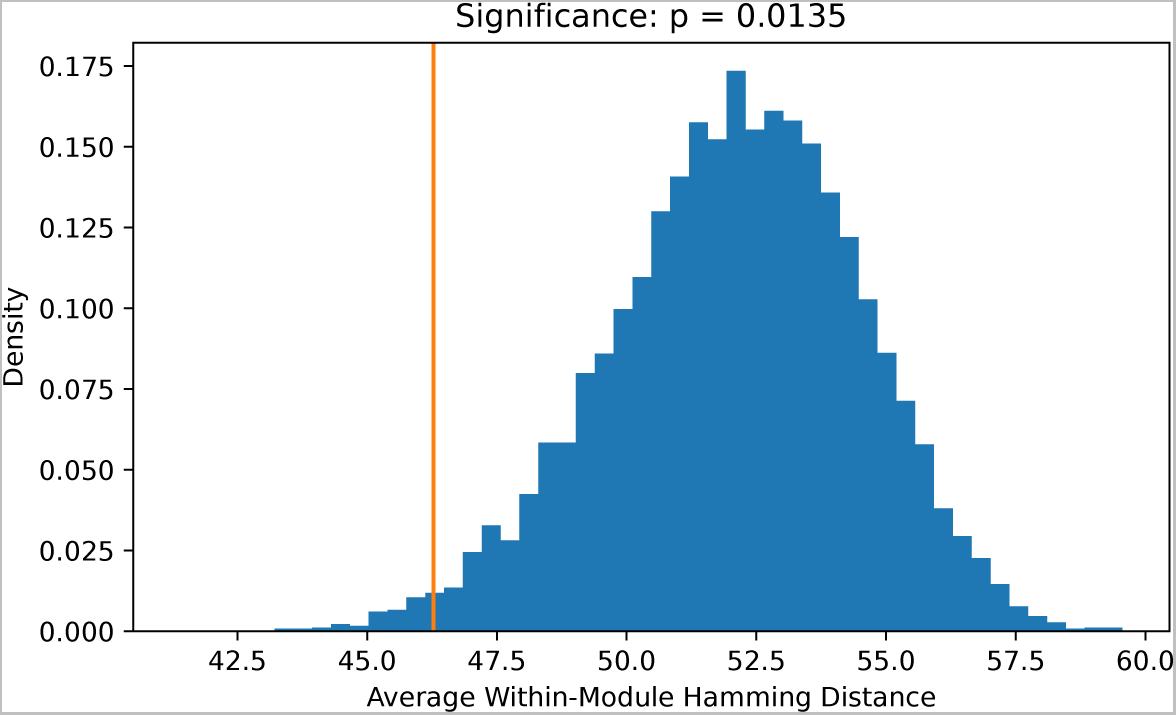
Phylogenetic distance of modules. Average phylogenetic distance of modules shown by orange line, compared to random permutations of taxa in modules for null distribution in blue. Permutation test results in p=0.0135.

**Supplemental Figure 4:**
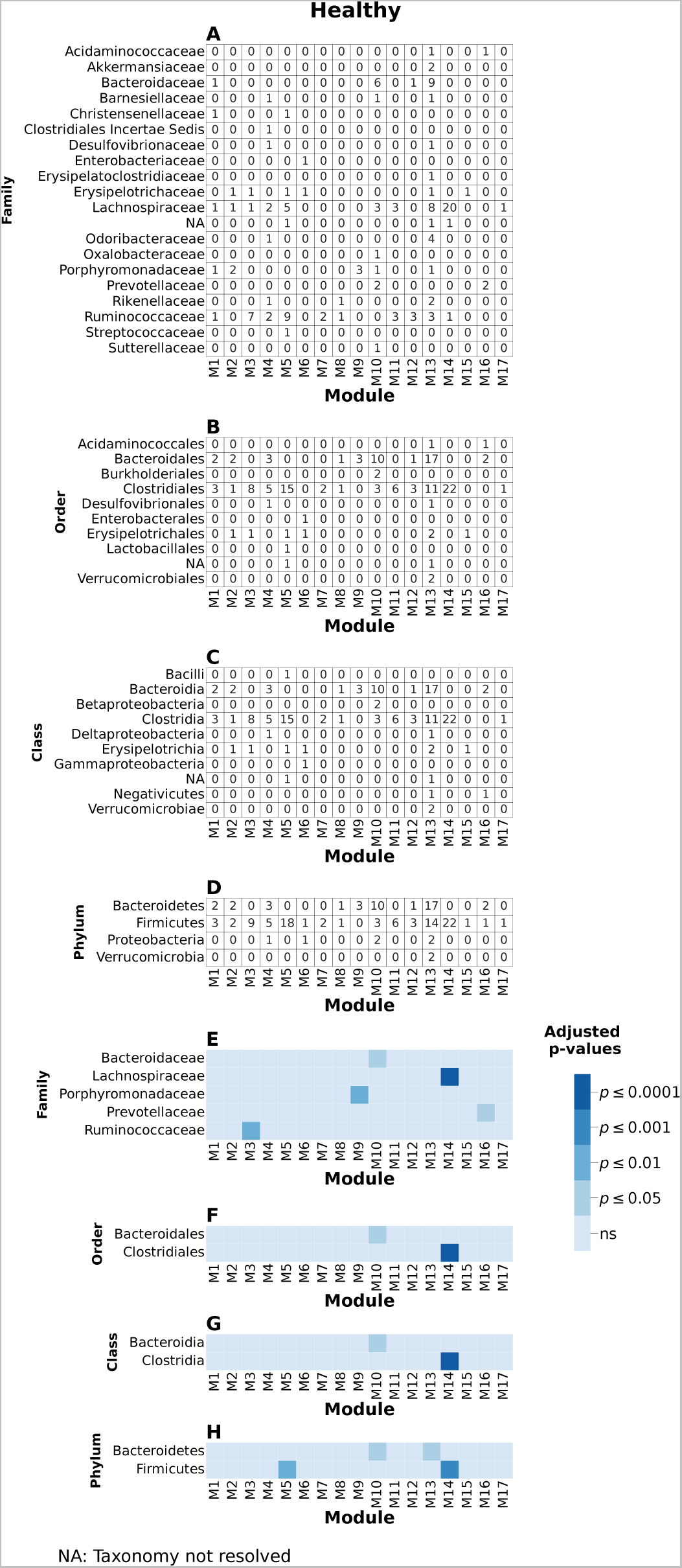
Taxonomic enrichment. Enrichment for the modules was performed at Phylum, Class, Order, and Family levels using the hypergeometric test, Methods §4.5.

**Supplemental Figure 5.**
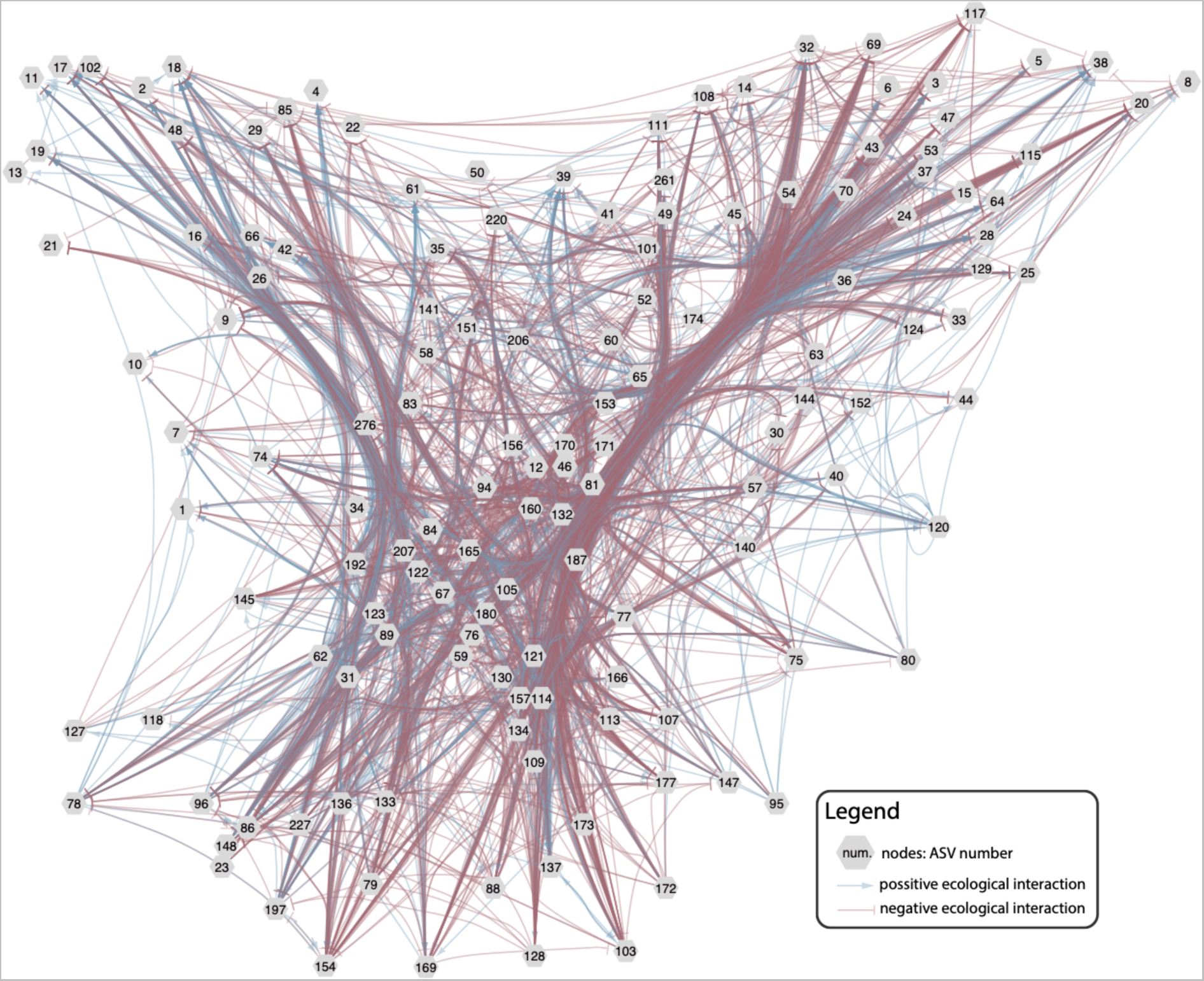
Taxon-taxon interaction network for model trained without module inference. Interaction network displaying only edges with BF > 100 (*decisive* evidence).

**Supplemental Figure 6:**
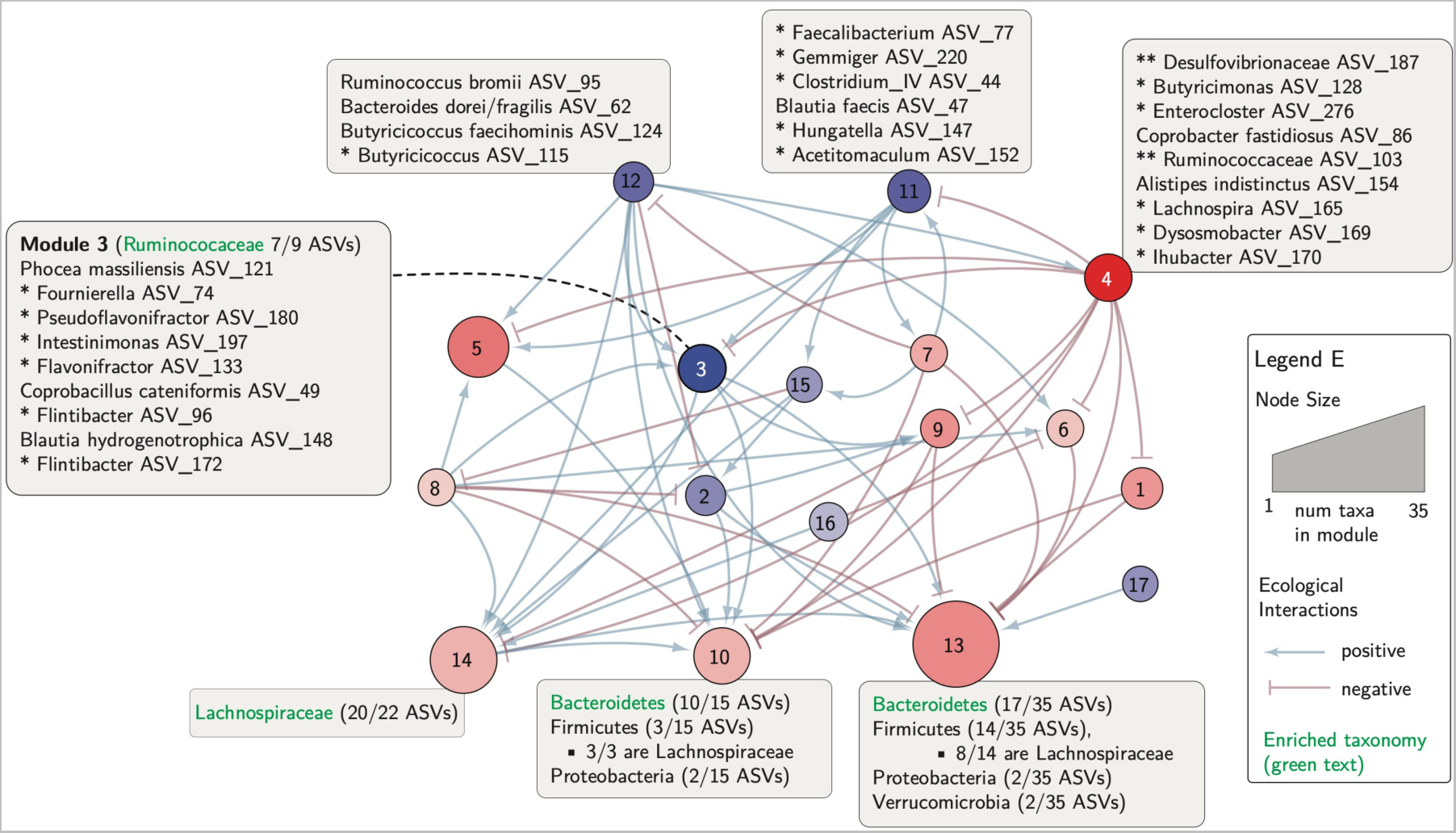
Annotated modules from Figure 5C. Taxa from specific modules denoted by ASVs along with taxonomic breakdown at higher levels. Enriched taxonomies in green. Only to aid in discussion, not intended to be comprehensive.

**Supplemental Table 1:**
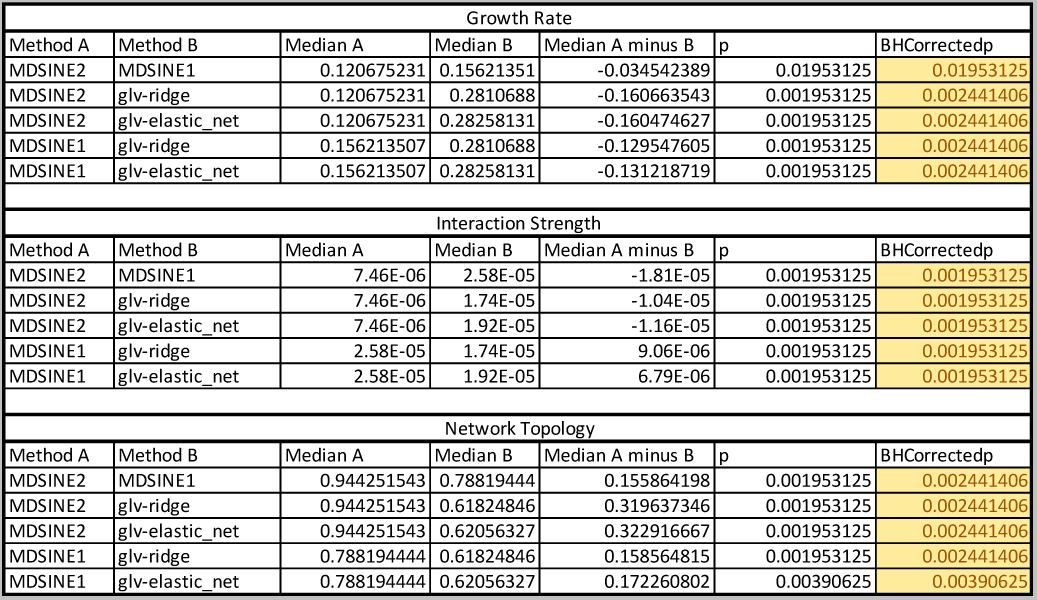
*p*-values for Figure 2.

**Supplemental Table 2:**
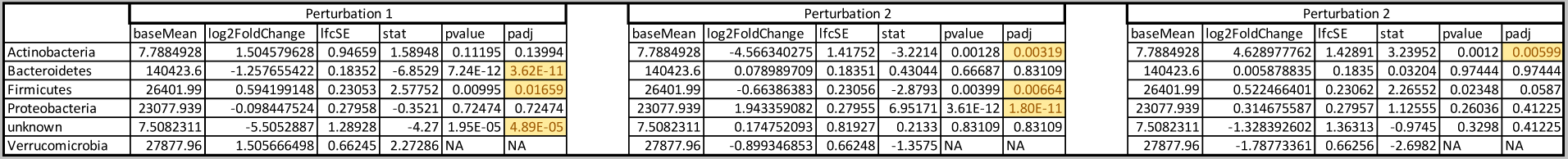
*p*-values for Supplemental Figure 1C.

**Supplemental Table 3:**
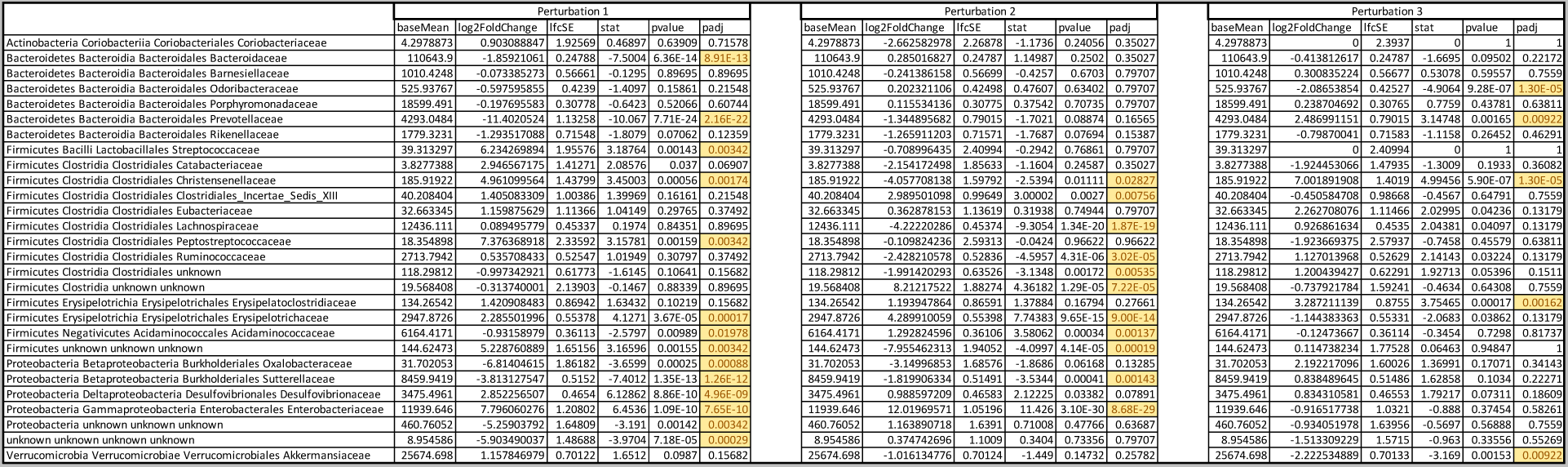
*p*-values for Figure 3B.

**Supplemental Table 4:**
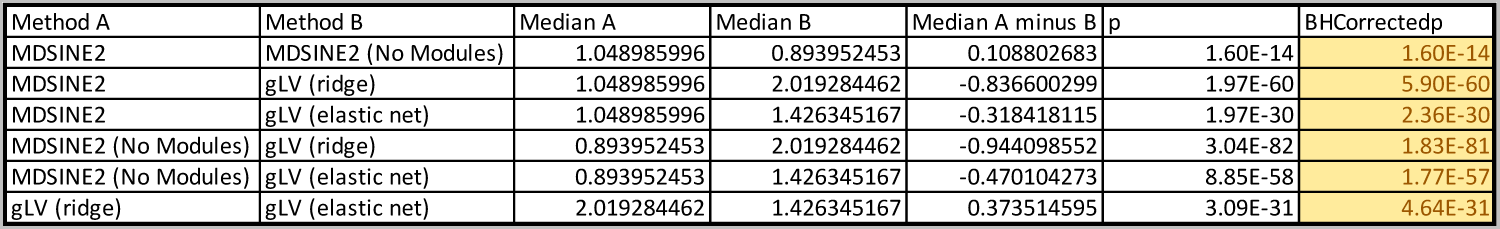
*p*-values associated with Figure 4A.

**Supplemental Table 5:**
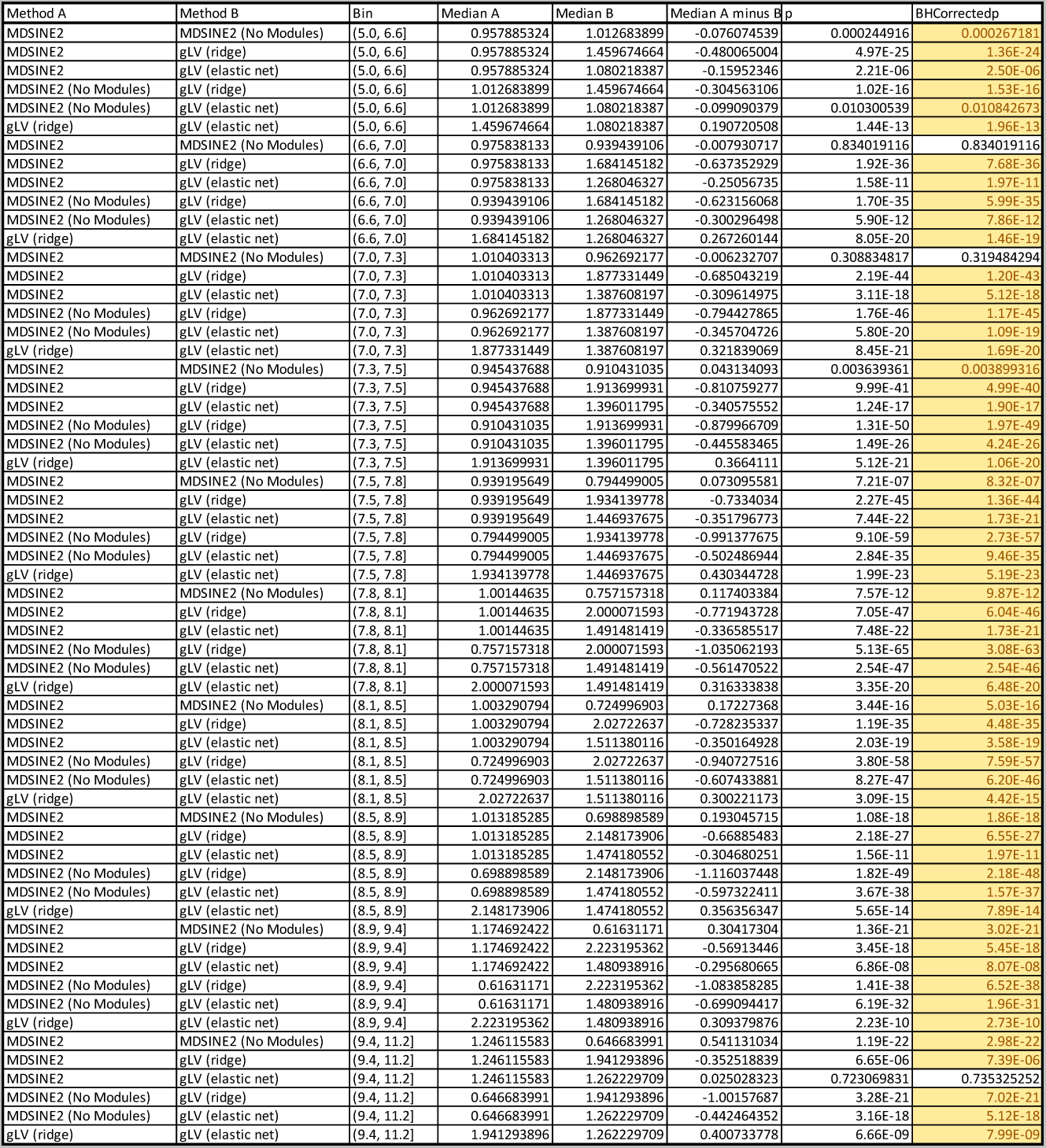
*p*-values associated with Figure 4C.

**Supplemental Table 6:** *p*-values associated with Figure 4D.

| Method A | Method B | Median A | Median B | Median A minus B | p | BHCorrectedp |
| --- | --- | --- | --- | --- | --- | --- |
| MDSINE2 | MDSINE2 (No Modules) | 0.957595993 | 0.8525636 | 0.081819895 | 8.80E-09 | 1.12E-08 |
| MDSINE2 | gLV (ridge) | 0.957595993 | 2.406869401 | -1.355265884 | 5.77E-82 | 4.04E-81 |
| MDSINE2 | gLV (elastic net) | 0.957595993 | 1.576258627 | -0.516491597 | 4.17E-57 | 9.72E-57 |
| MDSINE2 | gLV-RA (ridge) | 0.957595993 | 2.851254421 | -1.780842 | 7.15E-80 | 2.86E-79 |
| MDSINE2 | gLV-RA (elastic net) | 0.957595993 | 3.148218679 | -2.076211147 | 1.13E-80 | 5.26E-80 |
| MDSINE2 | cLV | 0.957595993 | 2.440354583 | -1.330361648 | 2.65E-78 | 9.27E-78 |
| MDSINE2 | LRA | 0.957595993 | 5.507975998 | -4.174724893 | 5.03E-71 | 1.41E-70 |
| MDSINE2 (No Modules) | gLV (ridge) | 0.8525636 | 2.406869401 | -1.406978101 | 1.57E-87 | 4.41E-86 |
| MDSINE2 (No Modules) | gLV (elastic net) | 0.8525636 | 1.576258627 | -0.618572159 | 1.53E-65 | 3.90E-65 |
| MDSINE2 (No Modules) | gLV-RA (ridge) | 0.8525636 | 2.851254421 | -1.871503505 | 5.07E-85 | 7.10E-84 |
| MDSINE2 (No Modules) | gLV-RA (elastic net) | 0.8525636 | 3.148218679 | -2.127072799 | 8.70E-83 | 8.12E-82 |
| MDSINE2 (No Modules) | cLV | 0.8525636 | 2.440354583 | -1.522558093 | 9.59E-81 | 5.26E-80 |
| MDSINE2 (No Modules) | LRA | 0.8525636 | 5.507975998 | -4.097417397 | 9.75E-73 | 3.03E-72 |
| gLV (ridge) | gLV (elastic net) | 2.406869401 | 1.576258627 | 0.619909899 | 2.68E-44 | 4.69E-44 |
| gLV (ridge) | gLV-RA (ridge) | 2.406869401 | 2.851254421 | -0.162335126 | 0.000122049 | 0.00014239 |
| gLV (ridge) | gLV-RA (elastic net) | 2.406869401 | 3.148218679 | -0.386976559 | 9.43E-07 | 1.15E-06 |
| gLV (ridge) | cLV | 2.406869401 | 2.440354583 | -0.101756525 | 0.262318692 | 0.262318692 |
| gLV (ridge) | LRA | 2.406869401 | 5.507975998 | -1.750185431 | 1.03E-25 | 1.60E-25 |
| gLV (elastic net) | gLV-RA (ridge) | 1.576258627 | 2.851254421 | -1.190920049 | 2.84E-48 | 5.30E-48 |
| gLV (elastic net) | gLV-RA (elastic net) | 1.576258627 | 3.148218679 | -1.407795771 | 1.53E-48 | 3.06E-48 |
| gLV (elastic net) | cLV | 1.576258627 | 2.440354583 | -0.639269082 | 3.77E-30 | 6.21E-30 |
| gLV (elastic net) | LRA | 1.576258627 | 5.507975998 | -3.18934168 | 1.09E-52 | 2.34E-52 |
| gLV-RA (ridge) | gLV-RA (elastic net) | 2.851254421 | 3.148218679 | -0.077225254 | 0.145522958 | 0.156717031 |
| gLV-RA (ridge) | cLV | 2.851254421 | 2.440354583 | 0.06285764 | 0.035139015 | 0.039355697 |
| gLV-RA (ridge) | LRA | 2.851254421 | 5.507975998 | -1.498749843 | 7.08E-15 | 9.91E-15 |
| gLV-RA (elastic net) | cLV | 3.148218679 | 2.440354583 | -0.181748402 | 0.201459803 | 0.208921277 |
| gLV-RA (elastic net) | LRA | 3.148218679 | 5.507975998 | -1.191689473 | 6.19E-12 | 8.26E-12 |
| cLV | LRA | 2.440354583 | 5.507975998 | -1.539773639 | 1.15E-22 | 1.69E-22 |

**Supplemental Table 7:**
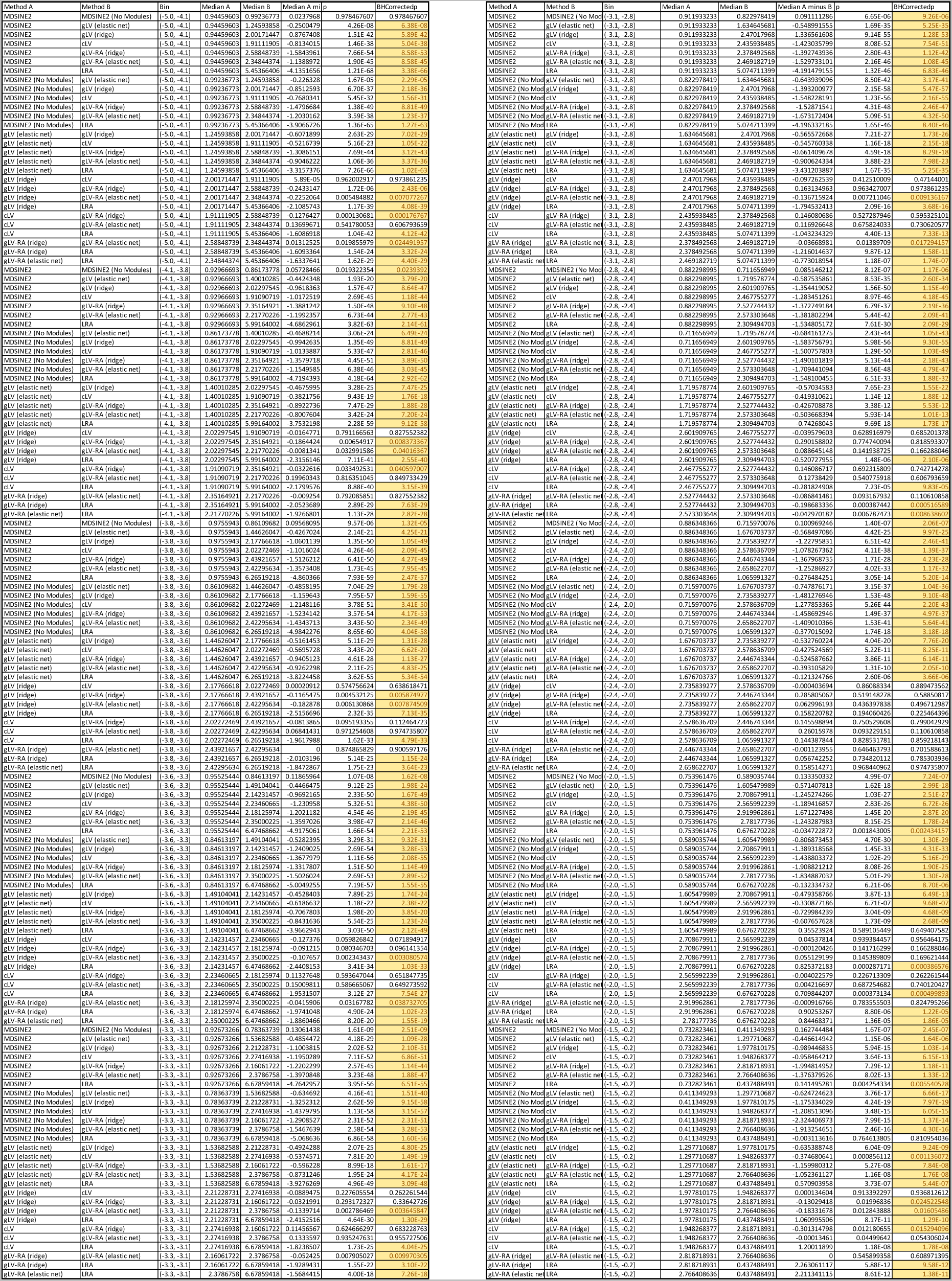
*p*-values associated with Figure 4F.

**Supplemental Table 8:**
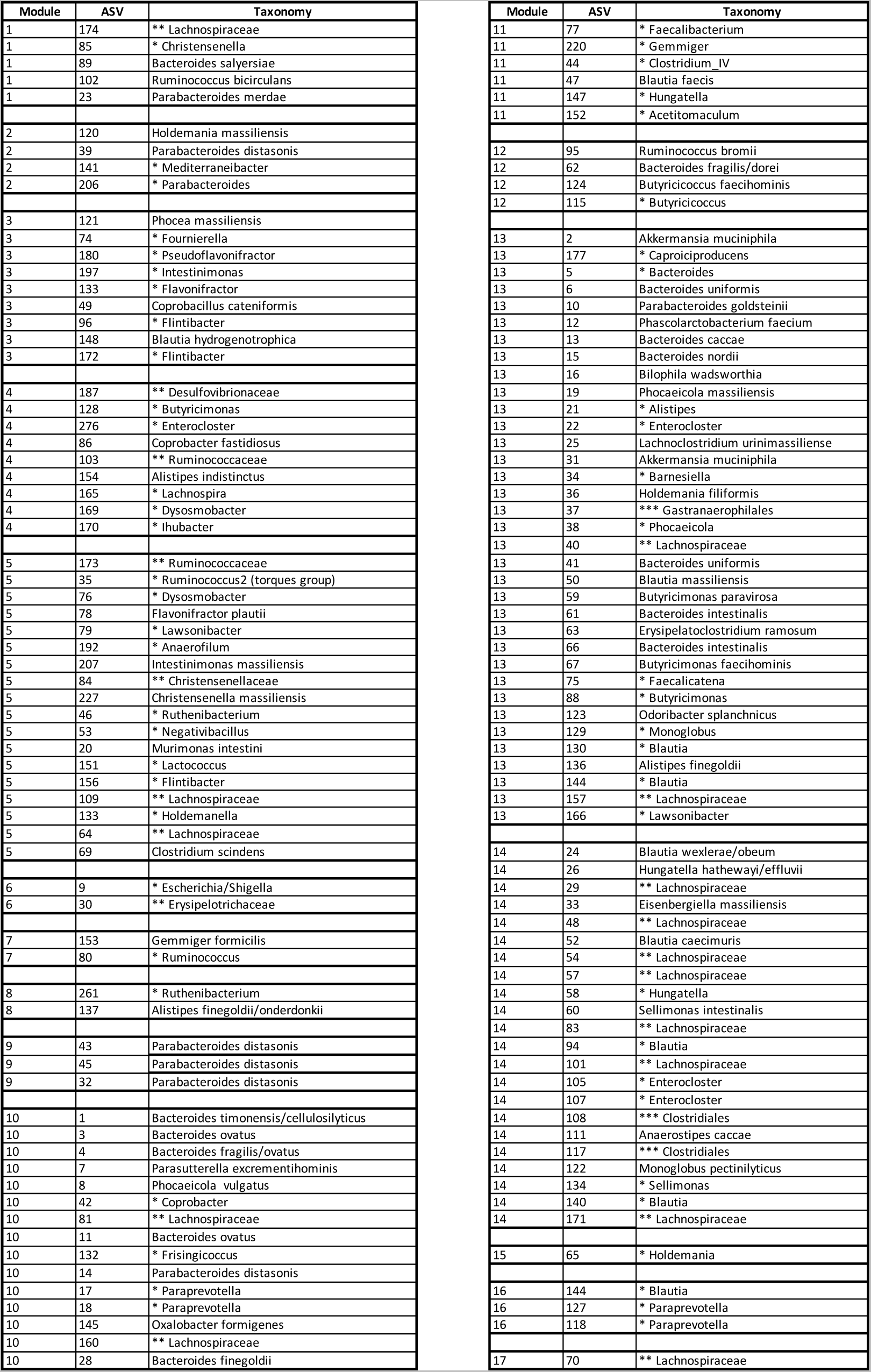
Module members. Taxa comprising each module along with ASV number.

## Notes

https://github.com/gerberlab/MDSINE2

https://github.com/gerberlab/MDSINE2_Paper

### Competing Interest Statement

The authors have declared no competing interest.

### Summary of Updates

Revised to focus on the bayesian model

## References

1 Bucci, V., et al. MDSINE: Microbial Dynamical Systems INference Engine for microbiome time-series analyses. Genome Biol 17 (2016). https://doi.org:10.1186/s13059-016-0980-6

2 Gerber, G. K. The dynamic microbiome. FEBS letters 588, 4131–4139 (2014).

3 Gilbert, J. A., et al. Current understanding of the human microbiome. Nature Medicine 24, 392–400 (2018). https://doi.org:10.1038/nm.4517

4 Gonze, D., Coyte, K. Z., Lahti, L. & Faust, K. Microbial communities as dynamical systems. Current opinion in microbiology 44, 41–49 (2018).

5 May, R. M. Will a large complex system be stable? Nature 238, 413–414 (1972).

6 Goh, B. S. Global Stability in Many-Species Systems. The American Naturalist 111, 135–143 (1977).

7 Coyte, K. Z., Schluter, J. & Foster, K. R. The ecology of the microbiome: networks, competition, and stability. Science 350, 663–666 (2015).

8 Rao, C., et al. Multi-kingdom ecological drivers of microbiota assembly in preterm infants. Nature 591, 633–638 (2021). https://doi.org:10.1038/s41586-021-03241-8

9 Lee, T. I., et al. Transcriptional regulatory networks in Saccharomyces cerevisiae. science 298, 799–804 (2002).

10 Milo, R., et al. Network motifs: simple building blocks of complex networks. Science 298, 824–827 (2002).

11 Barabasi, A.-L. & Oltvai, Z. N. Network biology: understanding the cell’s functional organization. Nature reviews genetics 5, 101–113 (2004).

12 Franzosa, E. A., et al. Gut microbiome structure and metabolic activity in inflammatory bowel disease. Nature Microbiology 4, 293–305 (2019). https://doi.org:10.1038/s41564-018-0306-4

13 Stein, R. R., et al. Ecological Modeling from Time-Series Inference: Insight into Dynamics and Stability of Intestinal Microbiota. PLoS Comput Biol 9 (2013).

14 Fisher, C. K. & Mehta, P. Identifying Keystone Species in the Human Gut Microbiome from Metagenomic Timeseries Using Sparse Linear Regression. PLoS ONE 9, e102451 (2014).

15 Joseph, T. A., Shenhav, L., Xavier, J. B., Halperin, E. & Pe’er, I. Compositional Lotka-Volterra describes microbial dynamics in the simplex. PLOS Computational Biology 16, e1007917 (2020). https://doi.org:10.1371/journal.pcbi.1007917

16 Thanissery, R., Winston, J. A. & Theriot, C. M. Inhibition of spore germination, growth, and toxin activity of clinically relevant C. difficile strains by gut microbiota derived secondary bile acids. Anaerobe 45, 86–100 (2017).

17 Ze, X., Duncan, S. H., Louis, P. & Flint, H. J. Ruminococcus bromii is a keystone species for the degradation of resistant starch in the human colon. The ISME Journal 6, 1535–1543 (2012). https://doi.org:10.1038/ismej.2012.4

18 Cao, H. T., Gibson, T. E., Bashan, A. & Liu, Y. Y. Inferring human microbial dynamics from temporal metagenomics data: Pitfalls and lessons. Bioessays 39 (2017). https://doi.org:10.1002/bies.201600188

19 Gerber, G. K., Onderdonk, A. B. & Bry, L. Inferring dynamic signatures of microbes in complex host ecosystems. PLoS Comput Biol 8, e1002624 (2012). https://doi.org:10.1371/journal.pcbi.1002624

20 Creswell, R., et al. High-resolution temporal profiling of the human gut microbiome reveals consistent and cascading alterations in response to dietary glycans. Genome Medicine 12, 59 (2020). https://doi.org:10.1186/s13073-020-00758-x

21 Simberloff, D. & Dayan, T. The Guild Concept and the Structure of Ecological Communities. Annual Review of Ecology and Systematics 22, 115–143 (1991). https://doi.org:10.1146/annurev.es.22.110191.000555

22 Hofman, J. M. & Wiggins, C. H. Bayesian approach to network modularity. Physical review letters 100, 258701 (2008).

23 Segal, E., Friedman, N., Kaminski, N., Regev, A. & Koller, D. From signatures to models: understanding cancer using microarrays. Nature genetics 37, S38–S45 (2005).

24 Gibson, T. & Gerber, G. in Proceedings of the 35th International Conference on Machine Learning Vol. 80 1763–1772 (2018).

25 Lavin, R., DiBenedetto, N., Yeliseyev, V., Delaney, M. & Bry, L. Gnotobiotic and Conventional Mouse Systems to Support Microbiota Based Studies. Curr Protoc Immunol 121, e48 (2018). https://doi.org:10.1002/cpim.48

26 Faith, J. J., Ahern, P. P., Ridaura, V. K., Cheng, J. & Gordon, J. I. Identifying gut microbe–host phenotype relationships using combinatorial communities in gnotobiotic mice. Science translational medicine 6, 220ra211-220ra211 (2014).

27 Planer, J. D., et al. Development of the gut microbiota and mucosal IgA responses in twins and gnotobiotic mice. Nature 534, 263–266 (2016).

28 Callahan, B. J., et al. DADA2: High-resolution sample inference from Illumina amplicon data. Nat Methods 13, 581–583 (2016). https://doi.org:10.1038/nmeth.3869

29 Love, M. I., Huber, W. & Anders, S. Moderated estimation of fold change and dispersion for RNA-seq data with DESeq2. Genome Biology 15, 550 (2014). https://doi.org:10.1186/s13059-014-0550-8

30 McMurdie, P. J. & Holmes, S. Waste not, want not: why rarefying microbiome data is inadmissible. PLoS computational biology 10, e1003531 (2014).

31 Greenacre, M. Compositional data analysis. Annual Review of Statistics and its Application 8, 271–299 (2021).

32 Kass, R. E. &Raftery, A. E. Bayes Factors. Journal of the American Statistical Association 90, 773–795 (1995). https://doi.org:10.1080/01621459.1995.10476572

33 Carmody, R. N., et al. Diet dominates host genotype in shaping the murine gut microbiota. Cell Host Microbe 17, 72–84 (2015). https://doi.org:10.1016/j.chom.2014.11.010

34 Paine, R. T. A note on trophic complexity and community stability. The American Naturalist 103, 91–93 (1969).

35 Banerjee, S., Schlaeppi, K. & van der Heijden, M. G. Keystone taxa as drivers of microbiome structure and functioning. Nature Reviews Microbiology 16, 567–576 (2018).

36 Röttjers, L. & Faust, K. Can we predict keystones? Nature Reviews Microbiology 17, 193–193 (2019). https://doi.org:10.1038/s41579-018-0132-y

37 Faith, J. J., et al. The Long-Term Stability of the Human Gut Microbiota. Science 341, 1237439 (2013). https://doi.org:10.1126/science.1237439

38 Lozupone, C. A., Stombaugh, J. I., Gordon, J. I., Jansson, J. K. & Knight, R. Diversity, stability and resilience of the human gut microbiota. Nature 489, 220–230 (2012). https://doi.org:10.1038/nature11550

39 Dethlefsen, L. & Relman, D. A. Incomplete recovery and individualized responses of the human distal gut microbiota to repeated antibiotic perturbation. Proceedings of the National Academy of Sciences 108, 4554–4561 (2011). https://doi.org:10.1073/pnas.1000087107

40 Gibson, T. E., Bashan, A., Cao, H.-T., Weiss, S. T. & Liu, Y.-Y. On the Origins and Control of Community Types in the Human Microbiome. PLOS Computational Biology 12, e1004688 (2016). https://doi.org:10.1371/journal.pcbi.1004688

41 Doyle, J. C., Francis, B. A. & Tannenbaum, A. R. Feedback control theory. (Courier Corporation, 2013).

42 Gibson, T. E. Sign Stability via Root Locus Analysis. arXiv preprint arXiv:*1512.06026* (2015).

43 Allesina, S. & Tang, S. Stability criteria for complex ecosystems. Nature 483, 205--208 (2012).

44 Sommers, H. J., Crisanti, A., Sompolinsky, H. & Stein, Y. Spectrum of large random asymmetric matrices. Physical Review Letters 60, 1895--1898 (1988).

45 Palmer, J. D. & Foster, K. R. Bacterial species rarely work together. Science 376, 581–582 (2022).

46 Gelman, A. et al. Bayesian data analysis. (Chapman and Hall/CRC, 2013).

47 Bar-Joseph, Z., et al. Computational discovery of gene modules and regulatory networks. Nature Biotechnology 21, 1337–1342 (2003). https://doi.org:10.1038/nbt890

48 Gerber, G. K. Computational discovery of gene modules, regulatory networks and expression programs PhD thesis, Harvard and MIT Division of Health Sciences and Technology, (2007).

49 Gerber, G. K., Dowell, R. D., Jaakkola, T. S. & Gifford, D. K. Automated discovery of functional generality of human gene expression programs. PLoS Comput Biol 3, e148 (2007). https://doi.org:10.1371/journal.pcbi.0030148

50. Gibson, T. E. in American Control Conference (IEEE, 2016).

51 Gibson, T. E., Bashan, A., Cao, H. T., Weiss, S. T. & Liu, Y. Y. On the Origins and Control of Community Types in the Human Microbiome. PLoS Comput Biol 12, e1004688 (2016). https://doi.org:10.1371/journal.pcbi.1004688

52 Angulo, M. T., Moog, C. H. & Liu, Y.-Y. A theoretical framework for controlling complex microbial communities. Nature Communications 10, 1045 (2019). https://doi.org:10.1038/s41467-019-08890-y

53 Baxter, N. T., et al. Dynamics of Human Gut Microbiota and Short-Chain Fatty Acids in Response to Dietary Interventions with Three Fermentable Fibers. mBio 10, e02566–02518 (2019). https://doi.org:doi:10.1128/mBio.02566-18

54 Furusawa, Y., et al. Commensal microbe-derived butyrate induces the differentiation of colonic regulatory T cells. Nature 504, 446–450 (2013).

55 Joossens, M., et al. Dysbiosis of the faecal microbiota in patients with Crohn&#039;s disease and their unaffected relatives. Gut 60, 631 (2011). https://doi.org:10.1136/gut.2010.223263

56 Kowalska-Duplaga, K., et al. Differences in the intestinal microbiome of healthy children and patients with newly diagnosed Crohn’s disease. Sci Rep 9, 18880 (2019). https://doi.org:10.1038/s41598-019-55290-9

57 Forbes, J. D., et al. A comparative study of the gut microbiota in immune-mediated inflammatory diseases—does a common dysbiosis exist? Microbiome 6, 221 (2018). https://doi.org:10.1186/s40168-018-0603-4

58 Ziesack, M., et al. Engineered interspecies amino acid cross-feeding increases population evenness in a synthetic bacterial consortium. Msystems 4, e00352–00319

59 Neal, R. M. Markov chain sampling methods for Dirichlet process mixture models. Journal of computational and graphical statistics 9, 249–265 (2000).

60 Antoniak, C. E. Mixtures of Dirichlet processes with applications to Bayesian nonparametric problems. The annals of statistics, 1152-1174 (1974).

61 Escobar, M. D. & West, M. Bayesian density estimation and inference using mixtures. Journal of the american statistical association 90, 577–588 (1995).

62 Harris, C. R., et al. Array programming with NumPy. Nature 585, 357–362 (2020).

63 Virtanen, P., et al. SciPy 1.0: fundamental algorithms for scientific computing in Python. Nature methods 17, 261–272 (2020).

64 Lam, S. K., Pitrou, A. & Seibert, S. in Proceedings of the Second Workshop on the LLVM Compiler Infrastructure in HPC. 1–6.

65 Hunter, J. D. Matplotlib: A 2D graphics environment. Computing in science & engineering 9, 90–95 (2007).

66 Waskom, M. & team, S. d. mwaskom/seaborn. (2020). <https://doi.org/10.5281/zenodo.592845>.

67 Shannon, P., et al. Cytoscape: a software environment for integrated models of biomolecular interaction networks. Genome Res 13, 2498–2504 (2003).

68 Hsu, B. B., et al. Dynamic Modulation of the Gut Microbiota and Metabolome by Bacteriophages in a Mouse Model. Cell Host Microbe 25, 803–814 e805 (2019). https://doi.org:10.1016/j.chom.2019.05.001

69 Kozich, J. J., Westcott, S. L., Baxter, N. T., Highlander, S. K. & Schloss, P. D. Development of a dual-index sequencing strategy and curation pipeline for analyzing amplicon sequence data on the MiSeq Illumina sequencing platform. Applied and environmental microbiology 79, 5112–5120 (2013).

70 Cole, J. R., et al. The Ribosomal Database Project: improved alignments and new tools for rRNA analysis. Nucleic Acids Res 37, D141–D145 (2009).

71 Nawrocki, E. P., Kolbe, D. L. & Eddy, S. R. Infernal 1.0: inference of RNA alignments. Bioinformatics 25, 1335–1337 (2009).

72 Price, M. N., Dehal, P. S. & Arkin, A. P. FastTree 2–approximately maximum-likelihood trees for large alignments. PloS one 5, e9490 (2010).

73 Eddy, S. R. Accelerated profile HMM searches. PLoS computational biology 7, e1002195 (2011).

74 Matsen, F. A., Kodner, R. B. & Armbrust, E. V. pplacer: linear time maximum-likelihood and Bayesian phylogenetic placement of sequences onto a fixed reference tree. BMC bioinformatics 11, 1–16 (2010).

