## Supplemental Text for "Microbial dynamics inference at ecosystem-scale"

### Contents


1 Description of Model S.14
- 1.1 Model of dynamics . . . S.14
- 1.2 Growth and self-limiting variables . . . S.15
  - 1.2.1 Growth and self-limiting hyperparameters . . . S.16
- 1.3 Dirichlet process prior on interaction modules . . . S.17
  - 1.3.1 Concentration parameter hyperparameters . . . S.17
  - 1.3.2 Connections between interaction modules and prior work . . . S.17
- 1.4 Interaction variables . . . S.17
  - 1.4.1 Interaction hyperparameters . . . S.18
- 1.5 Perturbation effects . . . S.18
  - 1.5.1 Perturbation hyperparameters . . . S.19
- 1.6 Process variance . . . S.19
  - 1.6.1 Process variance hyperparameters . . . S.19
- 1.7 Measurement error model . . . S.19

2 Model inference S.20
- 2.1 Sampling perturbation and interaction strengths and their priors . . . S.21
- 2.2 Sampling growth rates, self-interaction strengths and parameters of their priors . S.21
- 2.3 Sampling process variance . . . S.22
- 2.4 Sampling indicators variables and their priors . . . S.22
- 2.5 Sampling latent trajectories . . . S.23
- 2.6 Sampling module assignments and DP priors . . . S.23
- 2.7 Mixing . . . S.24

3 Extended discussion S.24
- 3.1 Stability . . . S.24

Appendix A Probability density functions S.25

Appendix B Discretizing GLV dynamics S.26

Appendix C Stick-breaking process S.26

Appendix D Bayesian regression: posterior and marginalization S.26

Appendix E Estimating negative binomial dispersion parameters from data S.28

Appendix F Metropolis-Hastings Proposal Tuning S.29

Appendix G Bayes Factor Computation S.29

**Table 1:** Model parameters and variables

| variable | space | description |
|---|---|---|
| $\alpha$ | $\mathbb{R}^{>0}$ | Concentration parameter for stick breaking process generating $\pi_{\mathbf{c}}$ in (21). Prior defined in (20). |
| $\mathbf{a}_{1,i}$ | $\mathbb{R}^{>0}$ | Growth rate of taxa $i$ for the dynamics in (2)-(3). Prior defined in (6). |
| $\mathbf{a}_{2,i}$ | $\mathbb{R}^{>0}$ | Self limiting term of taxa $i$ for the dynamics in (2)-(3). Prior defined in (9). |
| $\mathbf{b}_{\mathbf{c}_i,\mathbf{c}_j}$ | $\mathbb{R}$ | Interaction parameter for the strength of interaction from module $\mathbf{c}_j$ on $\mathbf{c}_i$ for the dynamics in (2)-(3). Prior defined in (28). |
| $b_{1,\mathbf{b}}, b_{2,\mathbf{b}}$ | $\mathbb{R}^{>0}$ | Shape hyperparameters for $\pi_{\mathbf{b}}$ in (29). Values defined (35)-(36). |
| $b_{1,\gamma}$, $b_{2,\gamma}$ | $\mathbb{R}^{>0}$ | Shape hyperparameters for $\pi_{\gamma,p}$ in (40). Values defined in (46) |
| $\mathbf{c}_i$ | $\mathbb{Z}^{+}$ | Module assignment for taxa $i$. Prior defined in (22) |
| $d_0, d_1$ | | Offset and slope parameters for negative binomial dispersion, estimated from sequencing replicates (see Appendix E). |
| $\Delta_{s,k}$ | $\mathbb{R}^{>0}$ | Time between samples $k+1$ and $k$ in subject $s$ defined as $\Delta_{s,k} = t_{s,k+1} - t_{s,k}$ |
| $\epsilon_{s,i}$ | $\mathbb{R}^{>0}$ | Dispersion parameter for sequencing read counts $\mathbf{y}_{s,i}(k)$ in (50). Parameterization defined in (52) |
| $\gamma_{\mathbf{c}_i,p}$ | $\mathbb{R}$ | Perturbation strength parameter for perturbation $p$ on module $\mathbf{c}_i$ for the dynamics in (2)-(3). Prior defined in (39). |
| $h_{s,p}(k)$ | $\{0,1\}$ | Perturbation indicator for when perturbation $p$ is active in subject $s$ for the dynamics in (2)-(3). |
| $i, j$ | $\mathbb{Z}^{+}$ | Taxa indices. |
| $k$ | $\mathbb{Z}$ | Discrete time index. |
| $\ell, m$ | $\mathbb{Z}^{>0}$ | Used to index over modules. |
| $\mu_{\mathbf{a}_1}$ | $\mathbb{R}^{>0}$ | Location parameter for $\mathbf{a}_{1,i}$ in (6). Prior defined in (4). |
| $\mu_{\mathbf{a}_2}$ | $\mathbb{R}^{>0}$ | Location parameter for $\mathbf{a}_{2,i}$ in (9). Prior defined in (7). |
| $\mu_{0,\mathbf{a}_1}$ | $\mathbb{R}^{>0}$ | Location hyperparameter for $\mu_{\mathbf{a}_1}$ in (4). Value defined in (10). |
| $\mu_{0,\mathbf{a}_2}$ | $\mathbb{R}^{>0}$ | Location hyperparameter for $\mu_{\mathbf{a}_2}$ in (7). Value defined in (13). |
| $\mu_{\mathbf{b}}$ | $\mathbb{R}$ | Location parameter for $\mathbf{b}_{\mathbf{c}_i,\mathbf{c}_j}$ in (28). Prior defined in (26). |
| $\mu_{0,\mathbf{b}}$ | $\mathbb{R}$ | Location hyperparameter for $\mu_{\mathbf{b}}$ in (26). Value defined in (31). |
| $\mu_{\gamma,p}$ | $\mathbb{R}$ | Location parameter for $\gamma_{\mathbf{c}_i,p}$ in (39). Prior defined in (37). |
| $\mu_{0,\gamma}$ | $\mathbb{R}$ | Location hyperparameter for $\mu_{\gamma,p}$ in (37). Value defined in (42). |
| $\pi_{\mathbf{c}}$ | $\Delta^{\infty}$ | Probabilities for $\mathbf{c}_i$ in (22). Prior defined in (21). |
| $\pi_{\mathbf{b}}$ | $(0,1)$ | Probability for $\mathbf{z}^{(\mathbf{b})}_{\mathbf{c}_i,\mathbf{c}_j}$ in (30). Prior distribution defined in (29) |
| $\pi_{\gamma,p}$ | $(0,1)$ | Probability for $\mathbf{z}^{(\gamma)}_{\mathbf{c}_i,p}$ in (41). Prior distribution defined in (40) |
| $\mathbf{Q}_{s,r}(k)$ | $\mathbb{R}^{>0}$ | Replicate qPCR measurement $r$ for subject $s$ at time $t_k$. Prior defined in (53). |
| $Q^{\text{data}}_{s,r}(k)$ | $\mathbb{R}^{>0}$ | Instantiation of the random variable for replicate qPCR measurement $r$ for subject $s$ at time $t_k$ with our data. |
| $r$ | $\mathbb{Z}^{+}$ | As a subscript this variable is the index for qPCR replicates. |
| $r_s(k)$ | $\mathbb{Z}$ | Read depth for sample at time index $k$ in subject $s$ |
| $\sigma^2_{\mathbf{a}_1}$ | $\mathbb{R}^{>0}$ | Squared scale parameter for $\mathbf{a}_{1,i}$ in (6). Prior defined in (5). |
| $\sigma^2_{\mathbf{a}_2}$ | $\mathbb{R}^{>0}$ | Squared scale parameter for $\mathbf{a}_{2,i}$ in (9). Prior defined in (8). |
| $\sigma^2_{0,\mathbf{a}_1}$ | $\mathbb{R}^{>0}$ | Squared scale hyperparameter for $\mu_{\mathbf{a}_1}$ in (4). Value defined in (11). |
| $\sigma^2_{0,\mathbf{a}_2}$ | $\mathbb{R}^{>0}$ | Squared scale hyperparameter for $\mu_{\mathbf{a}_2}$ in (7). Value defined in (14). |
| $\sigma^2_{\mathbf{b}}$ | $\mathbb{R}^{>0}$ | Squared scale parameter for $\mathbf{a}_{2,i}$ in (9). Prior defined in (27). |
| $\sigma^2_{0,\mathbf{b}}$ | $\mathbb{R}^{>0}$ | Squared scale hyperparameter for $\mu_{\mathbf{a}_2}$ in (7). Value defined in (32). |
| $\sigma^2_{\gamma,p}$ | $\mathbb{R}^{>0}$ | Squared scale parameter for $\mathbf{a}_{2,i}$ in (9). Prior defined in (38). |
| $\sigma^2_{0,\gamma}$ | $\mathbb{R}^{>0}$ | Squared scale hyperparameter for $\mu_{\gamma,p}$ in (37). Value defined in (43). |
| $\sigma^2_{\mathbf{Q}_s(k)}$ | $\mathbb{R}^{>0}$ | Squared scale hyperparameter for $\mathbf{Q}_{s,r}(k)$ in (53). Prior defined in (54). |
| $\sigma^2_w$ | $\mathbb{R}^{>0}$ | Process variance for the stochastic dynamics in (2)-(3). Prior defined in (47). |
| $\nu_{\mathbf{a}_1}$ | $\mathbb{R}^{>2}$ | DOF hyperparameter for $\sigma^2_{\mathbf{a}_1}$ in (5). Value defined in (16). |
| $\nu_{\mathbf{a}_2}$ | $\mathbb{R}^{>2}$ | DOF hyperparameter for $\sigma^2_{\mathbf{a}_2}$ in (8). Value defined in (17). |
| $\nu_{\mathbf{b}}$ | $\mathbb{R}^{>2}$ | DOF hyperparameter for $\sigma^2_{\mathbf{b}}$ in (27). Value defined in (33). |
| $\nu_{\gamma}$ | $\mathbb{R}^{>2}$ | DOF hyperparameter for $\sigma^2_{\gamma}$ in (38). Value defined in (44). |
| $\nu_w$ | $\mathbb{R}^{>2}$ | DOF hyperparameter for $\sigma^2_w$ in (47). Value defined in (48). |
| $\tau^2_{\mathbf{a}_1}$ | $\mathbb{R}^{>0}$ | Scale hyperparameter for $\sigma^2_{\mathbf{a}_1}$ in (5). Value defined in (18). |

Table 1 – *Continued on next page*

Table 1 – *Continued from previous page*

| variable | space | description |
|---|---|---|
| $\tau^2_{\mathbf{a}_2}$ | $\mathbb{R}^{>0}$ | Scale hyperparameter for $\sigma^2_{\mathbf{a}_2}$ in (8). Value defined in (19). |
| $\tau^2_{\mathbf{b}}$ | $\mathbb{R}^{>0}$ | Scale hyperparameter for $\sigma^2_{\mathbf{b}}$ in (27). Value defined in (34). |
| $\tau^2_{\gamma}$ | $\mathbb{R}^{>0}$ | Scale hyperparameter for $\sigma^2_{\gamma}$ in (5). Value defined in (45). |
| $\tau^2_{\sigma^2_w}$ | $\mathbb{R}^{>0}$ | Scale hyperparameter for $\sigma^2_w$ in (47). Value defined in (49). |
| $\theta_\alpha 1$ | $\mathbb{R}^{>0}$ | Shape hyperparameter for $\alpha$ in (20). Value defined in (24) |
| $\theta_\alpha 2$ | $\mathbb{R}^{>0}$ | Scale hyperparameter for $\alpha$ in (20). Value defined in (25) |
| $t_{s,k}$ | $\mathbb{R}$ | Time of sample index $k$ in subject $s$. |
| $\phi_{s,i}$ | $\mathbb{R}^{>0}$ | Location parameter for sequencing read counts $\mathbf{y}_{s,i}(k)$ in (50). Parameterization defined in (51) |
| $\mathbf{x}_{s,i}(k)$ | $\mathbb{R}^{>0}$ | Latent abundance of taxa $i$ at discrete time index $k$ in time-series $s$ for the dynamics in (2)-(3) |
| $\mathbf{y}_{s,i}(k)$ | $\mathbb{Z}$ | Sequencing reads for taxa $i$ at discrete time index $k$ in time-series $s$ |
| $y^{\text{data}}_{s,i}(k)$ | $\mathbb{Z}$ | Instantiation of the random variable for sequencing reads associated with taxa $i$ at discrete time index $k$ in time-series $s$ using the data collected in our experiments. |
| $\mathbf{z}^{(\mathbf{b})}_{\mathbf{c}_i,\mathbf{c}_j}$ | $\{0,1\}$ | Indicator variable for interaction from module $\mathbf{c}_j$ to $\mathbf{c}_i$ for the dynamics in (2)-(3). Prior defined in (30). |
| $\mathbf{z}^{(\gamma)}_{\mathbf{c}_i,p}$ | $\{0,1\}$ | Indicator variable for perturbation $p$ on module $\mathbf{c}_i$ for the dynamics in (2)-(3). Prior defined in (41). |

### 1 Description of Model

MDSINE2 is a Bayesian model for microbial dynamics based on generalized Lotka-Volterra (gLV) equations. Key attributes and novel components of the model that make it robust and allow it to scale to hundreds of taxa:

- It is fully Bayesian, explicitly modeling measurement error in amplicon sequencing and qPCR data, and propagating that uncertainty throughout the model.
- The model groups microbial taxa into *interaction modules*, or groups of taxa that share common responses to perturbations and interaction structure.
- The model includes indicator variables for both the perturbation affects and module-module interactions, which allows for structural learning and the computation of Bayes factors.
- Key posterior distributions (where feasible) utilize collapsed Gibbs sampling, decreasing mixing time in MCMC inference.

In the following subsections all model components are described in detail. To aid in the description the model, two versions of plate models are given in Figure 1 with all model parameters described in Table 1.

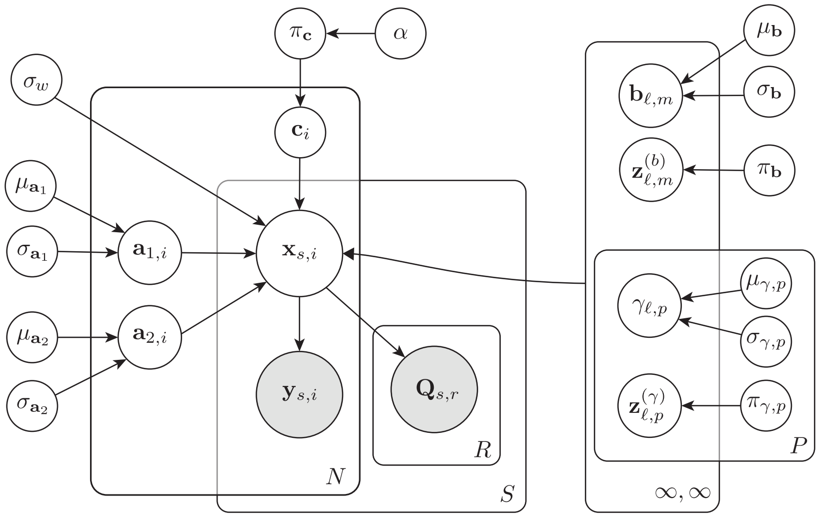


**Figure 1:** Graphical Model

#### 1.1 Model of dynamics

Let $\mathbf{x}_{s,i}(k) \in \mathbb{R}^{\geq 0}$ denote the latent concentration of taxa $i$ at time-point $t_{s,k}$ for subject $s$ (e.g., individual mouse or human subject). We model $\mathbf{x}_{s,i}(k)$ by discretizing the following continuous time $(t)$ stochastic differential equation (stochastic gLV dynamics):

$$\mathrm{d}\mathbf{x}_{s,i}(t) = \left[\mathbf{a}_{1,i}\left(1 + \sum_{p=1}^{P} \gamma_{\mathbf{c}_i,p}\mathbf{z}^{(\gamma)}_{\mathbf{c}_i,p} h_{s,p}(t)\right)\mathbf{x}_{s,i}(t) - \mathbf{a}_{2,i}(\mathbf{x}_{s,i}(t))^2 \right.$$
$$\left. + \sum_{j:\mathbf{c}_j \neq \mathbf{c}_i} \mathbf{b}_{\mathbf{c}_i,\mathbf{c}_j}\mathbf{z}^{(\mathbf{b})}_{\mathbf{c}_i,\mathbf{c}_j}\mathbf{x}_{s,i}(t)\mathbf{x}_{s,j}(t)\right]\mathrm{d}t + \mathbf{x}_{s,i}(t)\mathrm{d}w_{s,i}(t) \quad (1)$$

We discretize the above equation using a first-order approximation (see Appendix B for derivation) resulting in the following discrete time stochastic dynamics:

$$\log(\mathbf{x}_{s,i}(k+1)) \mid \mathbf{x}(k), \mathbf{a}, \mathbf{b}, \gamma, \mathbf{z}, \sigma_w^2 \sim \texttt{Normal}\big(\log(\mu_{s,i}(k+1)), \Delta_{s,k}\sigma_w^2\big) \quad (2)$$

where

$$\log(\mu_{s,i}(k+1)) \triangleq \log(\mathbf{x}_{s,i}(k)) + \Delta_{s,k}\left[\mathbf{a}_{1,i}\left(1+\sum_{p=1}^{P}\gamma_{\mathbf{c}_i,p}\mathbf{z}^{(\gamma)}_{\mathbf{c}_i,p}h_{s,p}(k)\right) - \mathbf{a}_{2,i}\mathbf{x}_{s,i}(k) + \sum_{j:\mathbf{c}_j\neq\mathbf{c}_i}\mathbf{b}_{\mathbf{c}_i,\mathbf{c}_j}\mathbf{z}^{(\mathbf{b})}_{\mathbf{c}_i,\mathbf{c}_j}\mathbf{x}_{s,j}(k)\right] \quad (3)$$

Starting from left to right in Equation (3) we have the step size in time $\Delta_{s,k} = t_{s,k+1} - t_{s,k}$ between two adjacent measurements $k$ and $k+1$ with corresponding times $t_{s,k}$ and $t_{s,k+1}$ in subject $s$. The next group of variables models the rate at which the taxa abundance increases over time with the unperturbed growth rate of taxa $i$ denoted as $\mathbf{a}_{1,i}$. During the course of the time series experiments, $P$ perturbations are introduced, which we model with a multiplicative effect on the growth rate. The strength of perturbation $p$ on taxa $i$ is denoted by $\gamma_{\mathbf{c}_i,p}$ along with a corresponding indicator variable $\mathbf{z}^{(\gamma)}_{\mathbf{c}_i,p}$, and $h_p(k)$, which is equal to one while the perturbation is active (zero otherwise). Note that the perturbation variables are indexed by $\mathbf{c}_i$ the module assignment for taxa $i$ (as described in detail below). The next variable appearing in the dynamics is the self interaction term $\mathbf{a}_{2,i}$. A classic logistic growth model would only have the growth rate term $\mathbf{a}_{1,i}$ (the slope on a log abundance plot over time) and the self interaction term $\mathbf{a}_{2,i}$, which together determine the steady state carrying capacity of the population being modeled (see Figure 2). The next set of parameters in the model capture the pairwise microbial interactions. The parameter $\mathbf{b}_{\mathbf{c}_i,\mathbf{c}_j}$ captures the strength of the interaction from taxa $j$ on taxa $i$ with $\mathbf{z}^{(\mathbf{b})}_{\mathbf{c}_i,\mathbf{c}_j}$ the corresponding indicator variable for that interaction. Note once again the interaction variables are indexed by the interaction module assignments $\mathbf{c}_i$ just as the perturbations were. With this model, the taxa in an interaction module share their response to perturbations and their module-module interaction strengths. Note that there are no interactions between taxa within the same module.

#### 1.2 Growth and self-limiting variables

We assume that each taxon has a positive growth rate $\mathbf{a}_{1,i} > 0$, as we seek to model colonizing taxa and not those that simply wash out. We also assume that each taxon has a positive carrying capacity $\frac{\mathbf{a}_{1,i}}{\mathbf{a}_{2,i}}$, and thus, $\mathbf{a}_{2,i} > 0$ as well (see Figure 2). We place hierarchical positively truncated Normal prior distributions on the growth rate and self interaction terms, respectively,

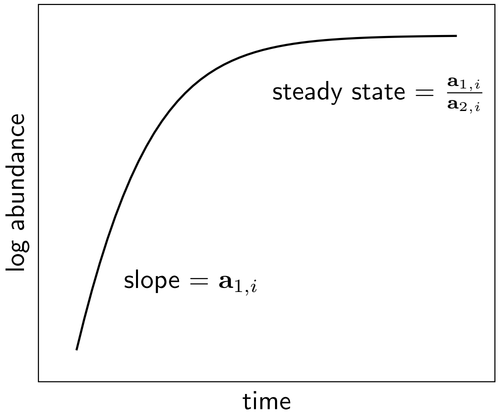


**Figure 2:** A simple logistic growth curve with growth rate $\mathbf{a}_{1,i}$ and limiting coefficient $\mathbf{a}_{2,i}$.

with their corresponding priors as follows:

$$\mu_{\mathbf{a}_1} \sim \texttt{TruncNormal}_{(0,\infty)}(\mu_{0,\mathbf{a}_1}, \sigma^2_{0,\mathbf{a}_1}) \tag{4}$$
$$\sigma^2_{\mathbf{a}_1} \sim \texttt{Scale-Inv-}\chi^2(\nu_{\mathbf{a}_1}, \tau^2_{\mathbf{a}_1}) \tag{5}$$
$$\mathbf{a}_{1,i} \mid \mu_{\mathbf{a}_1}, \sigma^2_{\mathbf{a}_1} \sim \texttt{TruncNormal}_{(0,\infty)}(\mu_{\mathbf{a}_1}, \sigma^2_{\mathbf{a}_1}) \tag{6}$$

and

$$\mu_{\mathbf{a}_2} \sim \texttt{TruncNormal}_{(0,\infty)}(\mu_{0,\mathbf{a}_2}, \sigma^2_{0,\mathbf{a}_2}) \tag{7}$$
$$\sigma^2_{\mathbf{a}_2} \sim \texttt{Scale-Inv-}\chi^2(\nu_{\mathbf{a}_2}, \tau^2_{\mathbf{a}_2}) \tag{8}$$
$$\mathbf{a}_{2,i} \mid \mu_{\mathbf{a}_2}, \sigma^2_{\mathbf{a}_2} \sim \texttt{TruncNormal}_{(0,\infty)}(\mu_{\mathbf{a}_2}, \sigma^2_{\mathbf{a}_2}). \tag{9}$$

The settings for the hyperparameters $\mu_{0,\mathbf{a}_1}$, $\sigma^2_{0,\mathbf{a}_1}$, $\nu_{\mathbf{a}_1}$, $\tau^2_{\mathbf{a}_1}$, $\mu_{0,\mathbf{a}_2}$, $\sigma^2_{0,\mathbf{a}_2}$, $\nu_{\mathbf{a}_2}$, $\tau^2_{\mathbf{a}_2}$ are given in the following subsection.

##### 1.2.1 Growth and self-limiting hyperparameters

The location and squared scale hyperparameters for the prior on $\mu_{\mathbf{a}_1}$ in Equation (4) are set as follows:

$$\mu_{0,\mathbf{a}_1} \triangleq 1 \tag{10}$$
$$\sigma^2_{0,\mathbf{a}_1} \triangleq 10^4 \cdot \texttt{median}(\hat{a}_1)^2 \tag{11}$$

where $\hat{a}_{1,i}$ is estimated from the deterministic logistic growth dynamics:

$$\log(x_{s,i}(k+1)) = \log(x_{s,i}(k)) + (a_{1,i} - a_{2,i}x_{s,i}(k))\Delta_{s,k}$$

which has the following least squares solution:

$$[\hat{a}_{1,i},\ \hat{a}_{2,i}] = \underset{[a_{1,i},\ a_{2,i}]}{\arg\min} \sum_{s,k} \|\log(x_{s,i}(k+1)) - \log(x_{s,i}(k)) + (a_{1,i} - a_{2,i}x_{s,i}(k))\Delta_{s,k}\|^2. \tag{12}$$

With $\mu_{0,\mathbf{a}_1}$ set to 1, this corresponds to a mean microbial doubling time of approximately 0.7 days and with the inflated variance in Equation (11) approximately 67% (assuming $\texttt{median}(\hat{a}_1)$ is approximately unity) of the support for $\mu_{\mathbf{a}_1}$ includes doubling times from 30 minutes to several days. This results in a relatively diffuse prior for $\mu_{\mathbf{a}_1}$ that reflects feasible doubling times for bacteria. The hyperparameters for $\mu_{\mathbf{a}_2}$ are set in a similar fashion with:

$$\mu_{0,\mathbf{a}_2} \triangleq \texttt{median}(\check{a}_2) \tag{13}$$
$$\sigma^2_{0,\mathbf{a}_2} \triangleq 10^4 \cdot \texttt{median}(\check{a}_2)^2 \tag{14}$$

where $\check{a}_{2,i}$ is inferred from the deterministic logistic growth dynamics:

$$\log(x_{s,i}(k+1)) = \log(x_{s,i}(k)) + (|\hat{a}_{1,i}| - a_{2,i}x_{s,i}(k))\Delta_{s,k}$$

which has the following least squares solution:

$$\check{a}_{2,i} = \underset{a_{2,i}}{\arg\min} \sum_{s,k} \|\log(x_{s,i}(k+1)) - \log(x_{s,i}(k)) + (|\hat{a}_{1,i}| - a_{2,i}x_{s,i}(k))\Delta_{s,k}\|^2. \tag{15}$$

This also results in a diffuse prior for $\mu_{\mathbf{a}_2}$, given that the squared scale is proportional to $10^4$ (scale is thus two orders of magnitude).

The hyperparameters for the prior on $\sigma^2_{\mathbf{a}_1}$ in Equation (5) and $\sigma^2_{\mathbf{a}_2}$ in Equation (8) are:

$$\nu_{\mathbf{a}_1} \triangleq 2.5 \tag{16}$$
$$\nu_{\mathbf{a}_2} \triangleq 2.5 \tag{17}$$
$$\tau^2_{\mathbf{a}_1} \triangleq 10^4 \cdot \texttt{median}(\hat{a}_1)^2 = \sigma^2_{0,\mathbf{a}_1} \tag{18}$$
$$\tau^2_{\mathbf{a}_2} \triangleq 10^4 \cdot \texttt{median}(\check{a}_2)^2 = \sigma^2_{0,\mathbf{a}_2}. \tag{19}$$

The degrees of freedom parameter for both is chosen as 2.5, because the scaled inverse $\chi^2$ distribution has an undefined first moment when the DOF is $\leq 2$.

#### 1.3 Dirichlet process prior on interaction modules

MDSINE2 automatically learns groups of taxa that share common interactions and perturbation effects, which we term *interaction modules*. The interaction module associated with each taxon $i$ is denoted $\mathbf{c}_i$. To model interaction modules, we use a Dirichlet Process (DP)-based prior probability distribution [16, 17].

The module assignment $\mathbf{c}_i$ for each taxon $i$ are generated can be viewed as being generated via the following stick-breaking process:

$$\alpha \sim \texttt{Gamma}(\theta_{\alpha 1}, \theta_{\alpha 2}) \tag{20}$$

$$\pi_{\mathbf{c}} \mid \alpha \sim \texttt{Stick}(\alpha) \tag{21}$$

$$\mathbf{c}_i \mid \pi_{\mathbf{c}} \sim \texttt{Multinomial}(\pi_{\mathbf{c}}) \tag{22}$$

where $\alpha$ is the concentration parameter, which implicitly controls the number of modules, $\pi_{\mathbf{c}}$ is an infinite dimensional vector where $\sum_{k=1}^{\infty} \pi_{\mathbf{c},k} = 1$ and $\pi_{\mathbf{c},k}$ is the probability that the $\mathbf{c}_i = k$ (taxa $i$ is in module $k$). An explicit construction for $\texttt{Stick}$ is given in Appendix C.

The number of modules in our model scales with how many taxa are present in the data and the concentration parameter $\alpha$. Specifically, the expected number of modules scales as: [1]:

$$\text{Expected number of modules} \approx \alpha \log\left(\frac{N_O + \alpha}{\alpha}\right) \tag{23}$$

where $N_O$ is the number of taxa.

##### 1.3.1 Concentration parameter hyperparameters

The shape and scale for the prior distribution on $\alpha$ in Equation (20) are defined as follows:

$$\theta_{\alpha 1} \triangleq 10^{-5} \tag{24}$$

$$\theta_{\alpha 2} \triangleq 10^{5}. \tag{25}$$

These parameters result in a diffuse prior on $\alpha$ with a mean of $1$ and a variance of $10^5$, and are consistent with our previous work on nonparametric models for microbial dynamics [4, 7]. With the diffuse prior for $\alpha$ in Equation (20), our model is not biased towards any particular number of modules (other than the log scaling with the data) and jointly learns the number of modules along with the rest of the model parameters.

##### 1.3.2 Connections between interaction modules and prior work

DPs have been widely used as priors in Bayesian clustering approaches, i.e., infinite mixture models [17]. Our model also has connections to Stochastic Block Models, Dependent Dirichlet Processes, and Topic Models [10, 13, 19, 15], which also can account for complex latent dependencies, similar in spirit to our model [8].

#### 1.4 Interaction variables

The variable $\mathbf{b}_{\mathbf{c}_i,\mathbf{c}_j}$ represents the interaction strength from interaction module $\mathbf{c}_j$ to interaction module $\mathbf{c}_i$, which can be a positive or negative value. We place a hierarchical Normal prior on the interaction strengths:

$$\mu_{\mathbf{b}} \sim \texttt{Normal}(\mu_{0,\mathbf{b}}, \sigma^2_{0,\mathbf{b}}) \tag{26}$$

$$\sigma^2_{\mathbf{b}} \sim \texttt{Scale-Inv-}\chi^2(\nu_{\mathbf{b}}, \tau^2_{\mathbf{b}}) \tag{27}$$

$$\mathbf{b}_{\mathbf{c}_i,\mathbf{c}_j} \mid \mu_{\mathbf{b}}, \sigma^2_{\mathbf{b}} \sim \texttt{Normal}(\mu_{\mathbf{b}}, \sigma^2_{\mathbf{b}}). \tag{28}$$

The variable $\mathbf{z}^{(\mathbf{b})}_{\mathbf{c}_i,\mathbf{c}_j}$ represents a binary indicator/selector, which determines whether or not there exists an interaction from interaction module $\mathbf{c}_j$ to interaction module $\mathbf{c}_i$. This variable effectively learns the underlying module interaction network topology. The hierarchical prior on $\mathbf{z}^{(\mathbf{b})}_{\mathbf{c}_i,\mathbf{c}_j}$ is as follows

$$\pi_{\mathbf{b}} \sim \texttt{Beta}(b_{1,\mathbf{b}}, b_{2,\mathbf{b}}) \tag{29}$$
$$\mathbf{z}^{(\mathbf{b})}_{\mathbf{c}_i,\mathbf{c}_j} \mid \pi_{\mathbf{b}} \sim \texttt{Bernoulli}(\pi_{\mathbf{b}}). \tag{30}$$

As previously stated, with this construction interactions only exist between modules. Taxa within the same module do not have pairwise ecological interactions. The hyperparameters $\mu_{0,\mathbf{b}}$, $\sigma^2_{0,\mathbf{b}}$, $\nu_{\mathbf{b}}$, $\tau^2_{\mathbf{b}}$, $b_{1,\mathbf{b}}$, and $b_{2,\mathbf{b}}$ are defined in the following subsection.

##### 1.4.1 Interaction hyperparameters

The hyperparameters for $\mu_{\mathbf{b}}$ in Equation (26) are:

$$\mu_{0,\mathbf{b}} \triangleq 0 \tag{31}$$
$$\sigma^2_{0,\mathbf{b}} \triangleq 10^4 \cdot \texttt{median}(\breve{a}_2)^2 = \sigma^2_{0,\mathbf{a}_2} \tag{32}$$

The location parameter $\mu_{0,\mathbf{b}}$ is chose to be 0 so that we are not biasing $\mu_{\mathbf{b}}$ to be positive or negative. The squared scale parameter $\sigma^2_{0,\mathbf{b}}$ is set to be identical to the squared scale parameter $\sigma^2_{0,\mathbf{a}_2}$. We set the hyperparameters for $\sigma^2_{\mathbf{b}}$ in Equation (27) to be the same as their counterpart hyperparameters for $\sigma^2_{\mathbf{a}_2}$ in Equation (8) which were originally defined in Equations (17) and (19) and are as follows

$$\nu_{\mathbf{b}} \triangleq 2.5 \tag{33}$$
$$\tau^2_{\mathbf{b}} \triangleq 10^4 \cdot \texttt{median}(\breve{a}_2)^2 = \tau^2_{\mathbf{a}_2}. \tag{34}$$

The hyperparameters for the prior on $\pi_{\mathbf{b}}$ are selected to bias the model towards a sparse topology, analogous to our previous MDSINE model [3], and are defined as follows

$$b_{1,\mathbf{b}} \triangleq 0.5 \tag{35}$$
$$b_{2,\mathbf{b}} \triangleq N_b\,(N_b - 1). \tag{36}$$

Here, $N_b$ is the expected number of clusters for a Dirichlet Process with concentration $\alpha$ and number of taxa $N_O$ as defined in Equation (23). Note that $N_b(N_b - 1)$ is the total number of ordered pairs (directed edges) of a network with $N_b$ modules. Thus, the prior distribution yields an expected number of interactions of $0.5$ (i.e., less than one expected interaction [3]), constituting a "strong" prior probability of no interactions in the interaction module network.

#### 1.5 Perturbation effects

The variable $\gamma_{\mathbf{c}_i,p}$ captures the multiplicative effect of the $p$-th perturbation on the growth rate of taxa in module $\mathbf{c}_i$ through the expression $(1+\gamma_{\mathbf{c}_i,p})$ in Equation (1). We place hierarchical, normal prior probability distributions on $\gamma_{\mathbf{c}_i,p}$ (analogous to the priors we used on the interactions)

$$\mu_{\gamma,p} \sim \texttt{Normal}(\mu_{0,\gamma}, \sigma^2_{0,\gamma}) \tag{37}$$
$$\sigma^2_{\gamma,p} \sim \texttt{Scale-Inv-}\chi^2(\nu_\gamma, \tau^2_\gamma) \tag{38}$$
$$\gamma_{\mathbf{c}_i,p} \mid \mu_{\gamma,p}, \sigma^2_{\gamma,p} \sim \texttt{Normal}(\mu_{\gamma,p}, \sigma^2_{\gamma,p}) \tag{39}$$

The variable $\mathbf{z}^{(\gamma)}_{\mathbf{c}_i,p}$ is a binary indicator/selector (analogous to $\mathbf{z}^{(\mathbf{b})}_{\mathbf{c}_i,\mathbf{c}_j}$, the interaction selector variable), which determines whether or not there exists an effect of perturbation $p$ on module $\mathbf{c}_i$. The hierarchical prior on $\mathbf{z}^{(\gamma)}_{\mathbf{c}_i,p}$ is as follows:

$$\pi_{\gamma,p} \sim \texttt{Beta}(b_{1,\gamma}, b_{2,\gamma}) \tag{40}$$
$$\mathbf{z}^{(\gamma)}_{\mathbf{c}_i,p} \mid \pi_{\gamma,p} \sim \texttt{Bernoulli}(\pi_{\gamma,p}). \tag{41}$$

##### 1.5.1 Perturbation hyperparameters

The hyperparameters for the prior on $\mu_{\gamma,p}$ in Equation (37) is centered around 0 and is set to be diffuse as for our previous priors:

$$\mu_{0,\gamma} \triangleq 0 \tag{42}$$

$$\sigma^2_{0,\gamma} \triangleq 10^4 \tag{43}$$

Similarly, the prior for $\sigma^2_{\gamma,p}$ in Equation (38) is diffuse with the scale chosen so that under the prior $\mathbb{E}\left[\sigma^2_{\gamma,p}\right] = 10^4$,

$$\nu_\gamma \triangleq 2.5 \tag{44}$$

$$\tau^2_\gamma \triangleq 10^4 \cdot \frac{\nu_\gamma - 2}{\nu_\gamma}. \tag{45}$$

The hyperparameters in (40) for $\pi_{\gamma,p}$ are selected so that the prior is diffuse and unbiased with regards to whether a perturbation impacts a module or not with:

$$b_{1,\gamma} \triangleq 0.5 \quad \text{and} \quad b_{2,\gamma} \triangleq 0.5 \tag{46}$$

which results in $\mathbb{E}[\pi_{\gamma,p}] = 0.5$ under the prior.

#### 1.6 Process variance

The process variance $\sigma^2_w$ has the following prior:

$$\sigma^2_w \sim \texttt{Scale-Inv-}\chi^2(\nu_w, \tau^2_{\sigma^2_w}). \tag{47}$$

##### 1.6.1 Process variance hyperparameters

The hyperparameters $\nu_w$ and $\tau^2_{\sigma^2_w}$ are defined as follows

$$\nu_w \triangleq 2.5 \tag{48}$$

$$\tau^2_{\sigma^2_w} \triangleq 0.2^2 \cdot \frac{\nu_w - 2}{\nu_w}, \tag{49}$$

where the prior is diffuse with a low degree of freedom and the scale parameter is chosen so that under the prior $\mathbb{E}[\sigma^2_w] = .2^2$ which corresponds to $20\%$ variation for the multiplicative process variance in our model.

#### 1.7 Measurement error model

The observed data consist of sequencing counts $\mathbf{y}_{s,i}(k)$ and qPCR measurements $\mathbf{Q}_{s,r}(k)$, for subject $s$, taxon $i$, time-point $t_k$ and qPCR replicate $r$. We assume that observed data are generated by the underlying latent taxa concentrations $\mathbf{x}_s(k)$. As in our previous MDSINE model [3], and other sequencing data error models [14, 12], we model sequencing counts $\mathbf{y}_{s,i}(k)$ using a negative binomial distribution parameterized by its mean, $\phi_{si}$, and dispersion parameter $\epsilon_{si}(\cdot)$:

$$\mathbf{y}_{s,i}(k) \mid \mathbf{x}_s(k) \sim \texttt{NegBin}\Big(\phi_{s,i}(\mathbf{x}_s(k), r_s(k)), \epsilon_{s,i}(\mathbf{x}_s(k), d_0, d_1)\Big). \tag{50}$$

The mean $\phi_{s,i}$ is proportional to the relative abundance of the taxon:

$$\phi_{s,i}(\mathbf{x}_s(k), r_s(k)) \triangleq r_s(k)\frac{\mathbf{x}_{s,i}(k)}{\sum_i \mathbf{x}_{s,i}(k)} \tag{51}$$

where $r_s(k)$ is the total number of reads at time $t_k$ for subject $s$. The dispersion parameter $\epsilon_{s,i}$ is defined as follows:

$$\epsilon_{s,i}(\mathbf{x}_s(k), d_0, d_1) \triangleq \frac{d_0}{\mathbf{x}_{s,i}(k)/\sum_i \mathbf{x}_{s,i}(k)} + d_1 \quad (52)$$

where $d_0$ and $d_1$ are parameters pre-trained on replicates before inference is performed on the model (see Appendix E).

We place a log-Normal prior distribution on the qPCR measurements, $\mathbf{Q}_{s,r}(k)$, with the mean of the distribution parametrized by the total latent concentration of taxa for the given sample being modeled:

$$\log(\mathbf{Q}_{s,r}(k)) \mid \mathbf{x}_s(k) \sim \texttt{Normal}\Big(\log\Big(\sum_i \mathbf{x}_{s,i}(k)\Big), \sigma^2_{\mathbf{Q}_s(k)}\Big) \quad (53)$$

where

$$\sigma^2_{\mathbf{Q}_s(k)} \triangleq \widetilde{\mathbb{V}}^3_{r=1}\left[\log Q^{\text{data}}_{s,r}(k)\right] \quad (54)$$

is the empirical variance for the log of the qPCR replicate measurement values, $Q^{\text{data}}_{s,r}(k)$, for subject $s$ at time-point $k$ over the $r$ replicates.

### 2 Model inference

We employ Markov Chain Monte-Carlo sampling with Gibbs and collapsed Gibbs sampling when possible and Metropolis-Hastings (MH) when direct sampling from the posterior is not possible. The order in which the parameters of the model are inferred is as follows:

1. Sample cluster interaction indicators $\mathbf{z}^{(\mathbf{b})}$, then their probabilities $\pi_{\mathbf{b}}$ (Section 2.4).
2. Sample perturbation indicators $\mathbf{z}^{(\gamma)}_p$, then their probabilities $\pi_{\gamma,p}$ (Section 2.4).
3. Sample interaction magnitudes $\mathbf{b}_{\mathbf{c}_i,\mathbf{c}_j}$ and perturbation magnitudes $\gamma_{\mathbf{c}_i,p}$ jointly (Section 2.1).
4. Sample growth rates $\mathbf{a}_1$ and self-interactions $\mathbf{a}_2$ (Section 2.2).
5. Sample prior means $\mu_{\mathbf{b}}$ and $\mu_{\gamma,p}$ (Section 2.1).
6. Sample prior means $\mu_{\mathbf{a}_1}$ and $\mu_{\mathbf{a}_2}$ (Section 2.2).
7. Sample prior variances $\sigma^2_{\mathbf{a}_1}$ and $\sigma^2_{\mathbf{a}_2}$ (Section 2.2).
8. Sample prior variances $\sigma^2_{\mathbf{b}}$ and $\sigma^2_{\gamma}$ (Section 2.1).
9. Sample process variance $\sigma^2_w$ (Section 2.3).
10. Sample latent trajectories $\mathbf{x}_{s,i}(k)$ (Section 2.5).
11. Sample cluster assignments $\mathbf{c}_i$ (Section 2.6).
12. Sample concentration parameter $\alpha$ (Section 2.6).

#### Notation

We let $\Omega$ denote the set of all model parameters and the $\setminus$ symbol is used to subtract from that set. For instance $\Omega \setminus \mathbf{b}$ is all model parameters except for the interaction strengths. To denote a partial vector of variables, we use the $_-$ symbol. For example $\mathbf{c}_{\text{-}i}$ indicates all cluster assignments except for that for taxon $i$. When referring to specific Gibbs samples we will use a superscript $[\cdot]$ to denote which one (e.g. $\mathbf{b}^{[g]}$ for the $g$-th sample of all the interaction strengths between modules) with $(\cdot)^{[0]}$ denoting initial conditions.

#### 2.1 Sampling perturbation and interaction strengths and their priors

The joint conditional distribution for $\mathbf{b}$ and $\gamma$ is:

$$p(\mathbf{b},\gamma \mid \Omega \setminus (\mathbf{b},\gamma)) = \prod_{s,i,k} \texttt{Normal}\big(\log(\mathbf{x}_{s,i}(k)) \mid \log(\mu_{s,i}(k)), \Delta_{s,k}\sigma_w^2\big)$$
$$\times \prod_{\ell,m} \texttt{Normal}\big(\mathbf{b}_{\ell,m} \mid \mu_{\mathbf{b}}, \sigma_{\mathbf{b}}^2\big)$$
$$\times \prod_{\ell,p} \texttt{Normal}\big(\gamma_{\ell,p} \mid \mu_{\gamma,p}, \sigma_{\gamma,p}^2\big)$$

The posterior can be written in closed form and sampled from directly (see Appendix D).

We initialize all interactions and perturbations to be zero, $\mathbf{b}^{[0]} = 0$ and $\gamma^{[0]} = 0$.

The parameters $\mu_{\mathbf{b}}$ and $\sigma_{\mathbf{b}}^2$ have conjugate priors with posteriors:

$$p(\mu_{\mathbf{b}} \mid \mathbf{b}) = \prod_{\ell,m} \texttt{Normal}(\mathbf{b}_{\ell,m} \mid \mu_{\mathbf{b}}, \sigma_{\mathbf{b}}^2) \cdot \texttt{Normal}(\mu_{\mathbf{b}}; \mu_{0,\mathbf{b}}, \sigma_{0,\mathbf{b}}^2)$$

and

$$p(\sigma_{\mathbf{b}}^2 \mid \mathbf{b}) = \prod_{\ell,m} \texttt{Normal}(\mathbf{b}_{\ell,m} \mid \mu_{\mathbf{b}}, \sigma_{\mathbf{b}}^2) \cdot \texttt{Scale-Inv-}\chi^2(\sigma_{\mathbf{b}}^2; \nu_{\mathbf{b}}, \tau_{\mathbf{b}}^2)$$

from which posteriors can be written in closed form and sampled from directly [6]. The parameters $\mu_{\gamma,p}$ and $\sigma_{\gamma,p}^2$ have conjugate priors of identical structure with corresponding posteriors of identical form that can be sampled from directly as well.

The location parameters are initialized as $\mu_{\mathbf{b}}^{[0]} = 0$ and $\mu_{\gamma,p}^{[0]} = 0$ with the squared scale parameters initialized as $\left(\sigma_{\mathbf{b}}^2\right)^{[0]} = \sigma_{0,\mathbf{b}}^2$ with $\sigma_{0,\mathbf{b}}^2$ defined in Equation (32) and $\left(\sigma_{\gamma,p}^2\right)^{[0]} = \sigma_{0,\gamma}^2$ with $\sigma_{0,\gamma}^2$ defined in Equation (43).

#### 2.2 Sampling growth rates, self-interaction strengths and parameters of their priors

The growth $\mathbf{a}_1$ and self-interactions $\mathbf{a}_2$ both have truncated Normal priors. We sample these parameters consecutively (but in a random order at each Gibbs step):

1. Sample $\mathbf{a}_1 \mid \Omega \setminus \mathbf{a}_1$
2. Sample $\mathbf{a}_2 \mid \Omega \setminus \mathbf{a}_2$

Under this conditioning, the posterior of $\mathbf{a}_{1,i}$ is a scalar truncated Normal distribution:

$$p(\mathbf{a}_{1,i} \mid \Omega \setminus \mathbf{a}_{1,i}) \propto \prod_{s,k} \texttt{Normal}\big(\log(\mathbf{x}_{s,i}(k)) \mid \log(\mu_{s,i}(k)), \Delta_{s,k}\sigma_w^2\big)$$
$$\times \texttt{TruncNormal}_{(0,\infty)}(\mathbf{a}_{1,i} \mid \mu_{\mathbf{a}_1}, \sigma_{\mathbf{a}_1}^2)$$

which can be sampled from directly; the posterior for $\mathbf{a}_{2,i}$ has an identical structure.

The growth and self interaction parameters are initialized using the regression results from Section 1.2.1, where $\mathbf{a}_1^{[0]} = |\hat{a}_1|$ in (12) and $\mathbf{a}_2^{[0]} = |\check{a}_2|$ in (15).

The location parameter $\mu_{\mathbf{a}_1}$ posterior is:

$$p(\mu_{\mathbf{a}_1} \mid \mathbf{a}_1) \propto \prod_{i} \texttt{TruncNormal}_{(0,\infty)}\big(\mathbf{a}_{1,i}; \mu_{\mathbf{a}_1}, \sigma_{\mathbf{a}_1}^2\big) \cdot \texttt{TruncNormal}_{(0,\infty)}\big(\mu_{\mathbf{a}_1}; \mu_{0,\mathbf{a}_1}, \sigma_{0,\mathbf{a}_1}^2\big)$$

which cannot be sampled from directly in closed form. Thus we use the MH algorithm [6] for sampling $\mu_{\mathbf{a}_1}$ with the following proposal distribution:

$$\mu_{\mathbf{a}_1}^{[*]} \sim \texttt{TruncNormal}_{(0,\infty)}(\mu_{\mathbf{a}_1}^{[g-1]}, \sigma_{1,\text{prop}}^2).$$

The proposal variance is initialized to $\sigma^2_{1,\text{prop}} = \mu^2_{0,\mathbf{a}_1}$ and is tuned during the first half of burn-in to adjust the acceptance rate to 0.44 (an empirically optimized rate for a MH step on a scalar variable [6]). For the details on how this tuning is done, see Appendix F. The location parameter $\mu_{\mathbf{a}_2}$ is updated analogously, using the MH algorithm with a similar proposal distribution that is centered on the previous Gibbs sample.

The location parameters are initialized as $\mu^{[0]}_{\mathbf{a}_1} = \mu_{0,\mathbf{a}_1}$ and $\mu^{[0]}_{\mathbf{a}_2} = \mu_{0,\mathbf{a}_2}$, where $\mu_{0,\mathbf{a}_1}$ and $\mu_{0,\mathbf{a}_2}$ are defined in Equations (10) and (13).

The squared scale parameters $\sigma^2_{\mathbf{a}_1}$ and $\sigma^2_{\mathbf{a}_2}$ both have a $\texttt{Scale-Inv-}\chi^2$ prior. However, these priors are not conjugate to the truncated normal distributions of $\mathbf{a}_1$ and $\mathbf{a}_2$. We use an MH update for these parameters as well. With $\sigma^2_{\mathbf{a}_1}$ as an example (the equations for $\sigma^2_{\mathbf{a}_2}$ are exactly analogous), the posterior for $\sigma^2_{\mathbf{a}_1}$ is:

$$p(\sigma^2_{\mathbf{a}_1} \mid \mathbf{a}_1) = \texttt{Scale-Inv-}\chi^2(\sigma^2_{\mathbf{a}_1}; \nu_{\mathbf{a}_1}, \tau^2_{\mathbf{a}_1}) \prod_i \texttt{TruncNormal}_{(0,\infty)}\big(\mathbf{a}_{1,i}; \mu_{\mathbf{a}_\ell}, \sigma^2_{\mathbf{a}_\ell}\big).$$

The proposal distribution for $\sigma^2_{\mathbf{a}_1}$ is a scaled inverse $\chi^2$ distribution (approximately) centered at the previous value. More precisely,

$$(\sigma^2_{\mathbf{a}_1})^{[*]} \sim \texttt{Scale-Inv-}\chi^2\Big(\nu_{1,\text{prop}}, (\sigma^2_{\mathbf{a}_1})^{[g-1]}\Big).$$

The parameter $\nu_{1,\text{prop}}$ is initialized to $15$ (which results in an initial proposal variance of $0.24 \cdot (\sigma^2_{\mathbf{a}_1})^{[g-1]}$), and is tuned during the first half of burn-in to adjust the acceptance rate to 0.44 (see Appendix F for details).

The squared scale parameters are initialized as $\left(\sigma^2_{\mathbf{a}_1}\right)^{[0]} = \sigma^2_{0,\mathbf{a}_1}$ and $\left(\sigma^2_{\mathbf{a}_2}\right)^{[0]} = \sigma^2_{0,\mathbf{a}_2}$, where $\sigma^2_{0,\mathbf{a}_1}$ and $\sigma^2_{0,\mathbf{a}_2}$ are defined in Equations (11) and (14).

#### 2.3 Sampling process variance

The process variance $\sigma^2_w$'s prior in (47) is conjugate to the distribution for the latent trajectory dynamics in (2). Its posterior can be sampled from directly with Gibbs updates being drawn from:

$$\sigma^2_w \mid \Omega_{\neg\sigma^2_w} \sim \texttt{Scale-Inv-}\chi^2\left(\nu_w + N_w, \frac{\nu_w \tau^2_{\sigma^2_w} + N_w S^2_w}{\nu_w + N_w}\right)$$

where $N_w$ is the a product of the number of subjects, the number of OTUs and one less the number of time points for the samples. Here, $S^2_w = \widetilde{\mathbb{V}}_{i,s,k}[\log \mathbf{x}_{s,i}(k) - \log \mathbf{x}_{s,i}(k-1)]$, which is the empirical variance of the differences in log-abundances. The process variance is initialized to the mean of its prior, Equation (47), as $\left(\sigma^2_w\right)^{[0]} = 0.2^2$.

#### 2.4 Sampling indicators variables and their priors

We update the interaction indicators $\mathbf{z}^{(\mathbf{b})}_{\ell,m}$ between modules $m$ and $\ell$ in a random order at each Gibbs step. For efficiency, we perform collapsed Gibbs sampling with marginalization over the interaction variables:

$$\begin{aligned} P\Big(\mathbf{z}^{(\mathbf{b})}_{\ell,m} = u \mid \Omega \setminus (\mathbf{b}, \mathbf{z}^{(\mathbf{b})}_{\ell,m})\Big) &= \int p\Big(\mathbf{z}^{(\mathbf{b})}_{\ell,m} = u, \mathbf{b} \mid \Omega \setminus (\mathbf{b}, \mathbf{z}^{(\mathbf{b})}_{\ell,m})\Big) \mathrm{d}\mathbf{b} \\ &= \texttt{Bernoulli}(u; \pi_{\mathbf{b}}) \int \prod_{s,i,k} \texttt{Normal}\big(\log \mathbf{x}_{s,i}(k); \log \mu_{s,i}(k), \Delta_{s,k}\sigma^2_w\big) \\ &\qquad \times \prod_{\bar{\ell},\bar{m}\,:\,\bar{m}\neq\bar{\ell}} \texttt{Normal}(\mathbf{b}_{\bar{\ell},\bar{m}}; \mu_{\mathbf{b}}, \sigma^2_{\mathbf{b}}) \mathrm{d}\mathbf{b} \end{aligned} \tag{55}$$

The prior probability for an interaction $\pi_{\mathbf{b}}$ has a conjugate beta prior and its posterior

$$\pi_{\mathbf{b}} \mid \mathbf{z}^{(\mathbf{b})} \sim \texttt{Beta}\left(b_{1,\mathbf{b}} + \sum_{\ell,m\,:\,m\neq\ell} \mathbf{z}^{(\mathbf{b})}_{\ell,m}, \; b_{2,\mathbf{b}} + \sum_{\ell,m\,:\,m\neq\ell} \Big(1 - \mathbf{z}^{(\mathbf{b})}_{\ell,m}\Big)\right) \tag{56}$$

can be sampled from directly. The corresponding variables $\pi_{\gamma,p}$ and $\mathbf{z}^{(\gamma)}_{\mathbf{c}_i,p}$ have posteriors of identical form that can be sampled from directly.

The indicators for both the interactions and the perturbations are initialized to zero,

$$\left(\mathbf{z}^{(\mathbf{b})}_{\ell,m}\right)^{[0]} = 0 \quad \text{and} \quad \left(\mathbf{z}^{(\gamma)}_{\mathbf{c}_i,p}\right)^{[0]} = 0.$$

The prior probabilities are initialized to the mean of their corresponding priors in Equations (29) and (40) as $\pi^{[0]}_{\mathbf{b}} = \frac{0.5}{0.5+N_b(N_b-1)}$ and $\pi^{[0]}_{\gamma,p} = 0.5$.

#### 2.5 Sampling latent trajectories

The latent state $\mathbf{x}_{s,i}(k)$ has a posterior:

$$\begin{aligned} p(\mathbf{x}_{s,i}(k) \mid \Omega \setminus \mathbf{x}_{s,i}(k)) = {} & \mathtt{NegBin}(\mathbf{y}_{s,i}(k); \phi_{s,i}, \epsilon_{s,i}) \\ & \times \prod_r \mathtt{Normal}\left(\log(\mathbf{Q}_{r,s}(k)); \log\left(\sum_i \mathbf{x}_{s,i}(k)\right), \sigma^2_{\mathbf{Q}_s(k)}\right) \\ & \times \mathtt{Normal}\big(\log(\mathbf{x}_{s,i}(k)); \log(\mu_{s,i}(k)), \Delta_{s,k}\sigma^2_w\big) \\ & \times \mathtt{Normal}\big(\log(\mathbf{x}_{s,i}(k+1)); \log(\mu_{s,i}(k+1)), \Delta_{s,k+1}\sigma^2_w\big) \end{aligned}$$

which is nonlinear in the latent state and does not have a closed form that can be sampled from. We use an MH step for $\mathbf{x}_{s,i}(k)$ with a proposal distribution centered at the previous MCMC state:

$$\log(\mathbf{x}^{[*]}_{s,i}(k)) \sim \mathtt{Normal}\left(\log\left(\mathbf{x}^{[g-1]}_{s,i}(k)\right), \sigma^2_{\mathbf{x},s,\text{prop}}\right)$$

where the proposal variance $\sigma^2_{\mathbf{x},s,\text{prop}}$ is optimized for each subject $s$ independently. The proposal variance is tuned to achieve the acceptance rate of 0.44 (see Appendix F).

We initialize the latent trajectory for each subject $s$ and taxa $i$ using a random truncated normal centered around a LOESS interpolation from the data. More specifically, for each subject $s$, we take

$$\mathbf{x}^{[0]}_{s,i}(k) \sim \mathtt{TruncNormal}_{(0,\infty)}\left(\hat{x}^{\text{loess}}_{s,i}(k), \sigma^2_{\hat{x},\text{loess}}\right)$$

where $\hat{x}^{\text{loess}}_{s,i}(k)$ is the degree-1 LOESS interpolation over each time series $s$, using as input bacterial abundances obtained by multiplying the relative abundance of each OTU $i$ from the sequencing reads for subject $s$ at time index $k$, $y_{s,i}(k)$, by the geometric mean of the qPCR measurements vector, $Q_s(k)$, that is:

$$\begin{aligned} \hat{x}^{\text{loess}}_{s} &\triangleq \mathtt{LOESS}(\hat{x}^{\text{data}}_{s}) \\ \hat{x}^{\text{data}}_{s,i}(k) &\triangleq \frac{y_{s,i}(k)}{\sum_i y_{s,i}(k)} \cdot \mathtt{geometricmean}(Q_s(k))). \end{aligned}$$

The squared scale parameter $\sigma^2_{\hat{x},\text{init}}$ is defined as

$$\sigma^2_{\hat{x},\text{init}} \triangleq 10^{-4}\left(\hat{x}^{\text{loess}}_{s,i}(k)\right)^2 + 10^{-4}$$

so that the sampled values for the initialization are close to the LOESS interpolation.

#### 2.6 Sampling module assignments and DP priors

We sample the module assignement for each taxon in a random order at each Gibbs step. For efficiency, we perform collapsed Gibbs sampling, integrating out the interaction and perturbation strength parameters. Let $\mathbf{c}_{\text{-}i}$ denote the current clustering assignments of all taxa except for taxon $i$. The likelihood of the latent trajectory for taxon $i$, given that the taxon has been assigned to module $m$, is:

$$P(\mathbf{x} \mid \mathbf{c}_i = m, \mathbf{z}, \mathbf{c}_{\text{-}i}, \sigma_w^2) = \int_{\gamma, \mathbf{b}} \prod_{s,i,k} \texttt{Normal}\big(\log(\mathbf{x}_{s,i}(k)); \log(\mu_{s,i}(k)), \Delta_{s,k}\sigma_w^2\big)$$

Using Algorithm 8 in [16], the posterior probability for taxon $i$ being assigned to module $m$ is:

$$P(\mathbf{c}_i = m \mid \mathbf{x}, \mathbf{z}, \mathbf{c}_{\text{-}i}, \sigma_w^2) \propto \begin{cases} n_{\neg i,m} \cdot P(\mathbf{x} \mid \mathbf{c}_i = m, \mathbf{z}, \mathbf{c}_{\text{-}i}, \sigma_w^2) & \text{if } n_{\text{-}i,m} > 0 \\ \alpha \cdot P(\mathbf{x} \mid \mathbf{c}_i = m, \mathbf{z}, \mathbf{c}_{\text{-}i}, \sigma_w^2) & \text{if } n_{\text{-}i,m} = 0 \end{cases}$$

where $n_{\text{-}i,m}$ are the number of taxa in module $m$ excluding taxon $i$ (equal to the size of module $m$ minus 1 if it previously contained $i$, and zero if it's a new module).

The module assignments are initialized by performing agglomerative clustering (stopped at 30 clusters) using Spearman rank correlation as the linkage measure on the taxa trajectories estimated from data. We sample the concentration parameter $\alpha$ using an auxiliary variable method as reported in [5, §6 Eqs. (13) and (14)]

#### 2.7 Mixing

We assess MCMC mixing using the metric $\hat{R}$ defined in [2], Equation 1.1:

$$\widehat{R} = \sqrt{\left(\frac{N-1}{N} + \frac{M+1}{MN}\frac{B}{W}\right)\frac{d+3}{d+1}}$$

where $M$ is the number of independent markov chains, $N$ is the length of each chain, $d$ is the method-of-moments estimator for the degrees of freedom as in [2], $B$ is the between-sequence variance (sample variance of the mean value of each chain), and $W$ is the within-sequence variance (the mean of the sample variance of each chain).

### 3 Extended discussion

#### 3.1 Stability

One of the necessary conditions for the global stability of the dynamics in Equation (1) is that the interaction matrix $A$, whose elements are defined as $[A]_{i,j} \triangleq \mathbf{b}_{\mathbf{c}_i,\mathbf{c}_j}\mathbf{z}^{(\mathbf{b})}_{\mathbf{c}_i,\mathbf{c}_j}$, is Diagonally Stable (D-stable), see [10, Supplementary Text §4.2 Theorem 5] and the original proof by Goh in [9]. A matrix is said to be diagonally stable if there exist a positive definite and diagonal matrix $D$ such that $A^{\mathsf{T}}D + DA < 0$. Necessary and sufficient conditions for a matrix being D-stable are non-trivial to check. However, one condition that is only necessary for D-stability is that all the eigenvalues of $A$ have real parts less than zero. This is equivalent to the relaxed condition satisfied by the Lyapunov equation, that there exists a positive definite $P$ (not necessarily diagonal) such that $A^{\mathsf{T}}P + PA < 0$. We chose to analyze this necessary condition in the stability analysis of our inferred dynamics because it is computationally efficient to check and necessary for D-stability.

### A Probability density functions

To resolve any ambiguity in the parameterization of standard probability distributions, we list their density functions here.

- The $\texttt{Scale-Inv-}\chi^2$distribution is parametrized with degrees of freedom $\nu$ and scale $\tau^2$, and has density

$$\texttt{Scale-Inv-}\chi^2(x;\nu,\tau^2) = \frac{(\tau^2\nu/2)^{-\nu/2}}{\Gamma(\nu/2)} x^{(-\nu/2)-1} \exp\Big(-\frac{\nu\tau^2}{2x}\Big).$$

- The gamma distribution is parameterized with shape $k$ and scale $\theta$, and has density

$$\texttt{Gamma}(x;k,\theta) = \frac{1}{\Gamma(k)\theta^k} x^{k-1} \exp\Big(-\frac{x}{\theta}\Big).$$

- The negative binomial distribution is parametrized with mean $\phi$ and dispersion $\epsilon$. Its density is

$$\texttt{NegBin}(y;\phi,\epsilon) = \frac{\Gamma(r+y)}{y!\Gamma(r)} \Big(\frac{\phi}{r+\phi}\Big)^y \Big(\frac{r}{r+\phi}\Big)^r$$

  where $r = 1/\epsilon$.

- The uniform distribution (over an interval) is parametrized with support $[a,b]$ as

$$\texttt{Uniform}(x;a,b) = \begin{cases} \frac{1}{b-a} & \text{for } x \in [a,b] \\ 0 & \text{otherwise.} \end{cases}$$

- The beta distribution is parametrized by two scale parameters $\alpha, \beta > 0$

$$\texttt{Beta}(x;\alpha,\beta) = x^{\alpha-1}(1-x)^{\beta-1} \frac{\Gamma(\alpha+\beta)}{\Gamma(\alpha)\Gamma(\beta)}.$$

- The truncated Normal distribution is parametrized with location $\mu$, scale squared $\sigma^2$, and support $(v_1, v_2)$ as

$$\texttt{TruncNormal}_{(v_1,v_2)}(x;\mu,\sigma^2) = \frac{1}{(2\pi)^{1/2}\sigma} \frac{1}{\Phi\Big(\frac{v_2-\mu}{\sigma}\Big) - \Phi\Big(\frac{v_1-\mu}{\sigma}\Big)} \exp\bigg(\frac{-(x-\mu)^2}{2\sigma^2}\bigg)$$

  where $\Phi(\cdot)$ is the the cumulative distribution function of a standard normal distribution.

- The log-normal distribution is parametrized with center $\mu$ and scale $\sigma^2$ as:

$$\texttt{Lognormal}(x;\mu,\sigma^2) = \frac{1}{x\sigma\sqrt{2\pi}} \exp\bigg(-\frac{(\ln x - \mu)^2}{2\sigma^2}\bigg)$$

At various points throughout this document, we use the following notation. Given $x_1, x_2, \dots, x_n \in \mathbb{R}$, we write $\tilde{\mathbb{E}}_{i=1}^n[x_i]$ and $\tilde{\mathbb{V}}_{i=1}^n[x_i]$ to denote their empirical mean and variances:

$$\tilde{\mathbb{E}}_{i=1}^n[x_i] = \frac{1}{n}\sum_{i=1}^n x_i$$

$$\tilde{\mathbb{V}}_{i=1}^n[x_i] = \frac{1}{n}\sum_{i=1}^n (x_i - \tilde{\mathbb{E}}_{i=1}^n[x_i])^2$$

### B Discretizing GLV dynamics

In this section we demonstrate how our model in Equation (2) is derived from Equation (1). Equation (1) can be written in the following form:

$$\mathrm{d}\mathbf{x}_{s,i}(t) = \mathbf{x}_{s,i}(t)[f(\mathbf{x}_s(t))\mathrm{d}t + \mathrm{d}w_{s,i}(t)].$$

where the factor $\mathbf{x}_{s,i}(t)$ has been pulled out. Dividing both sides by $\mathbf{x}_{s,i}(t)$ results in:

$$\frac{\mathrm{d}\mathbf{x}_{s,i}(t)}{\mathbf{x}_{s,i}(t)} = f(\mathbf{x}_s(t))\mathrm{d}t + \mathrm{d}w_{s,i}(t).$$

Integrating from time $t_{s,k}$ to $t_{s,k+1}$ where the lefthand side can be computed explicitly and approximating the the right hand side by the Euler-Maruyama method [11]:

$$\log \mathbf{x}_{s,i}(t_{s,k+1}) - \log \mathbf{x}_{s,i}(t_{s,k}) = f(\mathbf{x}_s(t_{s,k}))(t_{s,k+1} - t_{s,k}) + w_{s,i}(t_{s,k+1}) - w_{s,i}(t_{s,k}).$$

With $w_{s,i}(t_{s,k+1}) - w_{s,i}(t_{s,k})$ a Wiener process with variance $\Delta_{s,k}\sigma_w^2$ we have that

$$w_{s,i}(t_{s,k+1}) - w_{s,i}(t_{s,k}) \sim \mathtt{Normal}(0, \Delta_{s,k}\sigma_w^2)$$

Taken together these dynamics can be described as follows:

$$\log \mathbf{x}_{s,i}(t_{s,k+1}) \sim \mathtt{Normal}(\log \mathbf{x}_{s,i}(t_{s,k}) + f(\mathbf{x}_s(t_{s,k}))\Delta_{s,k}, \Delta_{s,k}\sigma_w^2)$$

which gives us Equation (2).

### C Stick-breaking process

Sethuraman describes a "stick-breaking" construction [18] for the DP. In this formulation, the vector of probabilities $\pi_{\mathbf{c},1}, \pi_{\mathbf{c},2}...$ is generated as:

$$\beta_j \sim \mathtt{Beta}(1, \alpha)$$
$$\pi_{\mathbf{c},k} = \beta_k \prod_{j=1}^{k-1}(1 - \beta_j)$$

By construction, $\sum_{i=1}^{\infty} \pi_{\mathbf{c},i} = 1$.

### D Bayesian regression: posterior and marginalization

In this section we review Bayesian regression and parameter marginalization with the following model:

$$y \mid w \sim \mathtt{Normal}(Xw, \Sigma_1) \quad \text{and} \quad w \sim \mathtt{Normal}(\mu_2, \Sigma_2),$$

where $X$ is an $n \times d$ matrix. We assume that $X, \mu_2, \Sigma_1, \Sigma_2$ are known. The goal of the following calculations is to compute the posterior $w \mid y$ and the marginal of $y$. Let

$$p_{y|w} \triangleq \mathtt{Normal}(y; Xw, \Sigma_1) \quad \text{and} \quad p_w \triangleq \mathtt{Normal}(w; \mu_2, \Sigma_2),$$

The joint density of $y$ and $w$ is:

$$
\begin{aligned}
p_{y,w} = p_{y|w}p_w = & \frac{1}{(2\pi)^{n/2}|\Sigma_1|^{1/2}} \frac{1}{(2\pi)^{d/2}|\Sigma_2|^{1/2}} \\
& \times \exp\left(-\frac{1}{2}(y - Xw)^\mathsf{T}\Sigma_1^{-1}(y - Xw) - \frac{1}{2}(w - \mu_2)^\mathsf{T}\Sigma_1^{-1}(w - \mu_2)\right) \\
= & \frac{1}{(2\pi)^{n/2}|\Sigma_1|^{1/2}} \frac{1}{(2\pi)^{d/2}|\Sigma_2|^{1/2}} \\
& \times \exp\left(-\frac{1}{2}(w^\mathsf{T}(X^\mathsf{T}\Sigma_1^{-1}X + \Sigma_2^{-1})w - (y^\mathsf{T}\Sigma_1^{-1}X + \mu_2^\mathsf{T}\Sigma_2^{-1})w)\right) \\
& \times \exp\left(-\frac{1}{2}(-w^\mathsf{T}(X^\mathsf{T}\Sigma_1^{-1}y + \Sigma_2^{-1}\mu_2) + \mu_2^\mathsf{T}\Sigma_2^{-1}\mu_2 + y^\mathsf{T}\Sigma_1^{-1}y)\right)
\end{aligned}
$$

To simplify the expression above, we introduce the following definitions:

$$
\begin{aligned}
\Sigma_3^{-1} &= X^\mathsf{T}\Sigma_1^{-1}X + \Sigma_2^{-1} \\
\mu_3 &= \Sigma_3(X^\mathsf{T}\Sigma_1^{-1}y + \Sigma_2^{-1}\mu_2)
\end{aligned}
$$

which results in:

$$
\begin{aligned}
p_{y|w}p_w = & \frac{1}{(2\pi)^{n/2}|\Sigma_1|^{1/2}} \frac{1}{(2\pi)^{d/2}|\Sigma_2|^{1/2}} \exp\big(-\frac{1}{2}(w^\mathsf{T}\Sigma_3^{-1}w - \mu_3^\mathsf{T}\Sigma_3^{-1}w - w^\mathsf{T}\Sigma_3^{-1}\mu_3)\big) \\
& \cdot \exp\big(-\frac{1}{2}(\mu_2^\mathsf{T}\Sigma_2^{-1}\mu_2 + y^\mathsf{T}\Sigma^{-1}y)\big).
\end{aligned}
$$

Completing the square of the quadratic term in the first exponential gives:

$$
\begin{aligned}
p_{y|w}p_w = & \frac{1}{(2\pi)^{n/2}|\Sigma_1|^{1/2}} \frac{1}{(2\pi)^{d/2}|\Sigma_2|^{1/2}} \exp\big(-\frac{1}{2}(w - \mu_3)^\mathsf{T}\Sigma_3^{-1}(w - \mu_3)\big) \\
& \cdot \exp\big(-\frac{1}{2}(-\mu_3^\mathsf{T}\Sigma_3^{-1}\mu_3 + \mu_2^\mathsf{T}\Sigma_2^{-1}\mu_2 + y^\mathsf{T}\Sigma_1^{-1}y)\big).
\end{aligned}
\tag{57}
$$

Since $p_{w|y} \propto p_{y|w}p_w$ from Equation (57) it follows that

$$
p_{w|y} \propto \exp\left(-\frac{1}{2}(w - \mu_3)^\mathsf{T}\Sigma_3^{-1}(w - \mu_3)\right).
$$

The posterior distribution for $w$ is then $w \mid y \sim \texttt{Normal}(\mu_3, \Sigma_3)$. We now discuss the marginalization of $w$ from the joint distribution in Equation (57). We begin by multiplying and dividing by $|\Sigma_3|^{1/2}$ resulting in:

$$
\begin{aligned}
p_{y|w}p_w = & \frac{1}{(2\pi)^{n/2}|\Sigma_1|^{1/2}} \frac{|\Sigma_3|^{1/2}}{|\Sigma_2|^{1/2}} \frac{1}{(2\pi)^{d/2}|\Sigma_3|^{1/2}} \exp\big(-\frac{1}{2}(w - \mu_3)^\mathsf{T}\Sigma_3^{-1}(w - \mu_3)\big) \\
& \cdot \exp\big(-\frac{1}{2}(-\mu_3^\mathsf{T}\Sigma_3^{-1}\mu_3 + \mu_2^\mathsf{T}\Sigma_2^{-1}\mu_2 + y^\mathsf{T}\Sigma_1^{-1}y)\big).
\end{aligned}
$$

Noting that

$$
\int \frac{1}{(2\pi)^{d/2}|\Sigma_3|^{1/2}} \exp\big(-\frac{1}{2}(w - \mu_3)^\mathsf{T}\Sigma_3^{-1}(w - \mu_3)\big)\mathrm{d}w
$$

we arrrive at the marginal distribution

$$
\begin{aligned}
p_y &= \int p_{y|w}p_w \, \mathrm{d}w \\
&= \frac{1}{(2\pi)^{n/2}|\Sigma_1|^{1/2}} \frac{|\Sigma_3|^{1/2}}{|\Sigma_2|^{1/2}} \exp\big(-\frac{1}{2}(\mu_2^\mathsf{T}\Sigma_2^{-1}\mu_2 + y^\mathsf{T}\Sigma_1^{-1}y - \mu_3^\mathsf{T}\Sigma_3^{-1}\mu_3)\big).
\end{aligned}
$$

### E Estimating negative binomial dispersion parameters from data

The negative binomial dispersion parameters $d_0$ and $d_1$ are estimated offline before we learn the other parameters of the model (once learned, $d_0$ and $d_1$ are fixed for the remainder of the main model inference). Inference of $d_0$ and $d_1$ is done using Metropolis-Hastings (MH) steps with a modified dynamics model. The change is that we model the latent trajectory $\mathbf{x}$ as being generated directly from the replicate data. This model is completely stand alone from the other model and so as to not cause confusion all model parameters here have an over bar (i.e. $\bar{\mathbf{x}}$ ).

#### Model details

We assume that the abundance of the latent state is generated from a normal distribution

$$\bar{\mathbf{x}}_{\kappa,i} \sim \texttt{Normal}\big(\mu_{\bar{x},\kappa,i}, 100 \cdot \sigma^2_{\bar{x},\kappa,i}\big)$$

where

$$\begin{aligned}
\bar{x}^{\text{data}}_{\kappa,\rho,i} &\triangleq \frac{\bar{y}_{\kappa,\rho,i}}{\sum_i \bar{y}_{\kappa,\rho,i}} \exp\left(\frac{1}{3}\sum_{r=1}^{3} \log \bar{Q}_{\kappa,r}\right) \\
\mu_{\bar{x},\kappa,i} &\triangleq \tilde{\mathbb{E}}^6_{\rho=1}\big[\bar{x}^{\text{data}}_{\kappa,\rho,i}\big] \\
\sigma^2_{\bar{x},\kappa,i} &\triangleq \tilde{\mathbb{V}}_i[\mu_{\bar{x},\kappa,i}]
\end{aligned}$$

Here, $\bar{y}_{\kappa,\rho,i}$ are the sequencing reads associated with taxon $i$ in sample $\kappa$ for sequencing replicate $\rho$, and $\bar{Q}_{\kappa,r}$ represents the bacterial concentration in CFU/g in sample $\kappa$ for qPCR replicate $r$. As before we assume that the reads $\mathbf{y}_{s,i}$ are negative-binomially distributed:

$$\bar{\mathbf{y}}_{\kappa,\rho,i} \mid \bar{\mathbf{x}}_\kappa, \bar{\mathbf{d}}_0, \bar{\mathbf{d}}_1 \sim \texttt{NegBin}\big(\phi(\bar{\mathbf{x}}_\kappa, \bar{r}_\kappa), \epsilon(\bar{\mathbf{x}}_\kappa, \bar{\mathbf{d}}_0, \bar{\mathbf{d}}_1)\big) \tag{58}$$

where $\phi$ and $\epsilon$ are defined in Equations (51) and (52), $\bar{r}_{\kappa,\rho}$ is the read depth for sample $\kappa$ and sequencing replicate $\rho$. We assume that $\bar{\mathbf{d}}_0$ and $\bar{\mathbf{d}}_1$ have uniform priors [3]:

$$\bar{\mathbf{d}}_0 \sim \texttt{Uniform}(0, 10^5) \tag{59}$$

$$\bar{\mathbf{d}}_1 \sim \texttt{Uniform}(0, 10^5) \tag{60}$$

We assume that the log of the qPCR measurements are log normally distributed as before:

$$\log(\bar{\mathbf{Q}}_{\kappa,r}) \sim \texttt{Normal}\left(\log\left(\sum_i \bar{\mathbf{x}}_{\kappa,i}\right), 100 \cdot \sigma^2_{\bar{\mathbf{Q}}_\kappa}\right) \tag{61}$$

where $\sigma^2_{\bar{\mathbf{Q}}_\kappa} \triangleq \tilde{\mathbb{V}}_r\big[\log \bar{Q}_{\kappa,r}\big]$ We set $\sigma^2_{\mathbf{Q}_s}$ to be the empirical variance of the triplicates (as in Section 1.7) but multiplied by 100 to make this a less informative prior.

#### Inference

For this offline procedure we learn $\mathbf{x}_{s,i}$, $\mathbf{d}_0$, and $\mathbf{d}_1$ in the following order:

1. Sample negative binomial dispersion parameters $\mathbf{d}_0$, and $\mathbf{d}_1$.
2. Sample latent trajectory $\mathbf{x}_{s,i}$.

We use MH steps with an adaptive proposal to update the values $\mathbf{d}_0$ and $\mathbf{d}_1$. Since the inference scheme is the same for both $\mathbf{d}_0$, and $\mathbf{d}_1$ we only give the details for $\mathbf{d}_0$. The proposal for the $g^{\text{th}}$ MH step is a positively truncated normal distribution centered around the previous value

$$\bar{\mathbf{d}}_0^{[*]} \sim \texttt{TruncNormal}_{(0,\infty)}(\bar{\mathbf{d}}_0^{[g-1]}, \sigma^2_{\bar{\mathbf{d}},0}) \tag{62}$$

The target distribution is proportional to Equation (58). The proposal variance $\sigma^2_{\bar{\mathbf{d}},0}$ is initialized as:

$$\sigma^2_{\bar{\mathbf{d}},0} = \frac{1}{100}\left(\bar{\mathbf{d}}_0^{[0]}\right)^2 \tag{63}$$

where $\bar{\mathbf{d}}_0^{[0]}$ is defined below. During the first half of the burn-in period, $\sigma^2_{\bar{\mathbf{d}},0}$ is tuned to adjust the acceptance rate towards 0.44, the optimal value of a scalar MH step [6] (see Appendix F). After tuning is finished, $\sigma^2_{\bar{\mathbf{d}},0}$ is fixed for the rest of inference. We initialize the negative binomial dispersion parameters as:

$$\bar{\mathbf{d}}_0^{[0]} = 0.001 \tag{64}$$

$$\bar{\mathbf{d}}_1^{[0]} = 0.05 \tag{65}$$

this is the same initialization that was used in our prior work [3], and the latent state is initialized as

$$\bar{\mathbf{x}}_{\kappa,i}^{[0]} \sim \texttt{TruncNormal}_{(0,\infty)}(\mu_{\bar{x},\kappa,i}, 10^{-4}). \tag{66}$$

### F Metropolis-Hastings Proposal Tuning

To encourage optimal mixing behavior of the Metropolis-Hastings steps we targeted an acceptance rate of 0.44 as suggested in [6]. To accomplish this, our burn-in iterations have an additional tuning-step built into the sampling step, applied once every 50 iterations. This tuning-step is applied immediately after the MH step is applied for that iteration. If the acceptance rate over the last 50 iterations is smaller than $0.44$, we encourage a more localized search by shrinking the proposal variance. If instead, the acceptance rate is larger than $0.44$, we encourage a more broad search by increasing the proposal variance. If the proposal distribution is a Normal distribution, then we shrink the variance by dividing the proposal's $\sigma^2$ by 1.5, or increase it by multiplying by 1.5. If the proposal distribution is a scaled inverse $\chi^2$ distribution then we shrink the variance by multiplying the degrees of freedom parameter $\nu$ by $1.5$, or increase the variance by dividing the degrees of freedom by $1.5$.

### G Bayes Factor Computation

In this section we let $\mathcal{D}$ to denote the observed data (qPCR measurements and sequencing read counts). With fixed module memberships the Bayes factor for an interaction from module $\ell$ to $m$ is

$$\frac{\Pr(\mathbf{z}^{(\mathbf{b})}_{\ell,m} = 1 \mid \mathcal{D}) / \Pr(\mathbf{z}^{(\mathbf{b})}_{\ell,m} = 1)}{\Pr(\mathbf{z}^{(\mathbf{b})}_{\ell,m} = 0 \mid \mathcal{D}) / \Pr(\mathbf{z}^{(\mathbf{b})}_{\ell,m} = 0)}$$

where the posteriors $\Pr(\cdot \mid \mathcal{D})$ are estimated from MCMC samples and the prior contributions are obtained by marginalizing out $\pi_{\mathbf{b}}$:

$$\begin{aligned}
\Pr(\mathbf{z}^{(\mathbf{b})}_{\ell,m} = 1) &= \frac{1}{B(b_{1,\mathbf{b}}, b_{2,\mathbf{b}})} \int_0^1 \pi_{\mathbf{b}}^{b_{1,\mathbf{b}}-1}(1-\pi_{\mathbf{b}})^{b_{2,\mathbf{b}}-1} \mathrm{d}\pi_{\mathbf{b}} \\
&= \frac{B(1+b_{1,\mathbf{b}}, b_{2,\mathbf{b}})}{B(b_{1,\mathbf{b}}, b_{2,\mathbf{b}})} \\
\Pr(\mathbf{z}^{(\mathbf{b})}_{\ell,m} = 0) &= \frac{1}{B(b_{1,\mathbf{b}}, b_{2,\mathbf{b}})} \int_0^1 (1-\pi_{\mathbf{b}})^{b_{1,\mathbf{b}}-1} \pi_{\mathbf{b}}^{b_{2,\mathbf{b}}-1} \mathrm{d}\pi_{\mathbf{b}} \\
&= \frac{B(b_{1,\mathbf{b}}, 1+b_{2,\mathbf{b}})}{B(b_{1,\mathbf{b}}, b_{2,\mathbf{b}})}
\end{aligned}$$

where $B$ is the beta function. The Bayes factors for each perturbations effect on a module is computed in the same manner.

#### References


[1] Charles E Antoniak. Mixtures of dirichlet processes with applications to bayesian nonparametric problems. *The annals of statistics*, pages 1152–1174, 1974.

[2] Stephen P Brooks and Andrew Gelman. General methods for monitoring convergence of iterative simulations. *Journal of computational and graphical statistics*, 7(4):434–455, 1998.

[3] Vanni Bucci, Belinda Tzen, Ning Li, Matt Simmons, Takeshi Tanoue, Elijah Bogart, Luxue Deng, Vladimir Yeliseyev, Mary L Delaney, Qing Liu, et al. Mdsine: Microbial dynamical systems inference engine for microbiome time-series analyses. *Genome biology*, 17(1):1–17, 2016.

[4] Richard Creswell, Jie Tan, Jonathan W Leff, Brandon Brooks, Michael A Mahowald, Ruth Thieroff-Ekerdt, and Georg K Gerber. High-resolution temporal profiling of the human gut microbiome reveals consistent and cascading alterations in response to dietary glycans. *Genome medicine*, 12(1):1–16, 2020.

[5] Michael D Escobar and Mike West. Bayesian density estimation and inference using mixtures. *Journal of the american statistical association*, 90(430):577–588, 1995.

[6] Andrew Gelman, John B Carlin, Hal S Stern, David B Dunson, Aki Vehtari, and Donald B Rubin. *Bayesian data analysis*. CRC press, 2013.

[7] Georg K Gerber, Andrew B Onderdonk, and Lynn Bry. Inferring dynamic signatures of microbes in complex host ecosystems. 2012.

[8] Travis Gibson and Georg Gerber. Robust and scalable models of microbiome dynamics. In *International Conference on Machine Learning*, pages 1763–1772, 2018.

[9] Bo S Goh. Global stability in many-species systems. *The American Naturalist*, 111(977):135–143, 1977.

[10] Charles Kemp, Joshua B Tenenbaum, Thomas L Griffiths, Takeshi Yamada, and Naonori Ueda. Learning systems of concepts with an infinite relational model. In *AAAI*, volume 3, page 5, 2006.

[11] Peter E Kloeden and Eckhard Platen. Stochastic differential equations. In *Numerical Solution of Stochastic Differential Equations*, pages 103–160. Springer, 1992.

[12] Michael I Love, Wolfgang Huber, and Simon Anders. Moderated estimation of fold change and dispersion for rna-seq data with deseq2. *Genome biology*, 15(12):1–21, 2014.

[13] Steven N MacEachern. Dependent dirichlet processes. *Technical Report, Department of Statistics, The Ohio State University*, pages 1–40, 2000.

[14] Paul J McMurdie and Susan Holmes. Waste not, want not: why rarefying microbiome data is inadmissible. *PLoS Comput Biol*, 10(4):e1003531, 2014.

[15] David Mimno, Wei Li, and Andrew McCallum. Mixtures of hierarchical topics with pachinko allocation. In *Proceedings of the 24th international conference on Machine learning*, pages 633–640, 2007.

[16] Radford M Neal. Markov chain sampling methods for dirichlet process mixture models. *Journal of computational and graphical statistics*, 9(2):249–265, 2000.

[17] Carl Edward Rasmussen. The infinite gaussian mixture model. In *Advances in neural information processing systems*, pages 554–560, 2000.

[18] Jayaram Sethuraman. A constructive definition of dirichlet priors. *Statistica sinica*, pages 639–650, 1994.

[19] Yee Whye Teh, Michael I Jordan, Matthew J Beal, and David M Blei. Hierarchical dirichlet processes. *Journal of the american statistical association*, 101(476):1566–1581, 2006.
